# Exploring differences between depression and bipolar disorder through the urinary proteome

**DOI:** 10.1101/2024.04.24.590930

**Authors:** Yuqing Liu, Zhiyu Li, Yeqing Dong, Jian Yang, Meijuan Li, Jingjing Zhou, Ying Gao, Jie Li, Gang Wang, Youhe Gao

**Affiliations:** Gene Engineering Drug and Biotechnology Beijing Key Laboratory, College of Life Sciences, Beijing Normal University, Beijing 100875, China; Beijing Anding Hospital, Capital Medical University, Beijing 100088, China; Tianjin Anding Hospital, Tianjin 300074, China

**Keywords:** urine, proteomics, depression, bipolar disorder

## Abstract

How to differentiate the diagnosis of depression and bipolar disorder has always been an important problem that needs to be solved urgently in clinical practice. In this study, from the perspective of urine proteomics, urine samples of similar age were collected from two hospitals to investigate the candidate biomarkers for differentiating the diagnosis of depression and bipolar disorder using both group analysis and one-to-many analysis(1 patient: many control samples). The experimental results of the paired group analysis showed that 108 differential proteins were identified in the depressed group compared to the bipolar group under strict screening conditions with screening criteria of FC ≥ 2 or ≤ 0.5 and a two-tailed unpaired t-test of P < 0.01, with an average of 3.7 randomly generated differential proteins, and a confidence level of 96.6 % for the correlation between these proteins and the disease difference. In the one-to-many analysis, 24 differential proteins were co-identified by the samples of 13 depressed patients, 16 of which showed a completely consistent trend of expression changes in all depressed patients studied, and 6 of which were associated with immunoglobulins; 41 differential proteins were co-identified by the samples of 12 depressed patients out of 13, and 19 of which showed a completely consistent trend of expression change in the These results reflect the strong consistency of differential proteins between the two groups of patients. 12 or more samples from depressed patients were enriched for differential proteins related to multiple biological processes and signaling pathways associated with the immune system, which is consistent with previous studies: immune mechanisms may be one of the pathogenetic mechanisms of major depression and that drugs with major immune targets can improve depressive symptoms. In the future, it may be possible to observe the immune status of patients with depression to provide direction and basis for the precise treatment of depression. The results of this paper show that urine proteomics can differentiate between depression and bipolar disorder, suggest possible mechanisms and potential targets for the treatment of depression and bipolar disorder, and provide a tool for future differential diagnosis and precision treatment of the diseases.

## 1. Introduction

### 1.1 Urine Biomarkers

Biomarkers are indicators that can reflect normal pathologic as well as physiologic processes from an objective point of view^[1]^, and in clinical practice, biomarkers are able to predict, monitor, and diagnose multifactorial diseases at different stages^[2]^. Compared to blood biomarkers, which are more widely used nowadays, the potential of urine biomarkers has not been fully developed. This is especially true for early diagnosis and state prediction of diseases. Blood proteomic changes caused by disease are metabolically excreted and cannot show significant changes in the early stages of disease due to the presence of homeostatic mechanisms regulating the internal environment in the blood. In contrast, urine, which is produced by glomerular filtration of plasma and is not regulated by homeostatic mechanisms, is sensitive to changes, and small changes in disease at an early stage can be observed in urine. And according to the results of the study, the observed changes are much earlier than in blood, earlier than in pathological sections, and even earlier than the manifestation of disease symptoms, which can be applied to the early diagnosis of diseases. For example, (1) Fanshuang Zhang et al^[3]^ from our research group found that 29 proteins changed in urine before the appearance of amyloid plaque deposition in the brain of transgenic mice with Alzheimer’s disease, and 24 of these proteins have been reported to be associated with or used as markers for Alzheimer’s disease; (2) Jianqiang Wu et al^[4]^ from our research group found in the Walker256 rat model of subcutaneous tumors that changes in urine occurred before subcutaneous tumors were touched. tumor was palpated before 10 proteins changed in the urine; (3) Zhang Y et al^[5]^ from our research group identified 15 differential proteins in the urine in a rat model of chronic pancreatitis before changes in the pathology had appeared at week 2, of which 5 had been reported to be associated with pancreatitis; (4) Ni Yanying et al^[6]^ from our research group identified 15 differential proteins in a rat model of C6 cell glioma injected into the brain before urinary proteins changed prior to the appearance of symptoms on imaging; (5) In a rat model of thioacetamide-induced hepatic fibrosis, 40 differential proteins were identified in the urine prior to the pathological changes, and 15 have been reported to be associated with fibrosis^[7]^; (6) Yin et al.^[8]^ in our research group found that prior to the elevation of blood glucose in obese type 2 diabetic rats, the urinary glucose level already showed a frequently disturbed state, which has the significance of indicating early diabetes; (7) Huang He et al^[9]^ from our research group exposed rats to smoke from conventional cigarettes and screened for biomarkers of chronic obstructive pulmonary disease (COPD), which have been reported, at only two weeks of exposure. After a comparative study, it was found that urinary proteins changed differently when tumor cells grew in different organs, such as subcutaneous^[4]^, liver^[7]^, bone ^[10]^, lungs^[11]^, and brain^[6]^, suggesting that urine has the potential to differentiate between the growth of the same tumor cells in different organs. And in terms of sample acquisition, urine is more non-invasive and easily available^[12]^, which shows that urine is a good source of biomarkers.

Currently, the detection of biomarkers in urine is receiving more and more attention from physicians and researchers, and has been applied to the treatment and research of many diseases to explore disease mechanisms, drug targets and gene functions. In our research group, Wu et al^[13]^ examined the urinary proteome of a rat model of bleomycin-induced pulmonary fibrosis and found that important biological pathways, such as the insulin-likegrowth factors-1 (IGF-1) pathway, were altered. Previously, Choi et al^[14]^ found that blocking the IGF-1 pathway with antibodies significantly prolonged the lifespan of an animal model of bleomycin-induced pulmonary fibrosis. This shows the great potential of urine proteomic studies for exploring disease mechanisms and finding drug targets. Bao Yijin et al^[15]^ from our research group established an acute hypoxia rat model, and the urinary proteomic changes showed that glutathione metabolism and hypoxic stress response were closely related, which provided clues for the prevention and treatment of plateau reaction. Meng Wenwen et al^[16]^ from our research group found that urinary proteins from stroke-prone hypertensive rats were enriched to biological pathways related to the molecular mechanisms of antihypertensive drugs, such as the signaling pathways of aldosterone and renin-angiotensin, showing the promising application of urinary proteomics for exploring disease drug targets. Identification of the urine proteome in transgenic animal models can also be used as a means to study gene function. Yuanrui Hua et al^[17]^ from our research group examined the urinary proteome of phenotype-free gene-edited animals to infer gene function from the protein level.

In pharmacology and toxicology, urine biomarkers can sensitively reflect the effects of drugs on the organism, and can be used for clinical prediction of drug efficacy, detection of drug toxicity and side effects, and provide a reference and basis for timely adjustment of therapeutic measures.Davies JC et al^[18]^ predicted the efficacy of rituximab treatment in adult systemic lupus erythematosus (SLE) patients, and the efficacy of sakubutramine valdesmokinin compared to valdesmokinin in the treatment of chronic heart failure was more effective. The group of markers found in urine by Wei Jing et al^[19]^ of our study group can be used to determine the degree of statin-related muscle damage symptoms. Bao Yijin et al^[20]^ from our research group found that the significantly changed proteins in the urine of rats after taking compound danshen drip pills were closely related to the biological pathways of glycolysis and lipid metabolism, which showed the overall effect of the traditional Chinese medicine on the organism and to a certain extent reflected the mechanism of its action in the treatment of cardiovascular diseases, which provided a new method for the study of the mechanism of action and therapeutic efficacy of traditional Chinese medicines. Zhao Chenyang et al^[21]^ from our research group compared and analyzed the urinary proteome of rats taking atorvastatin drugs produced by different manufacturers and in the same dosage form, which provided innovative methods and ideas for drug consistency evaluation. Similar studies were conducted on the liver fibrosis model obtained by thioacetamide induction^[7]^, the pulmonary fibrosis model induced by bleomycin^[12]^, the chronic pancreatitis model induced by diethyldithiocarbamate^[22]^, the renal disease model obtained by adriamycin induction^[23]^, and the urinary proteome of the chronic obstructive pulmonary disease model induced by cigarette smoke^[9]^, which were the animal models of the diseases as well as the toxicological models. Since urine can sensitively reflect the effects of drugs on the organism, the monitoring of urine proteome in animal models is a good way for pharmacological and toxicological studies^[24]^.

Urine biomarkers are not only applied in experimental animal models, but also widely used in clinical samples, which have a good indication of disease prediction, early diagnosis and disease typing. In respiratory diseases, urine proteome can provide reference for disease diagnosis and prognosis.Young BL et al^[25]^ found that urine proteome could identify tuberculosis-specific biomarkers in the urine of well-characterized active tuberculosis (TB) and non-TB patients.Denise Traxler et al^[26]^ performed urinary proteome assay on 253 COPD Patients underwent urine proteomic testing and found that urinary HSP27 levels decreased during acute exacerbations, potentially providing valuable information for the management of COPD patients hospitalized for acute exacerbations during the first year.Since 2019, the epidemic of novel coronavirus (CoVID-19) pneumonia is a threat to the health of populations worldwide. Researchers have successively found that the urinary proteome of CoVID-19 patients shows relevant changes in pathophysiological status^[27]^, which can be used to explore the pathogenesis of novel coronavirus pneumonia, and that significant immunosuppression exists in patients with early infection^[28]^; it has been found that more cytokines and their receptors have been detected in urine than in serum^[29]^, and analyzing the specific signaling pathways in the patients can search for potential therapeutic targets^[30]^. In renal diseases, the urine proteome can provide ideas about the pathogenesis of the disease.In 2020 Hao et al^[31]^ successfully identified 42 low abundance proteins and 46 high abundance proteins in urine samples from patients with chronic kidney disease (CDK). Seven KEGG pathways associated with CKD and its complications were identified from these regulatory proteins, providing new biomarkers for the pathogenesis of CDK and its complications. Urine proteome also plays a good role in the identification of benign and malignant tumors.2021 Ni et al^[32]^ performed urinary proteomic analysis of patients with benign and malignant tumors of ovarian cancer and used five of these proteins as classifiers of benign and malignant tumors. The results showed that the novel classifiers of urine proteome provide a promising noninvasive diagnostic biomarker for benign and malignant ovarian tumors.

In recent years, a number of studies have shown that urine proteomics can also observe biomarkers in neurodegenerative diseases.Mann’s research team combined proteomics technology, genetic screening and machine learning.Urine proteomics can distinguish people carrying different Parkinson’s disease-associated mutation genes with different disease manifestations, providing a promising strategy for the discovery of biomarkers for Parkinson’s disease and for the stratification of patients^[33]^. Yumi Watanabe et al.^[34]^ identified ApoC3, a potential marker for Alzheimer’s disease (AD), in the urine proteome, which may also provide clues for early diagnosis and individualized treatment of psychiatric disorders. Meng Wenwen et al^[35]^ from our research group found that a screened urine protein biomarker panel could effectively differentiate between healthy and autistic children of different age groups, which has the potential to assist in the early diagnosis and intervention of autism, and validated the results using a randomized grouping method.Wang et al^[36]^ found that the proteomes of patients with autism showed changes in biological pathways related to disease mechanisms. In our research group, Zhen Yuhang et al^[37]^ detected differentially expressed proteins in patients with major depression who had different responses to antidepressant medications, and found that urinary biomarkers have the potential to predict effective therapeutic measures in patients with major depression, which can provide clues and rationale for precise treatment and improve the quality of life of patients.

### 1.2 Depression and bipolar disorder

Depression is a mood disorder in which patients are characterized by a persistent sense of sadness as well as an inability to experience pleasure, accompanied by impairment of daily functioning. Globally, depression is the leading cause of disability and loss of productive life^[38]^. In the United States, the prevalence of depression is 5-10%, but it can be as high as 40-50% in certain primary care or specialty care settings^[39]^. Despite the existence of high-quality evidence-based therapies, only about half of patients with depression receive appropriate treatment^[40]^. Depression has a significant impact on the incidence, cost, and treatment outcomes of many common and common complications, including diabetes^[41]^, and is also a major risk factor for suicide, with suicide rates in the United States increasing by approximately 35% between 1999 and 2018^[42]^. The pathophysiologic causes of depression are unknown, and there are no clinically useful biodiagnostic markers or biological screening tests^[43]^.

Bipolar Disorder (BD), also known as manic-depressive disorder, is a chronic condition with severe debilitating symptoms that can have a profound impact on patients and their caregivers^[44]^. Bipolar disorder usually begins in adolescence or early adulthood and can have lifelong adverse effects on the patient’s physical and mental health, educational and occupational functioning, and interpersonal relationships^[45]^. Although BD is not as common as Major Depressive Disorder (MDD), the lifetime prevalence of BD in the United States is high (estimated to be approximately 4%), with similar prevalence rates across race, ethnicity, and gender^[46][47]^, and long-term outcomes have been suboptimal^[48]^. Because the manic or depressive symptoms of BD tend to be severe and recur throughout the patient’s life, the disorder can place a significant burden on patients, caregivers, and society.Episodes in patients with BD repeatedly switch between pathological mood states characterized by manic or depressive symptoms, interspersed with periods of relatively normal mood^[49]^. Formal definitions of manic and depressive symptoms are contained in the recently updated Diagnostic and Statistical Manual of Mental Disorders, Fifth Edition (DSM-5)^[50]^. Notably, in the DSM-5, depressive episodes of BD are defined by the same criteria as MDD, and thus distinguishing BD from MDD usually depends on the presence of a history of manic or hypomanic symptoms^[51]^.

Since BD usually presents with depressive symptoms early on, if depression is diagnosed and medicated at this time, it may exacerbate the patient’s condition. If clinicians are aware that a patient may have BD, they can increase the likelihood of successful identification and appropriate treatment, which can have a favorable impact on short-term outcomes and the long-term course of the disease^[48][51][52]^. The two main types of screening tools currently used for bipolar disorder are questionnaire tests and confirmatory clinical interviews, the Mood Disorder Questionnaire (MDQ) and the Composite International Diagnostic Interview, version 3.0 (CIDI 3 version 3.0, CIDI 3.0) are commonly used screening tools, with scores above specific thresholds raising suspicion of mania^[53][54]^. However, no objective physiologic indicators have been used for the differential diagnosis of bipolar disorder; therefore, in this paper, we collected urine samples from healthy individuals, patients with depression, and patients with bipolar disorder, and used urine proteomics as a basis for exploring whether biomarkers can be found in urine to differentiate between the diagnosis of depression and bipolar disorder (Fig. 1).

**Figure 1.**
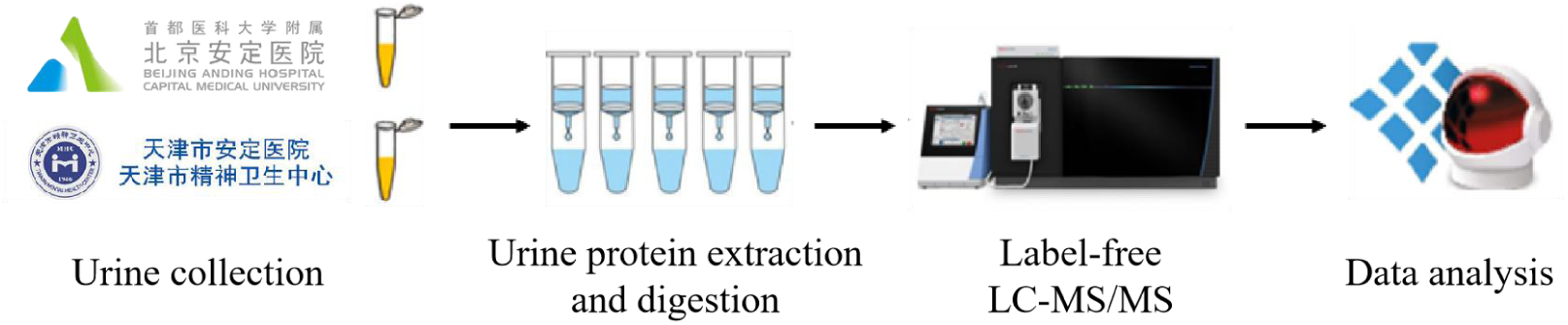
Workflow for exploring differences between depression and bipolar disorder via the urinary proteome.

## 2. Materials and Methods

### 2.1 Urine collection

The study was approved by the Human Research and Ethics Committee of Beijing Anding Hospital affiliated with Capital Medical University [(2022) Scientific Research No. (14)-202221FS-2] and by the Ethics Committee of Tianjin Anding Hospital (2019-18). All participants provided written informed consent. Participants were recruited from Beijing Anding Hospital affiliated with Capital Medical University and Tianjin Anding Hospital, and samples were collected from December 2022 to April 2023. Cases from people with schizophrenia, other psychotic disorders, personality disorders, or mental retardation diagnoses were excluded, and unmedicated samples aged between 18 and 23 were selected. The participant sample consisted of 11 samples from healthy individuals, 11 samples from individuals with bipolar disorder, and 13 samples from individuals with depression. The collected urine samples were stored in a −80°C refrigerator.

### 2.2 Treatment of the urine samples

Urine protein extraction and quantification: The collected urine samples were centrifuged at 12000×g for 30 min at 4°C, and the supernatant was transferred to a 50 mL centrifuge tube. After that, dithiothreitol solution (DTT, Sigma) was added to a final concentration of 20 mM, shaking and mixing, and then heated in a water bath at 37℃ for 1 h, and cooled to room temperature. Iodoacetamide (IAA, Sigma) was added to a final concentration of 50 mM, mixed well, and then reacted at room temperature for 40 min, avoiding light. six times the volume of pre-cooled anhydrous ethanol was added, mixed homogeneously, and then precipitated at −20°C for 24 h. On the next day, the mixed solution was centrifuged at 4°C for 30 min at 12,000×g, and the supernatant was discarded. The protein precipitate was resuspended in lysate (containing 8 mol/L urea, 2 mol/L thiourea, 25 mmol/L dithiothreitol, and 50 mmol/L Tris). The supernatant was centrifuged at 12000×g for 30 min at 4°C and placed in a new EP tube. Protein concentration was measured by Bradford method.

Uroprotein digestion: 100 μg of urinary protein sample was added to the membrane of a 10 kDa ultrafiltration tube (Pall, Port Washington, NY, USA), placed in an EP tube, and 25 mmol/L NH4HCO3 solution was added to make the total volume of 200 μL. Then the membrane was washed by: ① 200 μL of UA solution (8 mol/L urea, 0.1 mol/L Tris, 0.1 mol/L Tris, 0.1 mol/L Tris, 0.1 mol/L Tris). 0.1 mol/L Tris-HCl, pH 8.5), centrifuged at 14000×g for 5 min at 18℃ for two times; ② Sample loading: add the freshly treated sample, and centrifuged at 14000×g for 40 min at 18℃; ③ Add 200 μL of UA solution, and then centrifuged at 14000×g for 40 min at 18℃, and repeated twice; ④ Add 25 mmol/L NH4HCO3 solution to make a total volume of 200 μL, and then perform the membrane washing operation. Add 25 mmol/L NH4HCO3 solution and centrifuge at 14000×g 40 min 18°C, repeat 3-4 times; ⑤ Add trypsin (Trypsin Gold, Promega, Fitchburg, WI, USA) for digestion according to trypsin:protein ratio of 1:50 and water bath at 37°C for 12-16 h. The next day, add 200 μL of UA solution and centrifuge at 14000×g 40 min 4°C for 40 min. Peptides were collected by centrifugation at 13000 × g for 30 min at 4°C, desalted by HLB column (Waters, Milford, MA), dried using a vacuum dryer and stored at −80°C.

### 2.3 LC‒MS/MS analysis

The enzymatically digested samples were dissolved in 0.1% formic acid, and the peptides were quantified using the BCA kit, and the peptide concentration was diluted to 0.5 μg/μL. A mixed peptide sample was prepared by taking 4 μL of each sample, and the peptides were separated using the High pH Reversed-Phase Peptide Separation Kit (Thermo Fisher Scientific) according to the instructions. Ten fractions of effluent (Fractions) were collected by centrifugation, dried using a vacuum dryer and reconstituted with 0.1% formic acid. The iRT reagent (Biognosys, Switzerland) was added in a volumetric ratio of sample: iRT of 10:1 to calibrate the retention time of the extracted peptide peaks. For analysis, 1 μg of peptide from each sample was taken and analyzed by mass spectrometry using an EASY-nLC1200 chromatography system (Thermo Fisher Scientific, USA) and an Orbitrap Fusion Lumos Tribrid mass spectrometer (Thermo Fisher Scientific, USA) and data acquisition.

In order to generate a spectral library, the 10 Fractions obtained from the separation were analyzed by mass spectrometry in Data Dependent Acquisition (DDA) mode. The mass spectrometry data were acquired in high sensitivity mode. A complete mass spectral scan was acquired in the range of 350-1500 m/z with a resolution setting of 60,000.Individual samples were analyzed in Data Independent Acquisition (DIA) mode. A DIA method with 36 windows was used for DIA acquisition. A single DIA analysis of pooled peptides was performed as a quality control after every 8 samples.

### 2.4 Database searching and label-free quantitation

Raw data (RAW files), collected from LC-MS, were imported into Proteome Discoverer (version 2.1, Thermo Scientific) using the SwissProt database (taxonomy: Homo; contains 20346 sequences). Comparisons were performed and iRT sequences were added to the database. The search results were then imported into Spectronaut Pulsar (Biognosys AG, Switzerland) for processing and analysis. Peptide abundance was calculated by summing the peak areas of the respective fragment ions in MS2. Protein intensities were calculated by summing the respective peptide abundances to calculate protein abundance.

### 2.5 Statistical analysis

Three technical replicates were performed for each sample and the average was taken for statistical analysis. The identified proteins were compared to screen for differential proteins. Differential proteins were loosely screened under the following conditions: fold change between groups (FC, Fold change) ≥1.5 or ≤0. 67, and P-value <0.05 for two-tailed unpaired t-test analysis. differential proteins were strictly screened under the following conditions: fold change between groups (FC, Fold change) ≥2 or ≤0. 5, and P-value <0.01 for two-tailed unpaired t-test analysis. The screened differential proteins were analyzed using the Wukong platform (https://www.omicsolution.org/wkomic/main/), the Uniprot website (https://www.uniprot.org/) and the DAVID database (https://david.ncifcrf.gov/)) for functional enrichment analysis. And the reported literature was searched in Pubmed database (https://pubmed.ncbi.nlm.nih.gov), so as to functionally analyze the differential proteins.

## 3. Results and Discussion

### 3.1 Analysis of changes in the urinary proteome in group comparisons

#### 3.1.1 Identification and Functional Analysis of Urine Proteome

Peptides resulting from digestion of urine samples from 11 healthy individuals, 11 bipolar patients, and 13 depressed patients were subjected to LC-MS/MS tandem mass spectrometry. In total, 2612 proteins were identified (≥ 2 specific peptides and FDR < 1% at protein level).

##### 3.1.1.1 Identification and functional analysis of urine proteome in healthy and biphasic groups

Samples from the 11 healthy individuals group were compared to samples from the 11 bipolar patients group. Under relaxed conditions, the criteria for screening differential proteins were: FC ≥ 1.5 or ≤ 0.67, two-tailed unpaired t-test P < 0.05, and 67 differential proteins could be identified (Table S1). Strictly, seven differential proteins could be identified by screening criteria of FC ≥ 2 or ≤ 0.5 and a two-tailed unpaired t-test P < 0.01 (Table 2). Previously, Cannon et al^[55]^ found that brain serotonin transporter levels were elevated in the thalamus, striatum, insula, and cingulate cortex in unmedicated bipolar patients, while in 2011 X Liu et al56 conducted genome-wide association studies on this basis and concluded that non-synonymous polymorphisms in galactose mutase (GALM) are associated with the binding potential of serotonin transporters in the human thalamus. However, the results in this paper provide a strong supporting basis for the conclusions of X Liu et al.^[56]^.

**Table 1.**
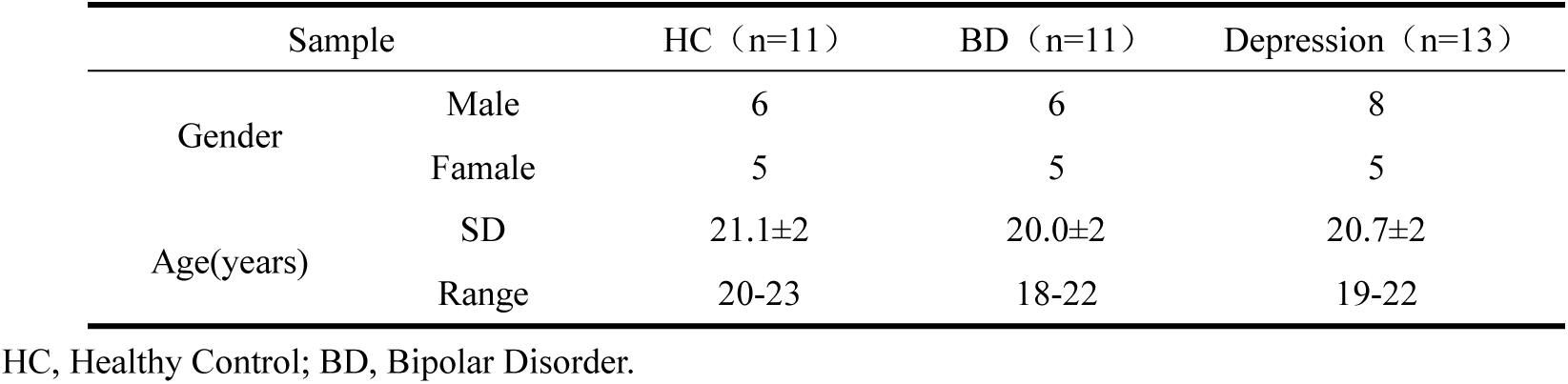
Information and statistical results of the collected samples.

**Table 2.** Differential proteins in healthy individuals compared with bipolar patients (FC ≥ 2 or ≤ 0.5, P < 0.01)

| Accession | Protein names | trend | FC | P |
| --- | --- | --- | --- | --- |
| Q96C23 | Galactose mutarotase | ↓ | 0.43 | 7.19E-04 |
| C9IZ46 | Protein shisa-5 | ↓ | 0.43 | 1.67E-03 |
| O43852 | Calumenin | ↓ | 0.47 | 4.54E-03 |
| P10451 | Osteopontin | ↑ | 2.00 | 7.75E-03 |
| O60293 | Zinc finger C3H1 domain-containing protein | ↑ | 2.36 | 8.18E-04 |
| O95433 | Activator of 90 kDa heat shock protein ATPase homolog 1 | ↑ | 2.40 | 8.47E-06 |
| P08473 | Neprilysin | ↑ | 2.55 | 1.33E-03 |

Functional analysis of the differential proteins identified in the healthy and biphasic groups under relaxed conditions enriched a total of 26 biological processes including negative regulation of endopeptidase activity, proteolysis, and cell adhesion and three signaling pathways: renin-angiotensin system (RAS), complement and coagulation cascades, and protein digestion and absorption (Table S2).Among them, Xu You et al.^[57]^ similarly enriched biological processes such as proteolysis based on potential BD diagnostic biomarkers screened, consistent with the findings presented here.In 2022, Meng-Yuan Shang et al. also enriched biological processes such as cell adhesion and calcium binding in experiments exploring the correlation between IQ levels and the potential risk of BD.Barbosa IG et al^[58]^ systematically reviewed the relationship between RAS and BD, and although Barbosa IG et al still failed to reach a firm conclusion on the relationship between the two, they summarized several studies supporting the role of RAS in central nervous system and neuropsychiatric disorders^[59–61]^and made recommendations that RAS needs to be assessed in BD patients.

##### 3.1.1.2 Identification and Functional Analysis of Urine Proteome in Healthy Group Compared with Depression Group

Samples from the 11 healthy individuals group were compared to samples from the 13 depressed patients group.Under relaxed conditions, the criteria for screening differential proteins were: FC ≥ 1.5 or ≤ 0.67, two-tailed unpaired t-test P < 0.05, and 276 differential proteins could be identified (Table S3).Under stringent conditions, the screening criteria were FC ≥ 2 or ≤ 0.5 and P < 0.01 by two-tailed unpaired t-test, and 61 differential proteins could be identified (Table 3).It has been shown that folic acid is mainly metabolized by neural tissue, 5-methyltetrahydrofolate (5-MTHF) is the main circulating form of folic acid in blood and cerebrospinal fluid, and folate receptor alpha (FR-α) is one of the main pathways mediating the transport of folic acid through biofilms, which is commonly expressed in epithelial cells, has a very high affinity for 5-MTHF, and can transport 5-MTHF from blood to epithelial cells at low folic acid levels^[63][64]^.In 2022, Nelson Siu Kei Lam et al^[62]^ systematically reviewed the potential use of folic acid and its derivatives in the treatment of psychiatric disorders and concluded that oral levorotatory folic acid or 5-methylfolic acid is associated with improved clinical efficacy in major depressive disorder.In addition, Carazo-Arias E et al.^[65]^ conducted a study on the responsiveness of fluoxetine in the treatment of depression in 2022 and showed that chronic fluoxetine treatment up-regulated the expression of proenkephalin in the dentate gyrus of mice, and this up-regulation was associated with treatment responsiveness.The encoded SDCBP gene of Mda-9/syntenin is able to activate the NF-κB pathway^[66]^, which is closely related to the occurrence and development of depression^[67–69]^, so Jiang W et al.^[70]^ concluded that SDCBP changes may control the occurrence of depression through a complex molecular regulatory network during the development of major depression in patients with rheumatoid arthritis and can be used as a potential diagnostic indicator of rheumatoid arthritis with major depression, which is mutually confirmed by the results of syntenin-2, a differential protein, screened in healthy individuals compared with depressed patients in this paper.

**Table 3.**
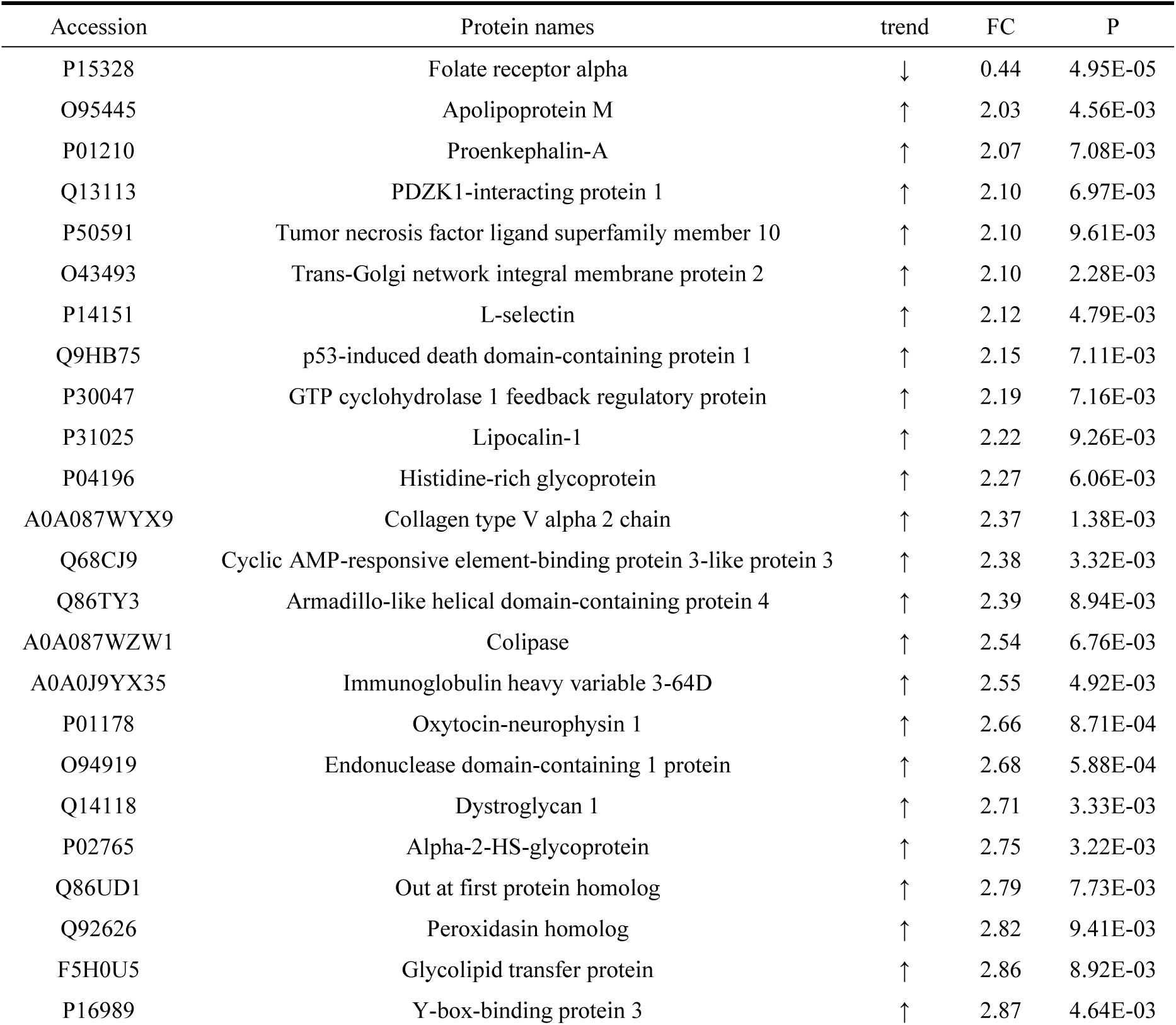
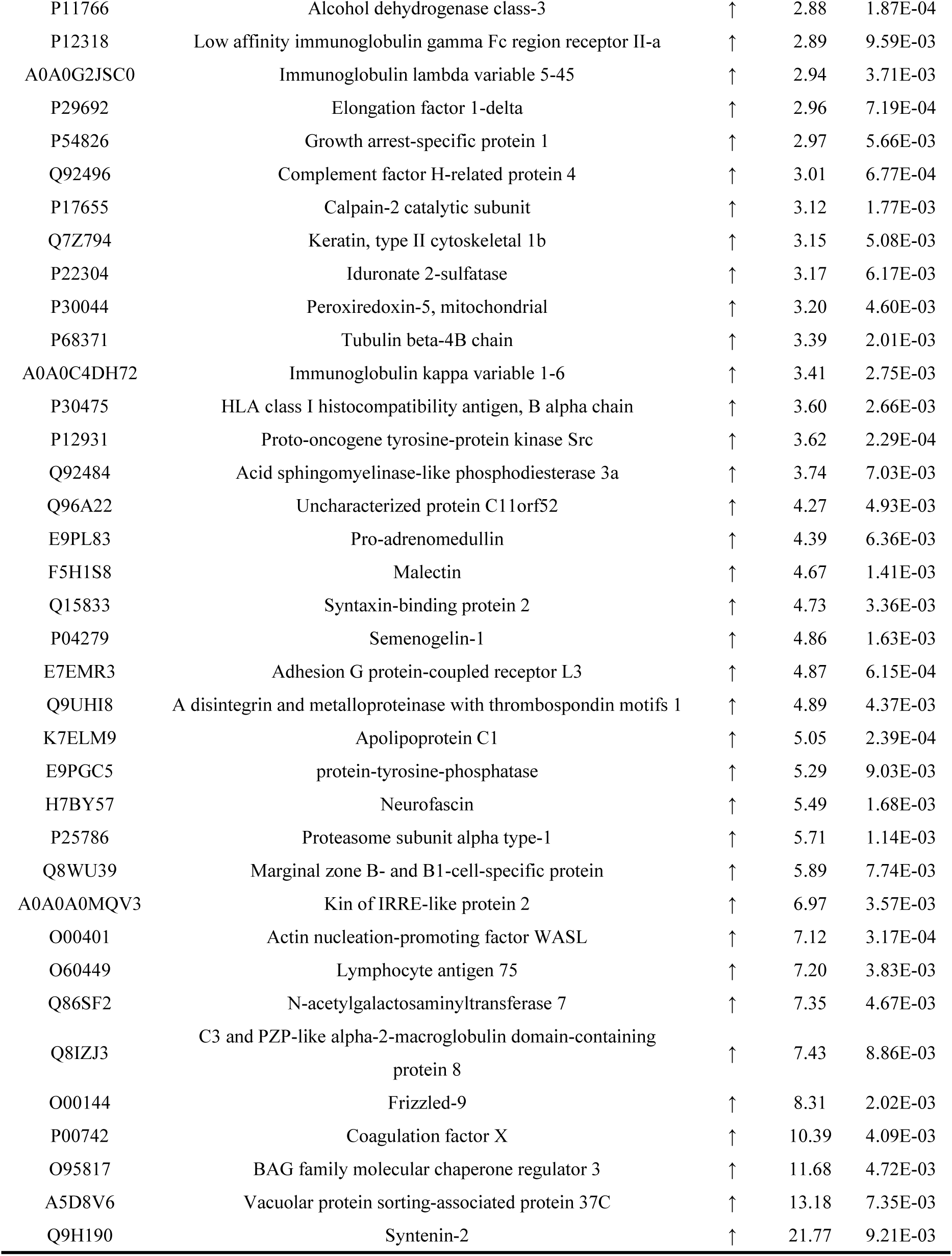
Differential proteins in healthy individuals compared with depressed patients (FC ≥ 2 or ≤ 0.5, P < 0.01)

| Accession | Protein names | trend | FC | P |
| --- | --- | --- | --- | --- |
| P15328 | Folate receptor alpha | ↓ | 0.44 | 4.95E-05 |
| O95445 | Apolipoprotein M | ↑ | 2.03 | 4.56E-03 |
| P01210 | Proenkephalin-A | ↑ | 2.07 | 7.08E-03 |
| Q13113 | PDZK1-interacting protein 1 | ↑ | 2.10 | 6.97E-03 |
| P50591 | Tumor necrosis factor ligand superfamily member 10 | ↑ | 2.10 | 9.61E-03 |
| O43493 | Trans-Golgi network integral membrane protein 2 | ↑ | 2.10 | 2.28E-03 |
| P14151 | L-selectin | ↑ | 2.12 | 4.79E-03 |
| Q9HB75 | p53-induced death domain-containing protein 1 | ↑ | 2.15 | 7.11E-03 |
| P30047 | GTP cyclohydrolase 1 feedback regulatory protein | ↑ | 2.19 | 7.16E-03 |
| P31025 | Lipocalin-1 | ↑ | 2.22 | 9.26E-03 |
| P04196 | Histidine-rich glycoprotein | ↑ | 2.27 | 6.06E-03 |
| A0A087WYX9 | Collagen type V alpha 2 chain | ↑ | 2.37 | 1.38E-03 |
| Q68CJ9 | Cyclic AMP-responsive element-binding protein 3-like protein 3 | ↑ | 2.38 | 3.32E-03 |
| Q86TY3 | Armadillo-like helical domain-containing protein 4 | ↑ | 2.39 | 8.94E-03 |
| A0A087WZW1 | Colipase | ↑ | 2.54 | 6.76E-03 |
| A0A0J9YX35 | Immunoglobulin heavy variable 3-64D | ↑ | 2.55 | 4.92E-03 |
| P01178 | Oxytocin-neurophysin 1 | ↑ | 2.66 | 8.71E-04 |
| O94919 | Endonuclease domain-containing 1 protein | ↑ | 2.68 | 5.88E-04 |
| Q14118 | Dystroglycan 1 | ↑ | 2.71 | 3.33E-03 |
| P02765 | Alpha-2-HS-glycoprotein | ↑ | 2.75 | 3.22E-03 |
| Q86UD1 | Out at first protein homolog | ↑ | 2.79 | 7.73E-03 |
| Q92626 | Peroxidasin homolog | ↑ | 2.82 | 9.41E-03 |
| F5H0U5 | Glycolipid transfer protein | ↑ | 2.86 | 8.92E-03 |
| P16989 | Y-box-binding protein 3 | ↑ | 2.87 | 4.64E-03 |
| P11766 | Alcohol dehydrogenase class-3 | ↑ | 2.88 | 1.87E-04 |
| P12318 | Low affinity immunoglobulin gamma Fc region receptor II-a | ↑ | 2.89 | 9.59E-03 |
| A0A0G2JSC0 | Immunoglobulin lambda variable 5-45 | ↑ | 2.94 | 3.71E-03 |
| P29692 | Elongation factor 1-delta | ↑ | 2.96 | 7.19E-04 |
| P54826 | Growth arrest-specific protein 1 | ↑ | 2.97 | 5.66E-03 |
| Q92496 | Complement factor H-related protein 4 | ↑ | 3.01 | 6.77E-04 |
| P17655 | Calpain-2 catalytic subunit | ↑ | 3.12 | 1.77E-03 |
| Q7Z794 | Keratin, type II cytoskeletal 1b | ↑ | 3.15 | 5.08E-03 |
| P22304 | Iduronate 2-sulfatase | ↑ | 3.17 | 6.17E-03 |
| P30044 | Peroxiredoxin-5, mitochondrial | ↑ | 3.20 | 4.60E-03 |
| P68371 | Tubulin beta-4B chain | ↑ | 3.39 | 2.01E-03 |
| A0A0C4DH72 | Immunoglobulin kappa variable 1-6 | ↑ | 3.41 | 2.75E-03 |
| P30475 | HLA class I histocompatibility antigen, B alpha chain | ↑ | 3.60 | 2.66E-03 |
| P12931 | Proto-oncogene tyrosine-protein kinase Src | ↑ | 3.62 | 2.29E-04 |
| Q92484 | Acid sphingomyelinase-like phosphodiesterase 3a | ↑ | 3.74 | 7.03E-03 |
| Q96A22 | Uncharacterized protein C11orf52 | ↑ | 4.27 | 4.93E-03 |
| E9PL83 | Pro-adrenomedullin | ↑ | 4.39 | 6.36E-03 |
| F5H1S8 | Malectin | ↑ | 4.67 | 1.41E-03 |
| Q15833 | Syntaxin-binding protein 2 | ↑ | 4.73 | 3.36E-03 |
| P04279 | Semenogelin-1 | ↑ | 4.86 | 1.63E-03 |
| E7EMR3 | Adhesion G protein-coupled receptor L3 | ↑ | 4.87 | 6.15E-04 |
| Q9UHI8 | A disintegrin and metalloproteinase with thrombospondin motifs 1 | ↑ | 4.89 | 4.37E-03 |
| K7ELM9 | Apolipoprotein C1 | ↑ | 5.05 | 2.39E-04 |
| E9PGC5 | protein-tyrosine-phosphatase | ↑ | 5.29 | 9.03E-03 |
| H7BY57 | Neurofascin | ↑ | 5.49 | 1.68E-03 |
| P25786 | Proteasome subunit alpha type-1 | ↑ | 5.71 | 1.14E-03 |
| Q8WU39 | Marginal zone B- and B1-cell-specific protein | ↑ | 5.89 | 7.74E-03 |
| A0A0A0MQV3 | Kin of IRRE-like protein 2 | ↑ | 6.97 | 3.57E-03 |
| O00401 | Actin nucleation-promoting factor WASL | ↑ | 7.12 | 3.17E-04 |
| O60449 | Lymphocyte antigen 75 | ↑ | 7.20 | 3.83E-03 |
| Q86SF2 | N-acetylgalactosaminyltransferase 7 | ↑ | 7.35 | 4.67E-03 |
| Q8IZJ3 | C3 and PZP-like alpha-2-macroglobulin domain-containing protein 8 | ↑ | 7.43 | 8.86E-03 |
| O00144 | Frizzled-9 | ↑ | 8.31 | 2.02E-03 |
| P00742 | Coagulation factor X | ↑ | 10.39 | 4.09E-03 |
| O95817 | BAG family molecular chaperone regulator 3 | ↑ | 11.68 | 4.72E-03 |
| A5D8V6 | Vacuolar protein sorting-associated protein 37C | ↑ | 13.18 | 7.35E-03 |
| Q9H190 | Syntenin-2 | ↑ | 21.77 | 9.21E-03 |

The differential proteins identified in the healthy and depressed groups under relaxed conditions were functionally analyzed and enriched into 116 biological processes such as cell adhesion, negative regulation of fibrinolysis, classical complement activation pathway, and 30 signaling pathways such as cell adhesion molecules (CAM) and lysosomes (Table S4). It is worth mentioning that the differential proteins screened in the healthy group compared with the BD group were similarly enriched in the biological process of cell adhesion. At the same time, John Jayakumar JAK et al.^[71]^ found that serotonin plays an important role in mental illness and the positive and negative effects of neuropsychiatric drugs through cell-associated adhesion, including depression, bipolar disorder, and schizophrenia.

##### 3.1.1.3 Identification and functional analysis of urine proteome in bipolar group compared with depression group

A sample of the group of 11 bipolar patients was compared to a sample of the group of 13 depressed patients. Under relaxed conditions, the criteria for screening differential proteins were: FC ≥ 1.5 or ≤ 0.67, two-tailed unpaired t-test P < 0.05, and 500 differential proteins could be identified (Table S5). Strictly, 108 differential proteins could be identified by the screening criteria of FC ≥ 2 or ≤ 0.5 with a 2-tailed unpaired t-test P < 0.01 (Table S6). Yuan Zhang et al.^[72]^ proposed that the Growth Hormone Secretagogue Receptor System is involved in the rapid and sustained antidepressant effects of paeoniflorin.

Functional analysis of the differential proteins identified in the bipolar versus depressed groups under relaxed conditions enriched 162 biological processes such as cell adhesion, classical pathway of complement activation, homophilic cell adhesion via plasma membrane adhesion molecules, and 27 signaling pathways such as complement and coagulation cascades and actin cytoskeleton regulation (Table S7). Most of these biological processes and signaling pathways are associated with the immune system, and this feature can also be reflected in the biological processes and signaling pathway results enriched by differential proteins screened from a single sample of depression compared with a healthy group.

#### 3.1.2 Unsupervised Cluster Analysis

Unsupervised cluster analysis of the whole proteins identified in the healthy and bipolar groups showed that most of the bipolar patient samples could be well opened to the healthy component (Figure 2). Unsupervised cluster analysis of the whole proteins identified in the healthy and depressed groups showed that basically the depressed patient samples could be completely separated from the healthy group, but there was still a phenomenon that some depressed patient samples were closer to the healthy human samples (Figure 3). This suggests that urine proteome can significantly distinguish healthy people from depressed patients and can provide a powerful objective indicator for the diagnosis of depression compared with traditional psychological scales. Moreover, it is demonstrated that urine proteome may have the potential to reflect the stage of depression disease, and depression samples closer to healthy individuals may have milder symptoms and be in the primary stage of depression than depression samples with greater differences from healthy individuals. When we performed unsupervised cluster analysis of the whole proteins identified in the depression group versus the bipolar group, it was found that the bipolar patient samples were more consistent with some of the depressed patient samples, while they could be well distinguished from others of the depressed patient samples (Figure 4). We speculate that a sample of depressed patients who may be more similar to the bipolar patient sample has the potential to potentially transform into bipolar disorder and may be able to provide some guidance on the patient’s future medication.

**Figure 2.**
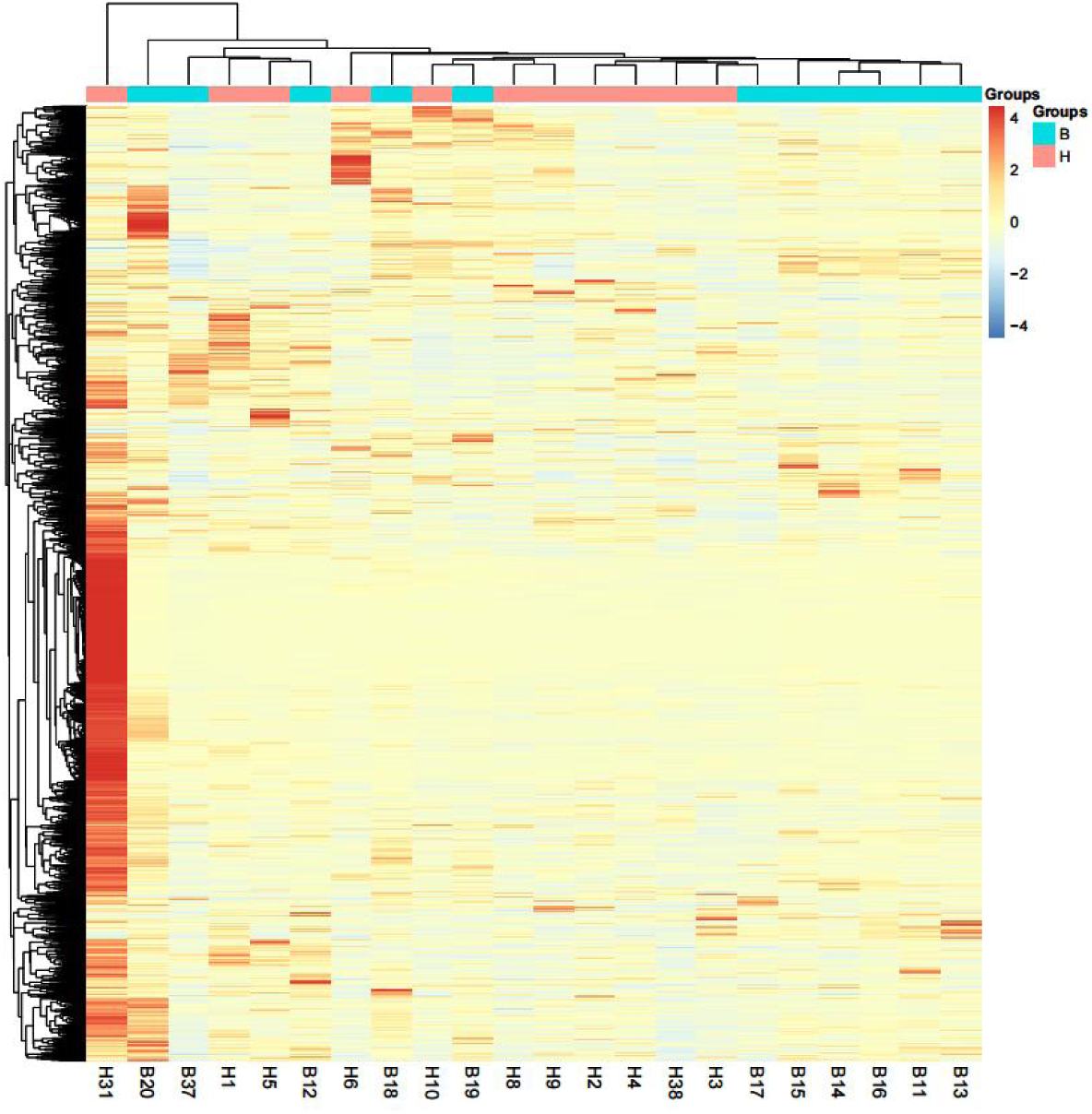
Heatmap of unsupervised clustering of whole protein in healthy versus biphasic groups.

**Figure 3.**
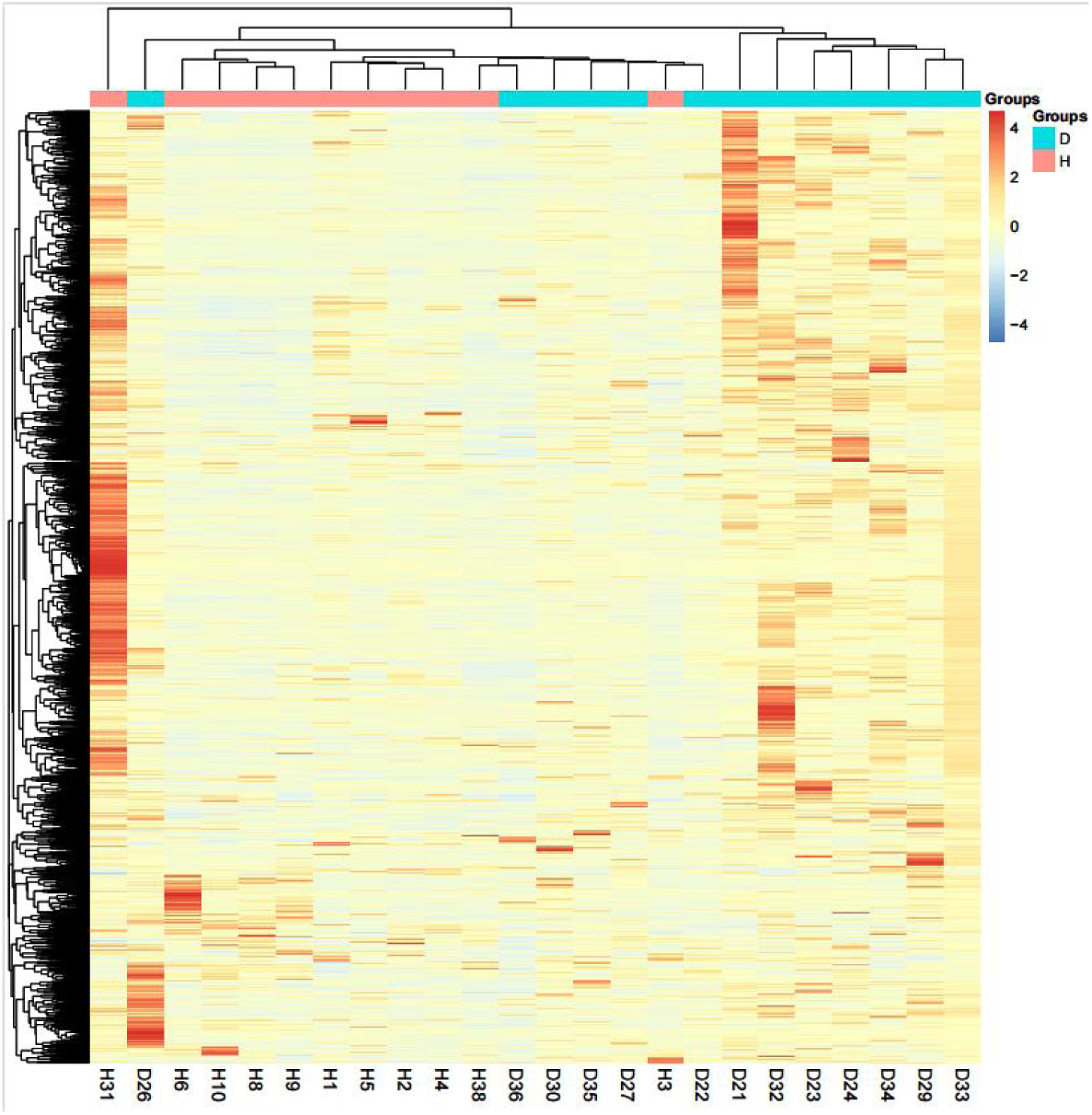
Heatmap of whole protein unsupervised clustering in healthy versus depressed groups.

**Figure 4.**
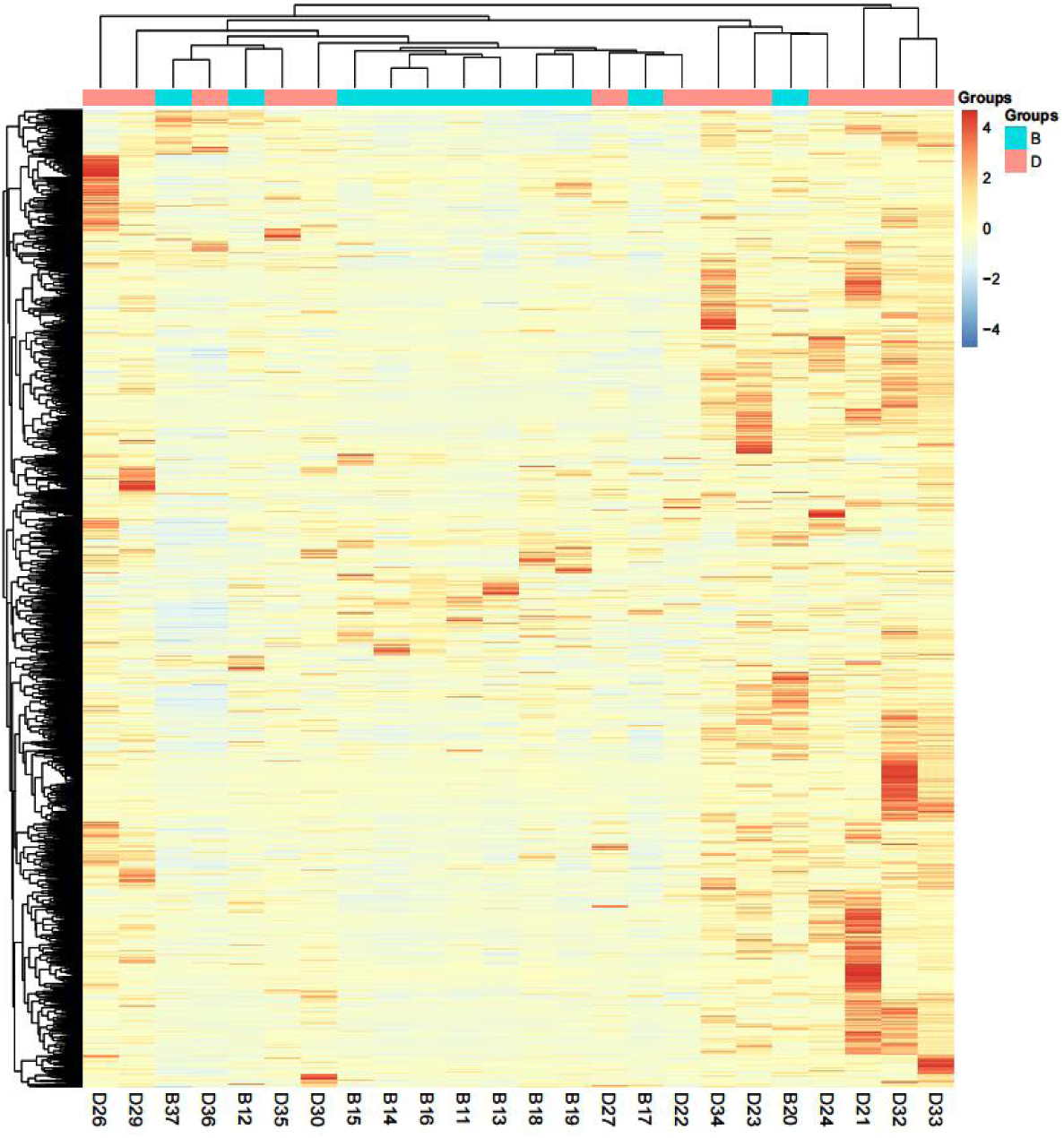
Heatmap of whole protein unsupervised clustering in bipolar versus depressed groups.

#### 3.1.3 Randomization test

In order to determine the possibility that the identified differential proteins were randomly generated, total proteins identified from 10 randomly selected healthy human samples and 10 randomly selected bipolar patient samples were randomly divided into groups for validation (FC ≥ 1.5 or ≤ 0.67, P < 0.05), resulting in an average of 41.0 differential proteins, indicating that at least 32.9% of differential proteins were not randomly generated (Table 4); random grouping validation was performed under stricter conditions (FC ≥ 2 or ≤ 0.5, P < 0.01), resulting in an average of 2.7 differential proteins, indicating that at least 60.8% of differential proteins were not randomly generated (Table 4).The total proteins identified from 10 randomly selected healthy human samples and 10 randomly selected depressed patients were randomly divided into groups for validation (FC ≥ 1.5 or ≤ 0.67, P < 0.05), resulting in an average of 54.7 differential proteins, indicating that at least 80.2% of differential proteins were not randomly generated (Table 4); randomization validation was performed under stricter conditions (FC ≥ 2 or ≤ 0.5, P < 0.01), resulting in an average of 4.4 differential proteins, indicating that at least 92.7% of differential proteins were not randomly generated (Table 4).The total proteins identified in samples from 10 randomized bipolar patients and 10 randomized depressed patients were randomized for validation (FC ≥ 1.5 or ≤ 0.67, P < 0.05), yielding an average of 53.4 differential proteins, indicating that at least 89.3% of differential proteins were not randomly generated (Table 4); randomization validation was performed under more stringent conditions (FC ≥ 2 or ≤ 0.5, P < 0.01), yielding an average of 3.7 differential proteins, indicating that at least 96.6% of differential proteins were not randomly generated (Table 4).

**Table 4.** Randomization results.

| Group | Screening criteria | Total number of random combinations | Average numbers of proteins with false random combinations | Ratio (average number of protein false random combinations/number of correctly identified differential proteins) |
| --- | --- | --- | --- | --- |
| HC/BD | FC≥1.5 or ≤0.67; P<0.05 | 67 | 41.0 | 32.9 % |
| | FC≥2 or ≤0.5; P<0.01 | 7 | 2.7 | 60.8 % |
| HC/D | FC≥1.5 or ≤0.67; P<0.05 | 276 | 54.7 | 80.2 % |
| | FC≥2 or ≤0.5; P<0.01 | 61 | 4.4 | 92.7 % |
| BD/D | FC≥1.5 or ≤0.67; P<0.05 | 500 | 53.4 | 89.3 % |
| | FC≥2 or ≤0.5; P<0.01 | 108 | 3.7 | 96.6 % |

By comparing the proportion of randomly generated differential proteins in different groups, it can be seen that the difference between healthy human samples and bipolar patient samples is less significant, the difference between healthy human samples and depressed patient samples is greater significant, while the difference between bipolar patient samples and depressed patient samples is greater significant and has the highest reliability.

### 3.2 Analysis of changes in one-to-many urine proteome

In order to better determine the disease condition of each patient in the bipolar and depressive groups and achieve the effect of precise treatment, we compared and analyzed a single patient sample with 11 samples in the healthy group for one more pair.

#### 3.2.1 Identification of Urine Proteome

Eleven samples from the biphasic group were compared with the healthy group, and the selected differential proteins and their expression trends under relaxed conditions are shown in Table 5. Thirteen samples from the depressed group were compared with the healthy group, and the selected differential proteins and their expression trends under relaxed conditions are shown in Table 5. By comparing the results of the two groups, we found that the number of differential proteins screened from a single depressed patient sample was generally greater than that from a single bipolar patient sample compared with a healthy group, indicating that depressed patients showed quite significant differences from healthy individuals, a phenomenon consistent with the results of unsupervised cluster analysis in group analysis.

**Table 5.** Expression changes of differential proteins in single samples of bipolar group and single samples of depression group compared with healthy group (FC ≥ 1.5 or ≤ 0.67, P < 0.05)

| Group | Trend | ID |  |  |  |  |  |  |  |  |  |  |  |  |
| --- | --- | --- | --- | --- | --- | --- | --- | --- | --- | --- | --- | --- | --- | --- |
|  |  | 1 | 2 | 3 | 4 | 5 | 6 | 7 | 8 | 9 | 10 | 11 | 12 | 13 |
| HC/BD | Total | 122 | 146 | 173 | 155 | 154 | 120 | 167 | 192 | 160 | 274 | 252 | - | - |
|  | ↑ | 78 | 92 | 92 | 82 | 98 | 74 | 67 | 143 | 97 | 225 | 98 | - | - |
|  | ↓ | 44 | 54 | 81 | 73 | 56 | 46 | 100 | 49 | 63 | 49 | 154 | - | - |
| HC/D | Total | 813 | 670 | 698 | 678 | 706 | 685 | 554 | 675 | 749 | 684 | 682 | 676 | 660 |
|  | ↑ | 674 | 413 | 561 | 521 | 550 | 383 | 375 | 420 | 554 | 437 | 473 | 387 | 334 |
|  | ↓ | 139 | 257 | 137 | 157 | 156 | 302 | 179 | 255 | 195 | 247 | 209 | 289 | 326 |

#### 3.2.2 Analysis of common differential proteins in single samples from the bipolar group and single samples from the depressed group compared with the healthy group

In order to investigate whether there were common differential proteins between the 11 bipolar patients, Venn diagram analysis of differential proteins screened under relaxed conditions in a single sample of the bipolar group compared with the healthy group revealed that 4 differential proteins were co-screened by 8 of the 11 bipolar patients and showed a more consistent trend of expression; 9 differential proteins were co-screened by 7 bipolar patients (Table 6). Yachen Shi et al.^[73]^ identified significantly elevated levels of antithrombin III (AT III) expression in plasma of patients with major depression and were listed as candidate biomarkers. In 2021, Cem Cerit et al^[74]^ found that the intensity of AMBP protein discriminated manic episodes in BD patients from remitted and healthy controls, indicating that AMBP protein may be a candidate biomarker for manic episodes in BD patients.

**Table 6.**
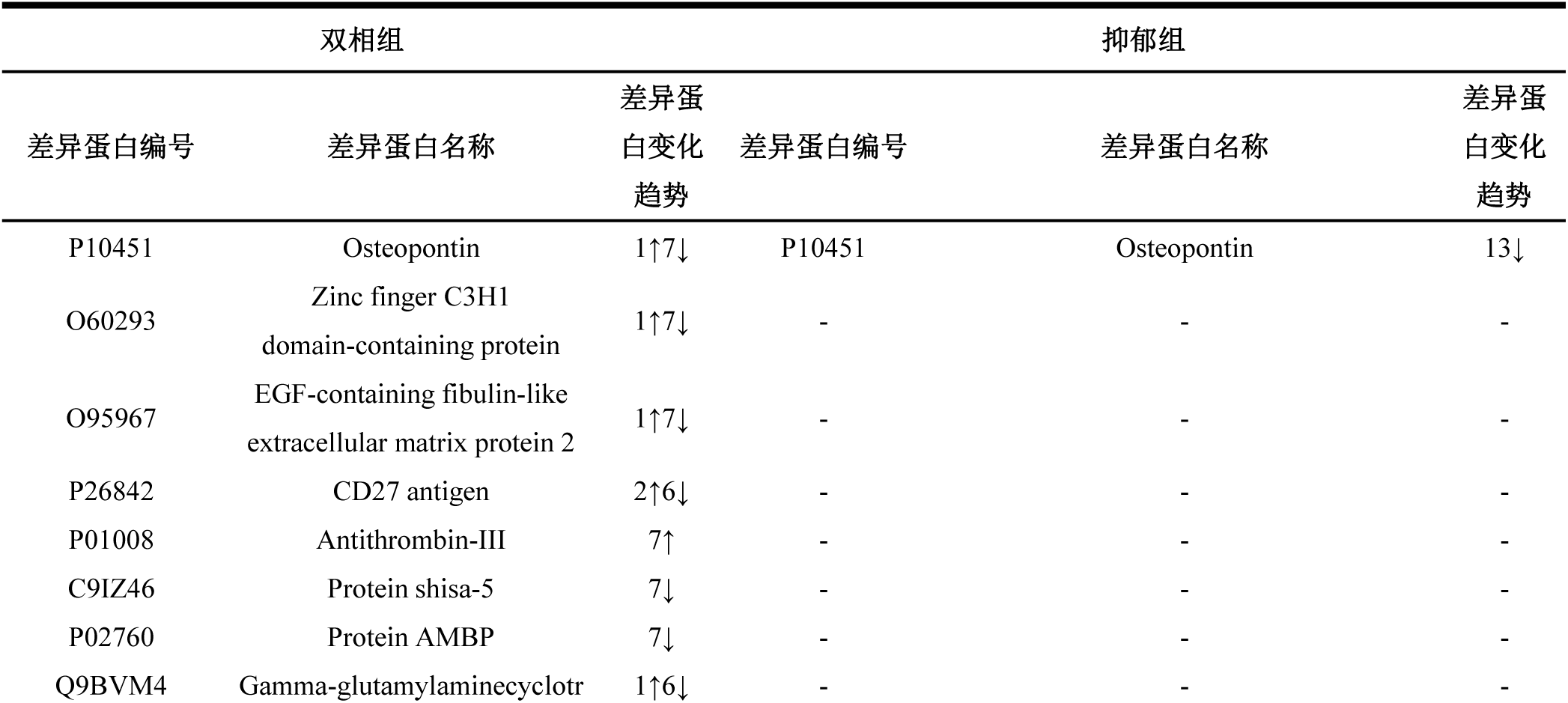
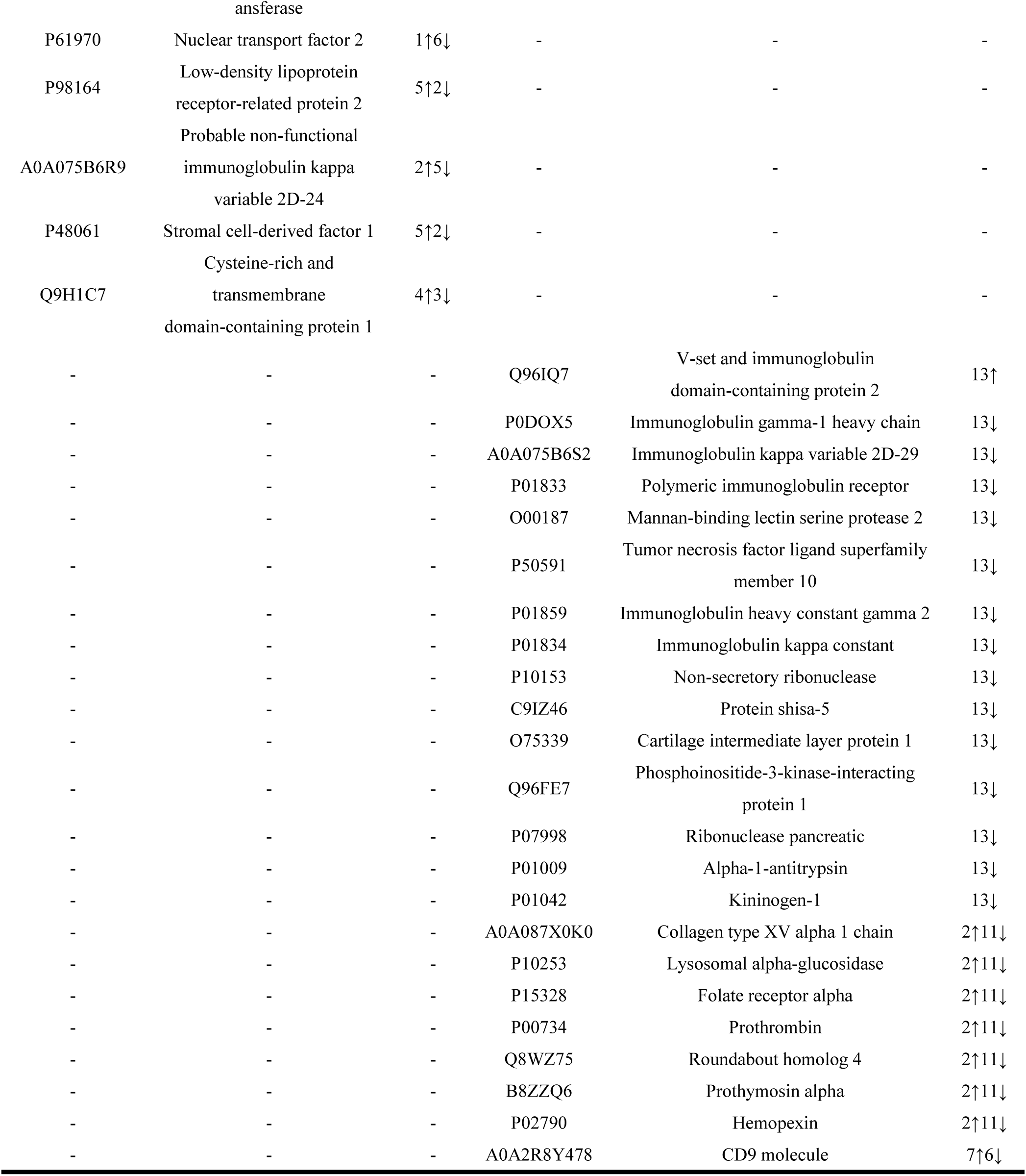
Common differential proteins in single samples from the bipolar group and single samples from the depressed group compared with the healthy group (FC ≥ 1.5 or ≤ 0.67, P < 0.05).

Venn diagram analysis of differential proteins screened under relaxed conditions in a single sample of the depression group compared with the healthy group revealed that 24 differential proteins were co-screened by 13 samples of depression patients, of which 16 differential proteins showed a completely consistent trend of expression in 13 depression patients (Table 6), and 6 differential proteins were related to immunoglobulins (immunoglobulin); 41 differential proteins were co-screened by 12 of 13 depression patient samples(Table S8), and 19 differential proteins showed a completely consistent trend of expression in 12 depression patients, and these results reflected extremely strong consistency.It is worth mentioning that osteopontin (OPN), one of the key molecules involved in neuroinflammation^[75]^, was screened in both bipolar and depressed groups, and Tingting Li et al.^[76]^ showed that blocking osteopontin expression reduced neuroinflammation and alleviated osteopontin depression-like behavior induced by lipopolysaccharide in mice.Kadriu, B. et al.^[77]^ found that OPN levels in plasma were significantly lower in patients with major depression.Lee H et al.^[78]^ screened biomarkers in plasma that were significantly associated with the severity of depressive symptoms (e.g., anhedonia/retardation): Mannan-binding lectin serine protease 2 (MASP2), and were similarly enriched in the complement activation pathway, which matched the results presented here.Members of the tumor necrosis factor (TNF) superfamily play a role in the pathogenesis of neuropsychiatric disorders because of their relationship with inflammation and neurogenesis, and increases in pro-inflammatory factors such as TNF may be typical of low-grade inflammation in major depressive disorder^[79]^.

In order to further investigate the consistent differences between individual samples from the depression group and the healthy group, we performed functional analysis of 65 differential proteins co-screened from 12 and more depression patient samples, enriched a total of 25 biological processes such as the classical pathway of complement activation, retinal homeostasis, adaptive immune response, and cell adhesion, most of which were associated with the immune system (Table 7), while we also enriched the signaling pathway of complement and coagulation cascade (P < 0.05). Immune mechanisms may be one of the pathogenesis of major depression, while drugs with major immune targets can improve depressive symptoms^[80]^. The results of this paper undoubtedly confirm the correlation between depression and the immune system, and may provide direction and basis for the precise treatment of depression by observing the immune status of depressed patients. This result also reflects that the urinary proteome can propose possible mechanisms and potential targets for the treatment of depression and bipolar disorder, providing tools for the differential diagnosis and precise treatment of diseases in the future.

**Table 7.**
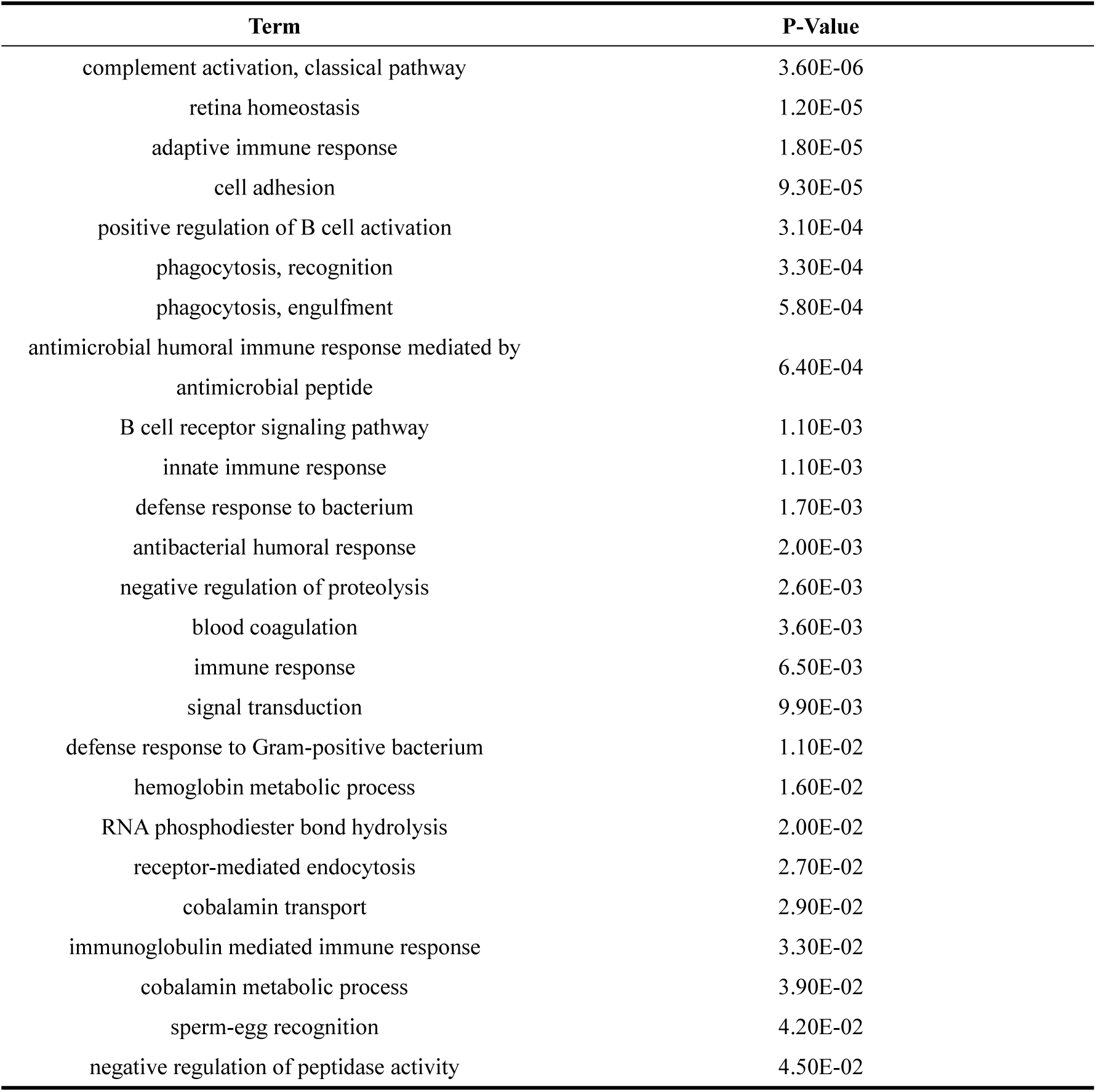
Biological processes enriched by common differential proteins in 12 or more samples from the depression group compared with the healthy group (P < 0.05)

## 4. Conclusion

In this study, we found differences in the urinary proteome between depression and bipolar disorder, confirming the association between depression and the immune system, a result consistent with previous studies: immune mechanisms may be one of the pathogenesis of major depression, while drugs with major immune targets can improve depressive symptoms. In the future, it may be possible to provide direction and basis for precise treatment of depression by observing the immune status of depression patients. The results of this study showed that urine proteome can distinguish the diagnosis of depression and bipolar disorder, and put forward possible mechanisms and potential targets for the treatment of depression and bipolar disorder, providing a tool for the differential diagnosis and precise treatment of diseases in the future.

