## Supplemental table1-8 for "Exploring differences between depression and bipolar disorder through the urinary proteome"

TableS1 Differential proteins in healthy individuals compared to bipolar patients (FC≥1.5 or ≤0.67, P<0.05)

| **Accession** | **Protein names** | **Trend** | **FC** | **P-Value** |
| --- | --- | --- | --- | --- |
| A0A0C4ZNX3 | Killer cell immunoglobulin-like receptor | ↓ | 0.28 | 2.92E-02 |
| O15511 | Actin-related protein 2/3 complex subunit 5 | ↓ | 0.37 | 3.32E-02 |
| Q96C23 | Galactose mutarotase | ↓ | 0.43 | 7.19E-04 |
| C9IZ46 | Protein shisa-5 | ↓ | 0.43 | 1.67E-03 |
| O43852 | Calumenin | ↓ | 0.47 | 4.54E-03 |
| P14174 | Macrophage migration inhibitory factor | ↓ | 0.48 | 1.24E-02 |
| O95388 | CCN family member 4 | ↓ | 0.49 | 3.98E-02 |
| P15121 | Aldo-keto reductase family 1 member B1 | ↓ | 0.50 | 4.58E-02 |
| P32942 | Intercellular adhesion molecule 3 | ↓ | 0.51 | 1.37E-03 |
| O95967 | EGF-containing fibulin-like extracellular matrix protein 2 | ↓ | 0.51 | 1.81E-04 |
| P21695 | Glycerol-3-phosphate dehydrogenase | ↓ | 0.53 | 4.89E-02 |
| Q8N386 | Leucine-rich repeat-containing protein 25 | ↓ | 0.54 | 9.40E-03 |
| P00918 | Carbonic anhydrase 2 | ↓ | 0.56 | 2.09E-02 |
| P83110 | Serine protease HTRA3 | ↓ | 0.57 | 3.84E-02 |
| A6NI73 | Leukocyte immunoglobulin-like receptor subfamily A member 5 | ↓ | 0.58 | 1.90E-03 |
| O95297 | Myelin protein zero-like protein 1 | ↓ | 0.62 | 2.48E-02 |
| P35555 | Fibrillin-1 [Cleaved into: Asprosin] | ↓ | 0.62 | 3.20E-02 |
| Q9BVM4 | Gamma-glutamylaminecyclotransferase | ↓ | 0.64 | 1.28E-02 |
| A0A087X0K0 | Collagen type XV alpha 1 chain | ↓ | 0.64 | 2.82E-02 |
| O60704 | Protein-tyrosine sulfotransferase 2 | ↓ | 0.64 | 2.60E-02 |
| O94760 | Dimethylarginine dimethylaminohydrolase 1 | ↓ | 0.64 | 7.44E-03 |
| P78417 | Glutathione S-transferase omega-1 | ↓ | 0.65 | 3.38E-02 |
| P36955 | Pigment epithelium-derived factor | ↓ | 0.65 | 2.37E-02 |
| Q9Y5E4 | Protocadherin beta-5 | ↓ | 0.65 | 5.03E-03 |
| P15328 | Folate receptor alpha | ↓ | 0.66 | 5.69E-03 |
| P29279 | CCN family member 2 | ↓ | 0.67 | 2.04E-02 |
| P08582 | Melanotransferrin | ↓ | 0.67 | 2.77E-02 |
| Q86UD1 | Out at first protein homolog | ↑ | 1.50 | 3.72E-02 |
| Q6EMK4 | Vasorin | ↑ | 1.52 | 4.06E-02 |
| Q9H3G5 | Probable serine carboxypeptidase CPVL | ↑ | 1.52 | 4.57E-02 |
| P30047 | GTP cyclohydrolase 1 feedback regulatory protein | ↑ | 1.52 | 4.10E-02 |
| A8MTF8 | FAM3 metabolism regulating signaling molecule B | ↑ | 1.52 | 2.14E-02 |
| Q8IV08 | 5'-3' exonuclease PLD3 | ↑ | 1.55 | 4.71E-02 |
| Q9H1C7 | Cysteine-rich and transmembrane domain-containing protein 1 | ↑ | 1.56 | 3.56E-02 |
| P00338 | L-lactate dehydrogenase A chain | ↑ | 1.57 | 4.76E-02 |
| P01008 | Antithrombin-III | ↑ | 1.58 | 1.04E-02 |
| E9PHN6 | Glutathione S-transferase | ↑ | 1.58 | 4.46E-02 |
| O43493 | Trans-Golgi network integral membrane protein 2 | ↑ | 1.59 | 2.60E-02 |
| P25774 | Cathepsin S | ↑ | 1.60 | 4.19E-02 |
| Q92743 | Serine protease HTRA1 | ↑ | 1.60 | 2.95E-02 |
| P10619 | Lysosomal protective protein | ↑ | 1.61 | 4.08E-02 |
| O43505 | Beta-1,4-glucuronyltransferase 1 | ↑ | 1.64 | 8.96E-03 |
| P48061 | Stromal cell-derived factor 1 | ↑ | 1.65 | 1.17E-02 |
| O43895 | Xaa-Pro aminopeptidase 2 | ↑ | 1.65 | 3.23E-02 |
| O95445 | Apolipoprotein M | ↑ | 1.68 | 1.15E-03 |
| P01009 | Alpha-1-antitrypsin | ↑ | 1.68 | 2.92E-02 |
| Q08380 | Galectin-3-binding protein | ↑ | 1.69 | 3.35E-02 |
| Q14314 | Fibroleukin | ↑ | 1.71 | 4.73E-02 |
| E9PMR4 | Tetraspanin | ↑ | 1.72 | 3.37E-02 |
| P01019 | Angiotensinogen | ↑ | 1.75 | 4.72E-03 |
| P12821 | Angiotensin-converting enzyme | ↑ | 1.77 | 2.23E-02 |
| P0C7U0 | Protein ELFN1 | ↑ | 1.78 | 1.64E-02 |
| C9JXF9 | Insulin-like growth factor-binding protein 1 | ↑ | 1.81 | 2.60E-02 |
| Q9UBD6 | Ammonium transporter Rh type C | ↑ | 1.84 | 3.42E-02 |
| Q1EHB4 | Sodium-coupled monocarboxylate transporter 2 | ↑ | 1.90 | 1.93E-02 |
| A0A087WYX9 | Collagen type V alpha 2 chain | ↑ | 1.90 | 5.47E-03 |
| P49006 | MARCKS-related protein | ↑ | 1.93 | 1.83E-02 |
| P12724 | Eosinophil cationic protein | ↑ | 1.99 | 3.68E-02 |
| P10451 | Osteopontin | ↑ | 2.00 | 7.75E-03 |
| Q92820 | Gamma-glutamyl hydrolase | ↑ | 2.14 | 2.14E-02 |
| P05154 | Plasma serine protease inhibitor | ↑ | 2.14 | 1.43E-02 |
| P01024 | Complement C3 | ↑ | 2.16 | 2.11E-02 |
| E7EMR3 | Adhesion G protein-coupled receptor L3 | ↑ | 2.33 | 1.38E-02 |
| O60293 | Zinc finger C3H1 domain-containing protein | ↑ | 2.36 | 8.18E-04 |
| O95433 | Activator of 90 kDa heat shock protein ATPase homolog 1 | ↑ | 2.40 | 8.47E-06 |
| O00144 | Frizzled-9 | ↑ | 2.42 | 2.11E-02 |
| P08473 | Neprilysin | ↑ | 2.55 | 1.33E-03 |

TableS2 Biological Processes and Signaling Pathways Enriched for Differential Proteins Produced in the Healthy Group Compared to the Biphasic Group Under Relaxed Conditions(P<0.05)

| **BP** | **P-Value** | **Pathway** | **P-Value** |
| --- | --- | --- | --- |
| negative regulation of endopeptidase activity | 3.10E-05 | Renin-angiotensin system | 1.50E-04 |
| proteolysis | 5.60E-05 | Complement and coagulation cascades | 7.10E-03 |
| cell adhesion | 7.00E-05 | Protein digestion and absorption | 1.20E-02 |
| aging | 1.00E-04 | - | - |
| ossification | 2.30E-03 | - | - |
| positive regulation of protein tyrosine kinase activity | 4.10E-03 | - | - |
| positive regulation of neurogenesis | 4.70E-03 | - | - |
| positive regulation of inflammatory response | 5.10E-03 | - | - |
| regulation of renal output by angiotensin | 6.40E-03 | - | - |
| kidney development | 6.40E-03 | - | - |
| signal transduction | 7.10E-03 | - | - |
| substance P catabolic process | 9.60E-03 | - | - |
| hormone catabolic process | 1.30E-02 | - | - |
| positive regulation of cell activation | 1.30E-02 | - | - |
| negative regulation of gene expression | 1.70E-02 | - | - |
| regulation of systemic arterial blood pressure by renin-angiotensin | 1.90E-02 | - | - |
| bradykinin catabolic process | 2.20E-02 | - | - |
| cellular oxidant detoxification | 2.30E-02 | - | - |
| positive regulation of ryanodine-sensitive calcium-release channel activity | 2.80E-02 | - | - |
| negative regulation of ryanodine-sensitive calcium-release channel activity | 3.50E-02 | - | - |
| angiotensin-activated signaling pathway | 3.80E-02 | - | - |
| beta-amyloid clearance | 4.10E-02 | - | - |
| beta-amyloid metabolic process | 4.10E-02 | - | - |
| negative regulation of transforming growth factor beta receptor signaling pathway | 4.10E-02 | - | - |
| response to acidic pH | 4.40E-02 | - | - |
| negative regulation of defense response to virus | 4.70E-02 | - | - |

TableS3 Differential Proteins in Healthy Individuals Compared to Depressed Patients (FC≥1.5 or ≤0.67, P<0.05)

| **Accession** | **Protein names** | **Trend** | **FC** | **P-Value** |
| --- | --- | --- | --- | --- |
| A0A182DWH7 | Selenoprotein P | ↓ | 0.44 | 1.36E-02 |
| P15328 | Folate receptor alpha | ↓ | 0.44 | 4.95E-05 |
| P68104 | Elongation factor 1-alpha 1 | ↓ | 0.48 | 1.64E-02 |
| A0A2R8Y5S7 | Radixin | ↓ | 0.55 | 1.52E-03 |
| Q8WUM4 | Programmed cell death 6-interacting protein | ↓ | 0.57 | 4.98E-03 |
| C9IZ46 | Protein shisa-5 | ↓ | 0.58 | 1.14E-02 |
| P14618 | Pyruvate kinase PKM | ↓ | 0.59 | 2.67E-02 |
| P26038 | Moesin | ↓ | 0.60 | 4.89E-02 |
| Q6PK18 | 2-oxoglutarate and iron-dependent oxygenase domain-containing protein 3 | ↓ | 0.60 | 1.28E-02 |
| F8VNT9 | CD63 molecule | ↓ | 0.62 | 1.04E-02 |
| Q96C23 | Galactose mutarotase | ↓ | 0.62 | 8.13E-03 |
| A0A087X0K0 | Collagen type XV alpha 1 chain | ↓ | 0.63 | 1.96E-02 |
| Q9UNZ2 | NSFL1 cofactor p47 | ↓ | 0.65 | 4.68E-02 |
| P61970 | Nuclear transport factor 2 | ↓ | 0.65 | 4.03E-03 |
| P00734 | Prothrombin | ↓ | 0.65 | 3.06E-02 |
| A6NI73 | Leukocyte immunoglobulin-like receptor subfamily A member 5 | ↓ | 0.66 | 5.40E-03 |
| O00115 | Deoxyribonuclease-2-alpha | ↓ | 0.67 | 4.95E-02 |
| P18206 | Vinculin | ↑ | 1.50 | 4.02E-02 |
| P07204 | Thrombomodulin | ↑ | 1.50 | 8.00E-04 |
| A0A3B3ISV3 | Collagen type IV alpha 1 chain | ↑ | 1.51 | 1.26E-02 |
| A0A075B6R2 | Immunoglobulin heavy variable 4-4 | ↑ | 1.52 | 2.69E-02 |
| P28799 | Progranulin | ↑ | 1.52 | 1.98E-02 |
| P24593 | Insulin-like growth factor-binding protein 5 | ↑ | 1.53 | 2.74E-02 |
| P05362 | Intercellular adhesion molecule 1 | ↑ | 1.53 | 2.76E-02 |
| Q14376 | UDP-glucose 4-epimerase | ↑ | 1.53 | 2.52E-02 |
| P51148 | Ras-related protein Rab-5C | ↑ | 1.54 | 4.54E-02 |
| Q68D85 | Natural cytotoxicity triggering receptor 3 ligand 1 | ↑ | 1.54 | 2.16E-02 |
| Q92692 | Nectin-2 | ↑ | 1.54 | 9.85E-03 |
| A0A1B0GU58 | Propionyl-CoA carboxylase alpha chain, mitochondrial | ↑ | 1.54 | 2.83E-02 |
| P0DOX5 | Immunoglobulin gamma-1 heavy chain | ↑ | 1.54 | 1.98E-02 |
| E5RIW3 | Tubulin-specific chaperone A | ↑ | 1.55 | 3.68E-02 |
| P07711 | Procathepsin L | ↑ | 1.55 | 1.69E-02 |
| P39060 | Collagen alpha-1 | ↑ | 1.56 | 2.60E-02 |
| P78324 | Tyrosine-protein phosphatase non-receptor type substrate 1 | ↑ | 1.57 | 4.09E-02 |
| A0A0A0MS15 | Immunoglobulin heavy variable 3-49 | ↑ | 1.58 | 3.27E-02 |
| J3QQX6 | Intercellular adhesion molecule 2 | ↑ | 1.59 | 2.10E-03 |
| Q15907 | Ras-related protein Rab-11B | ↑ | 1.59 | 7.48E-03 |
| P55957 | BH3-interacting domain death agonist | ↑ | 1.59 | 1.44E-02 |
| E7ES19 | Thrombospondin 4 | ↑ | 1.60 | 3.70E-02 |
| P11362 | Fibroblast growth factor receptor 1 | ↑ | 1.62 | 1.85E-02 |
| A0A087X0T8 | Cell adhesion molecule 1 | ↑ | 1.64 | 2.59E-03 |
| P08571 | Monocyte differentiation antigen CD14 | ↑ | 1.65 | 4.61E-02 |
| H3BP20 | Beta-hexosaminidase | ↑ | 1.65 | 2.26E-02 |
| A8MTF8 | FAM3 metabolism regulating signaling molecule B | ↑ | 1.66 | 2.47E-02 |
| P02760 | Protein AMBP | ↑ | 1.66 | 4.78E-02 |
| P13671 | Complement component C6 | ↑ | 1.66 | 1.81E-02 |
| B8ZZ73 | Interleukin 1 receptor type 1 | ↑ | 1.67 | 2.51E-02 |
| P15169 | Carboxypeptidase N catalytic chain | ↑ | 1.68 | 4.97E-03 |
| Q6UX71 | Plexin domain-containing protein 2 | ↑ | 1.68 | 2.00E-02 |
| Q496F6 | CMRF35-like molecule 2 | ↑ | 1.68 | 2.94E-02 |
| P25325 | 3-mercaptopyruvate sulfurtransferase | ↑ | 1.69 | 4.70E-02 |
| O43505 | Beta-1,4-glucuronyltransferase 1 | ↑ | 1.70 | 9.66E-03 |
| P27797 | Calreticulin | ↑ | 1.70 | 2.63E-02 |
| Q9H756 | Leucine-rich repeat-containing protein 19 | ↑ | 1.70 | 2.39E-02 |
| P02753 | Retinol-binding protein 4 | ↑ | 1.71 | 3.13E-02 |
| P04004 | Vitronectin | ↑ | 1.71 | 1.99E-02 |
| Q9NUM4 | Transmembrane protein 106B | ↑ | 1.72 | 4.39E-02 |
| O14773 | Tripeptidyl-peptidase 1 | ↑ | 1.75 | 2.54E-02 |
| A0A075B6S2 | Immunoglobulin kappa variable 2D-29 | ↑ | 1.75 | 3.89E-02 |
| Q92820 | Gamma-glutamyl hydrolase | ↑ | 1.75 | 5.11E-03 |
| P48061 | Stromal cell-derived factor 1 | ↑ | 1.75 | 4.22E-02 |
| Q5JS37 | NHL repeat-containing protein 3 | ↑ | 1.76 | 3.01E-03 |
| Q9NPG4 | Protocadherin-12 | ↑ | 1.76 | 8.40E-03 |
| M0QZG5 | CD209 antigen, isoform CRA_j | ↑ | 1.76 | 2.70E-02 |
| O75339 | Cartilage intermediate layer protein 1 | ↑ | 1.77 | 3.22E-02 |
| P01834 | Immunoglobulin kappa constant | ↑ | 1.77 | 8.37E-03 |
| Q16769 | Glutaminyl-peptide cyclotransferase | ↑ | 1.78 | 2.03E-02 |
| K3W4U1 | Fc epsilon receptor II | ↑ | 1.78 | 2.81E-02 |
| Q12864 | Cadherin-17 | ↑ | 1.78 | 4.84E-02 |
| P01859 | Immunoglobulin heavy constant gamma 2 | ↑ | 1.78 | 1.26E-02 |
| P00450 | Ceruloplasmin | ↑ | 1.79 | 3.16E-02 |
| J3KPQ0 | Fibroblast growth factor receptor | ↑ | 1.80 | 2.32E-02 |
| P19961 | Alpha-amylase 2B | ↑ | 1.80 | 2.34E-02 |
| Q9H461 | Frizzled-8 | ↑ | 1.81 | 4.49E-04 |
| P20933 | N(4)-(beta-N-acetylglucosaminyl)-L-asparaginase | ↑ | 1.83 | 2.50E-02 |
| P02749 | Beta-2-glycoprotein 1 | ↑ | 1.85 | 2.12E-02 |
| A0A087WTY6 | Neuroblastoma suppressor of tumorigenicity 1 | ↑ | 1.85 | 4.84E-04 |
| A0A0C4DH38 | Immunoglobulin heavy variable 5-51 | ↑ | 1.85 | 2.88E-02 |
| Q96IU4 | Putative protein-lysine deacylase ABHD14B | ↑ | 1.86 | 5.27E-03 |
| A0A087WX80 | Laminin subunit alpha 2 | ↑ | 1.87 | 3.06E-02 |
| Q02413 | Desmoglein-1 | ↑ | 1.87 | 9.15E-03 |
| P48745 | CCN family member 3 | ↑ | 1.88 | 2.09E-02 |
| P10253 | Lysosomal alpha-glucosidase | ↑ | 1.88 | 6.66E-03 |
| Q96GW7 | Brevican core protein | ↑ | 1.89 | 2.02E-02 |
| Q9UFM8 | Neuroplastin | ↑ | 1.89 | 4.48E-02 |
| P55000 | Secreted Ly-6/uPAR-related protein 1 | ↑ | 1.90 | 2.68E-03 |
| A0A0C4DFZ2 | Arylsulfatase A | ↑ | 1.91 | 8.68E-03 |
| P10619 | Lysosomal protective protein | ↑ | 1.92 | 1.21E-02 |
| P20674 | Cytochrome c oxidase subunit 5A, mitochondrial | ↑ | 1.92 | 2.42E-02 |
| Q9H6B4 | CXADR-like membrane protein | ↑ | 1.94 | 2.61E-02 |
| A8MW49 | Fatty acid-binding protein, liver | ↑ | 1.95 | 9.39E-03 |
| A0A0A0MQS9 | Laminin subunit alpha 4 | ↑ | 1.96 | 3.60E-02 |
| A0A0U1RQC5 | Neurexin 3 | ↑ | 1.97 | 2.34E-02 |
| Q8NFT8 | Delta and Notch-like epidermal growth factor-related receptor | ↑ | 1.97 | 1.22E-02 |
| P22792 | Carboxypeptidase N subunit 2 | ↑ | 1.98 | 8.60E-04 |
| A0A0C4DH43 | Immunoglobulin heavy variable 2-70D | ↑ | 1.99 | 9.02E-03 |
| A6NC48 | ADP-ribosyl cyclase/cyclic ADP-ribose hydrolase | ↑ | 1.99 | 6.40E-03 |
| Q9Y646 | Carboxypeptidase Q | ↑ | 1.99 | 3.83E-03 |
| Q9Y376 | Calcium-binding protein 39 | ↑ | 2.01 | 1.42E-02 |
| O95445 | Apolipoprotein M | ↑ | 2.03 | 4.56E-03 |
| P02774 | Vitamin D-binding protein | ↑ | 2.03 | 1.64E-02 |
| P01210 | Proenkephalin-A | ↑ | 2.07 | 7.08E-03 |
| Q5T123 | SH3 domain-binding glutamic acid-rich-like protein 3 | ↑ | 2.08 | 3.02E-02 |
| P08473 | Neprilysin | ↑ | 2.09 | 2.87E-02 |
| A0A096LP69 | CD99 molecule | ↑ | 2.09 | 2.29E-02 |
| P10109 | Adrenodoxin, mitochondrial | ↑ | 2.09 | 3.44E-02 |
| Q13113 | PDZK1-interacting protein 1 | ↑ | 2.10 | 6.97E-03 |
| P50591 | Tumor necrosis factor ligand superfamily member 10 | ↑ | 2.10 | 9.61E-03 |
| O43493 | Trans-Golgi network integral membrane protein 2 | ↑ | 2.10 | 2.28E-03 |
| P14151 | L-selectin | ↑ | 2.12 | 4.79E-03 |
| Q9HB75 | p53-induced death domain-containing protein 1 | ↑ | 2.15 | 7.11E-03 |
| A0A0G2JNI0 | Transmembrane protease serine | ↑ | 2.17 | 4.90E-02 |
| O96009 | Napsin-A | ↑ | 2.17 | 3.54E-02 |
| P30047 | GTP cyclohydrolase 1 feedback regulatory protein | ↑ | 2.19 | 7.16E-03 |
| P01024 | Complement C3 | ↑ | 2.19 | 4.47E-02 |
| Q14019 | Coactosin-like protein | ↑ | 2.21 | 1.10E-02 |
| P31025 | Lipocalin-1 | ↑ | 2.22 | 9.26E-03 |
| O95841 | Angiopoietin-related protein 1 | ↑ | 2.23 | 1.58E-02 |
| P02787 | Serotransferrin | ↑ | 2.23 | 4.56E-02 |
| P01011 | Alpha-1-antichymotrypsin | ↑ | 2.24 | 1.09E-02 |
| Q13443 | Disintegrin and metalloproteinase domain-containing protein 9 | ↑ | 2.27 | 4.49E-02 |
| P32320 | Cytidine deaminase | ↑ | 2.27 | 4.55E-02 |
| P04196 | Histidine-rich glycoprotein | ↑ | 2.27 | 6.06E-03 |
| P07437 | Tubulin beta chain | ↑ | 2.32 | 3.34E-02 |
| P05556 | Integrin beta-1 | ↑ | 2.33 | 1.44E-02 |
| Q9UJ72 | Annexin A10 | ↑ | 2.33 | 4.24E-02 |
| P32004 | Neural cell adhesion molecule L1 | ↑ | 2.35 | 2.14E-02 |
| O00592 | Podocalyxin | ↑ | 2.36 | 2.86E-02 |
| B1APH0 | Basonuclin 2 | ↑ | 2.36 | 1.97E-02 |
| A0A087WYX9 | Collagen type V alpha 2 chain | ↑ | 2.37 | 1.38E-03 |
| P01111 | GTPase NRas | ↑ | 2.38 | 1.91E-02 |
| A0A3B3IUC4 | Alpha-galactosidase | ↑ | 2.38 | 4.72E-02 |
| Q68CJ9 | Cyclic AMP-responsive element-binding protein 3-like protein 3 | ↑ | 2.38 | 3.32E-03 |
| Q86TY3 | Armadillo-like helical domain-containing protein 4 | ↑ | 2.39 | 8.94E-03 |
| P25391 | Laminin subunit alpha-1 | ↑ | 2.40 | 2.17E-02 |
| G5E9G7 | Neurexin 2 | ↑ | 2.41 | 1.84E-02 |
| O75487 | Glypican-4 | ↑ | 2.42 | 3.92E-02 |
| I3L0L6 | E3 ubiquitin-protein ligase RNF167 | ↑ | 2.43 | 4.72E-02 |
| P12830 | Cadherin-1 | ↑ | 2.43 | 1.26E-02 |
| P49788 | Retinoic acid receptor responder protein 1 | ↑ | 2.44 | 1.14E-02 |
| A0A3B3IRL2 | Cellular repressor of E1A stimulated genes 1 | ↑ | 2.48 | 2.31E-02 |
| A0A087WZW1 | Colipase | ↑ | 2.54 | 6.76E-03 |
| A0A0J9YX35 | Immunoglobulin heavy variable 3-64D | ↑ | 2.55 | 4.92E-03 |
| K7EKI0 | Envoplakin | ↑ | 2.55 | 3.19E-02 |
| A0A286YFJ8 | Immunoglobulin heavy constant gamma 4 | ↑ | 2.57 | 2.05E-02 |
| P29401 | Transketolase | ↑ | 2.60 | 3.61E-02 |
| P49006 | MARCKS-related protein | ↑ | 2.66 | 3.09E-02 |
| P01178 | Oxytocin-neurophysin 1 | ↑ | 2.66 | 8.71E-04 |
| P06858 | Lipoprotein lipase | ↑ | 2.68 | 4.66E-02 |
| O94919 | Endonuclease domain-containing 1 protein | ↑ | 2.68 | 5.88E-04 |
| P01019 | Angiotensinogen | ↑ | 2.69 | 1.67E-02 |
| P30046 | D-dopachrome decarboxylase | ↑ | 2.70 | 2.46E-02 |
| Q14118 | Dystroglycan 1 | ↑ | 2.71 | 3.33E-03 |
| P06731 | Carcinoembryonic antigen-related cell adhesion molecule 5 | ↑ | 2.74 | 2.32E-02 |
| Q9Y4D7 | Plexin-D1 | ↑ | 2.74 | 2.14E-02 |
| P02765 | Alpha-2-HS-glycoprotein | ↑ | 2.75 | 3.22E-03 |
| Q86UD1 | Out at first protein homolog | ↑ | 2.79 | 7.73E-03 |
| Q92626 | Peroxidasin homolog | ↑ | 2.82 | 9.41E-03 |
| P28072 | Proteasome subunit beta type-6 | ↑ | 2.82 | 4.29E-02 |
| F5H0U5 | Glycolipid transfer protein | ↑ | 2.86 | 8.92E-03 |
| P16989 | Y-box-binding protein 3 | ↑ | 2.87 | 4.64E-03 |
| P11766 | Alcohol dehydrogenase class-3 | ↑ | 2.88 | 1.87E-04 |
| H3BMA1 | Mesothelin | ↑ | 2.88 | 4.37E-02 |
| P12318 | Low affinity immunoglobulin gamma Fc region receptor II-a | ↑ | 2.89 | 9.59E-03 |
| A0A1B0GTG2 | Aldehyde dehydrogenase 7 family member A1 | ↑ | 2.89 | 1.63E-02 |
| P27348 | 14-3-3 protein theta | ↑ | 2.91 | 1.20E-02 |
| M0QYN0 | Myeloid derived growth factor | ↑ | 2.93 | 4.53E-02 |
| Q6PI73 | Leukocyte immunoglobulin-like receptor subfamily A member 6 | ↑ | 2.94 | 1.49E-02 |
| A0A0G2JSC0 | Immunoglobulin lambda variable 5-45 | ↑ | 2.94 | 3.71E-03 |
| P29692 | Elongation factor 1-delta | ↑ | 2.96 | 7.19E-04 |
| Q9BY89 | Uncharacterized protein KIAA1671 | ↑ | 2.96 | 2.67E-02 |
| P54826 | Growth arrest-specific protein 1 | ↑ | 2.97 | 5.66E-03 |
| Q8N8N7 | Prostaglandin reductase 2 | ↑ | 2.98 | 4.54E-02 |
| Q92496 | Complement factor H-related protein 4 | ↑ | 3.01 | 6.77E-04 |
| Q16348 | Solute carrier family 15 member 2 | ↑ | 3.03 | 3.53E-02 |
| P48304 | Lithostathine-1-beta | ↑ | 3.11 | 3.93E-02 |
| P17655 | Calpain-2 catalytic subunit | ↑ | 3.12 | 1.77E-03 |
| Q7Z794 | Keratin, type II cytoskeletal 1b | ↑ | 3.15 | 5.08E-03 |
| P22304 | Iduronate 2-sulfatase | ↑ | 3.17 | 6.17E-03 |
| P30044 | Peroxiredoxin-5 | ↑ | 3.20 | 4.60E-03 |
| Q9UJ99 | Cadherin-22 | ↑ | 3.23 | 4.90E-02 |
| B7Z4G8 | Amyloid beta precursor like protein 1 | ↑ | 3.24 | 4.41E-02 |
| C9JM33 | Interferon alpha and beta receptor subunit 2 | ↑ | 3.25 | 1.82E-02 |
| P52961 | GPI-linked NAD | ↑ | 3.28 | 2.00E-02 |
| Q5T2L0 | V-set domain containing T cell activation inhibitor 1 | ↑ | 3.29 | 2.81E-02 |
| P68371 | Tubulin beta-4B chain | ↑ | 3.39 | 2.01E-03 |
| P36941 | Tumor necrosis factor receptor superfamily member 3 | ↑ | 3.40 | 3.00E-02 |
| A0A0C4DH72 | Immunoglobulin kappa variable 1-6 | ↑ | 3.41 | 2.75E-03 |
| P68363 | Tubulin alpha-1B chain | ↑ | 3.51 | 1.25E-02 |
| P30475 | HLA class I histocompatibility antigen, B alpha chain | ↑ | 3.60 | 2.66E-03 |
| P09525 | Annexin A4 | ↑ | 3.62 | 3.83E-02 |
| P12931 | Proto-oncogene tyrosine-protein kinase Src | ↑ | 3.62 | 2.29E-04 |
| O75891 | Cytosolic 10-formyltetrahydrofolate dehydrogenase | ↑ | 3.65 | 2.68E-02 |
| P50395 | Rab GDP dissociation inhibitor beta | ↑ | 3.67 | 1.36E-02 |
| Q92484 | Acid sphingomyelinase-like phosphodiesterase 3a | ↑ | 3.74 | 7.03E-03 |
| A0A0A0MSA0 | Laminin subunit alpha-3 | ↑ | 3.74 | 4.77E-02 |
| A0A0D9SG04 | Cordon-bleu WH2 repeat protein like 1 | ↑ | 3.74 | 1.96E-02 |
| A0A075B6K4 | Immunoglobulin lambda variable 3-10 | ↑ | 3.93 | 2.77E-02 |
| A0A1W2PNV4 | Actin-related protein 2/3 complex subunit 1A | ↑ | 3.94 | 4.88E-02 |
| Q9NS68 | Tumor necrosis factor receptor superfamily member 19 | ↑ | 4.08 | 1.10E-02 |
| Q9BXJ7 | Protein amnionless | ↑ | 4.13 | 2.03E-02 |
| Q8WWY7 | WAP four-disulfide core domain protein 12 | ↑ | 4.21 | 3.05E-02 |
| P05186 | Alkaline phosphatase, tissue-nonspecific isozyme | ↑ | 4.21 | 1.33E-02 |
| Q96A22 | Uncharacterized protein C11orf52 | ↑ | 4.27 | 4.93E-03 |
| Q15181 | Inorganic pyrophosphatase | ↑ | 4.35 | 2.72E-02 |
| E9PL83 | Pro-adrenomedullin | ↑ | 4.39 | 6.36E-03 |
| Q15848 | Adiponectin | ↑ | 4.44 | 2.37E-02 |
| Q3LXA3 | Triokinase/FMN cyclase | ↑ | 4.45 | 2.33E-02 |
| Q15149 | Plectin | ↑ | 4.55 | 1.72E-02 |
| Q6UXN8 | C-type lectin domain family 9 member A | ↑ | 4.65 | 4.80E-02 |
| Q04917 | 14-3-3 protein eta | ↑ | 4.67 | 3.66E-02 |
| F5H1S8 | Malectin | ↑ | 4.67 | 1.41E-03 |
| Q15833 | Syntaxin-binding protein 2 | ↑ | 4.73 | 3.36E-03 |
| E7END7 | Ras-related protein Rab-1A | ↑ | 4.78 | 2.11E-02 |
| K7EIK7 | EMAP like 2 | ↑ | 4.79 | 2.38E-02 |
| Q8N436 | Inactive carboxypeptidase-like protein X2 | ↑ | 4.80 | 4.80E-02 |
| P04279 | Semenogelin-1 | ↑ | 4.86 | 1.63E-03 |
| E7EMR3 | Adhesion G protein-coupled receptor L3 | ↑ | 4.87 | 6.15E-04 |
| Q9UHI8 | A disintegrin and metalloproteinase with thrombospondin motifs 1 | ↑ | 4.89 | 4.37E-03 |
| A0A075B788 | Protein tyrosine phosphatase receptor type C | ↑ | 4.90 | 2.91E-02 |
| D6RHW5 | Endomucin | ↑ | 4.92 | 3.15E-02 |
| A0A0B4J2C3 | Translationally-controlled tumor protein | ↑ | 4.95 | 4.63E-02 |
| K7ELM9 | Apolipoprotein C1 | ↑ | 5.05 | 2.39E-04 |
| Q8NFY4 | Semaphorin-6D | ↑ | 5.09 | 3.00E-02 |
| P14091 | Cathepsin E | ↑ | 5.14 | 2.63E-02 |
| O43396 | Thioredoxin-like protein 1 | ↑ | 5.17 | 4.49E-02 |
| Q56A73 | Spindlin-4 | ↑ | 5.19 | 1.11E-02 |
| Q1EHB4 | Sodium-coupled monocarboxylate transporter 2 | ↑ | 5.19 | 1.60E-02 |
| P22894 | Neutrophil collagenase | ↑ | 5.19 | 1.99E-02 |
| Q14142 | Tripartite motif-containing protein 14 | ↑ | 5.20 | 2.13E-02 |
| P21583 | Kit ligand | ↑ | 5.20 | 4.34E-02 |
| Q9BY43 | Charged multivesicular body protein 4a | ↑ | 5.25 | 1.61E-02 |
| B0YIW2 | Apolipoprotein C-III | ↑ | 5.27 | 2.25E-02 |
| E9PGC5 | protein-tyrosine-phosphatase | ↑ | 5.29 | 9.03E-03 |
| P26641 | Elongation factor 1-gamma | ↑ | 5.33 | 1.39E-02 |
| J3KNP4 | Semaphorin-4B | ↑ | 5.35 | 4.36E-02 |
| Q9UHG3 | Prenylcysteine oxidase 1 | ↑ | 5.41 | 4.06E-02 |
| H7BY57 | Neurofascin | ↑ | 5.49 | 1.68E-03 |
| J3KNF4 | Superoxide dismutase copper chaperone | ↑ | 5.54 | 4.95E-02 |
| F5H4M7 | Transmembrane p24 trafficking protein 3 | ↑ | 5.60 | 4.49E-02 |
| A0A0A0MSS8 | Aldo-keto reductase family 1 member C3 | ↑ | 5.71 | 1.28E-02 |
| P25786 | Proteasome subunit alpha type-1 | ↑ | 5.71 | 1.14E-03 |
| A0A0U1RQV3 | EGF containing fibulin extracellular matrix protein 1 | ↑ | 5.79 | 3.16E-02 |
| P12277 | Creatine kinase B-type | ↑ | 5.81 | 1.49E-02 |
| Q8WU39 | Marginal zone B- and B1-cell-specific protein | ↑ | 5.89 | 7.74E-03 |
| A0A0C4DGE4 | Syntaxin 3 | ↑ | 5.92 | 1.72E-02 |
| A0A087WSY5 | Carboxypeptidase B2 | ↑ | 6.13 | 3.36E-02 |
| P31327 | Carbamoyl-phosphate synthase [ammonia], mitochondrial | ↑ | 6.27 | 3.68E-02 |
| Q9NRA1 | Platelet-derived growth factor C | ↑ | 6.32 | 4.69E-02 |
| O00764 | Pyridoxal kinase | ↑ | 6.39 | 4.16E-02 |
| A0A3B3IQ51 | Complement factor H related 2 | ↑ | 6.53 | 2.01E-02 |
| P00533 | Epidermal growth factor receptor | ↑ | 6.66 | 4.06E-02 |
| A0A0A0MQV3 | Kin of IRRE-like protein 2 | ↑ | 6.97 | 3.57E-03 |
| Q6ZMJ2 | Scavenger receptor class A member 5 | ↑ | 7.05 | 1.67E-02 |
| O00401 | Actin nucleation-promoting factor WASL | ↑ | 7.12 | 3.17E-04 |
| Q6H9L7 | Isthmin-2 | ↑ | 7.16 | 3.18E-02 |
| O60449 | Lymphocyte antigen 75 | ↑ | 7.20 | 3.83E-03 |
| Q86SF2 | N-acetylgalactosaminyltransferase 7 | ↑ | 7.35 | 4.67E-03 |
| Q8IZJ3 | C3 and PZP-like alpha-2-macroglobulin domain-containing protein 8 | ↑ | 7.43 | 8.86E-03 |
| B5MBX2 | Transcobalamin-2 | ↑ | 7.43 | 4.49E-02 |
| O00144 | Frizzled-9 | ↑ | 8.31 | 2.02E-03 |
| A0A2R8YE63 | Epidermal growth factor receptor pathway substrate 8 | ↑ | 8.93 | 1.35E-02 |
| P24298 | Alanine aminotransferase 1 | ↑ | 9.03 | 3.68E-02 |
| Q9BR76 | Coronin-1B | ↑ | 9.62 | 3.28E-02 |
| P00742 | Coagulation factor X | ↑ | 10.39 | 4.09E-03 |
| Q9HCH3 | Copine-5 | ↑ | 10.69 | 2.73E-02 |
| H0Y4H3 | CD99 antigen-like protein 2 | ↑ | 10.74 | 1.19E-02 |
| O95154 | Aflatoxin B1 aldehyde reductase member 3 | ↑ | 11.67 | 1.11E-02 |
| O95817 | BAG family molecular chaperone regulator 3 | ↑ | 11.68 | 4.72E-03 |
| Q8N8Z6 | Discoidin, CUB and LCCL domain-containing protein 1 | ↑ | 12.44 | 2.60E-02 |
| O60784 | Target of Myb1 membrane trafficking protein | ↑ | 12.51 | 1.32E-02 |
| A5D8V6 | Vacuolar protein sorting-associated protein 37C | ↑ | 13.18 | 7.35E-03 |
| P98161 | Polycystin-1 | ↑ | 13.42 | 2.48E-02 |
| Q9P1F3 | Costars family protein ABRACL | ↑ | 13.76 | 1.90E-02 |
| P00568 | Adenylate kinase isoenzyme 1 | ↑ | 15.07 | 1.19E-02 |
| Q9H190 | Syntenin-2 | ↑ | 21.77 | 9.21E-03 |

TableS4 Biological Processes and Signaling Pathways Enriched for Differential Proteins Produced in the Healthy Group Compared to the Depressed Group Under Relaxed Conditions(P<0.05)

| **BP** | **P-Value** | **Pathway** | **P-Value** |
| --- | --- | --- | --- |
| cell adhesion | 7.40E-14 | Cell adhesion molecules | 9.70E-07 |
| negative regulation of fibrinolysis | 5.50E-07 | Lysosome | 4.90E-06 |
| complement activation, classical pathway | 1.20E-06 | ECM-receptor interaction | 3.30E-05 |
| axon guidance | 1.30E-05 | PI3K-Akt signaling pathway | 7.80E-05 |
| homophilic cell adhesion via plasma membrane adhesion molecules | 2.90E-05 | Focal adhesion | 8.50E-05 |
| positive regulation of B cell activation | 6.50E-05 | Amoebiasis | 9.80E-05 |
| leukocyte cell-cell adhesion | 6.90E-05 | Phagosome | 1.00E-04 |
| phagocytosis, recognition | 7.00E-05 | Complement and coagulation cascades | 1.70E-04 |
| proteolysis | 7.80E-05 | Regulation of actin cytoskeleton | 2.80E-04 |
| B cell receptor signaling pathway | 1.10E-04 | Proteoglycans in cancer | 3.70E-04 |
| cell migration | 1.20E-04 | Human papillomavirus infection | 1.10E-03 |
| regulation of cell adhesion | 1.70E-04 | Pathogenic Escherichia coli infection | 3.30E-03 |
| phagocytosis, engulfment | 2.00E-04 | Gap junction | 5.40E-03 |
| extracellular matrix organization | 2.50E-04 | Adherens junction | 7.00E-03 |
| antibacterial humoral response | 2.90E-04 | Pathways in cancer | 8.20E-03 |
| peptidyl-tyrosine phosphorylation | 3.90E-04 | Protein digestion and absorption | 1.10E-02 |
| calcium-dependent cell-cell adhesion via plasma membrane cell adhesion molecules | 4.90E-04 | Leukocyte transendothelial migration | 1.80E-02 |
| adaptive immune response | 6.20E-04 | Carbon metabolism | 1.90E-02 |
| positive regulation of MAP kinase activity | 6.60E-04 | Legionellosis | 2.00E-02 |
| positive regulation of extrinsic apoptotic signaling pathway | 7.10E-04 | Galactose metabolism | 2.00E-02 |
| cell-matrix adhesion | 7.60E-04 | Viral myocarditis | 2.40E-02 |
| female pregnancy | 9.80E-04 | Small cell lung cancer | 2.70E-02 |
| response to glucocorticoid | 1.10E-03 | AGE-RAGE signaling pathway in diabetic complications | 3.60E-02 |
| adherens junction organization | 1.20E-03 | Apoptosis | 3.90E-02 |
| immunoglobulin mediated immune response | 1.20E-03 | Bladder cancer | 3.90E-02 |
| cellular response to amino acid stimulus | 1.30E-03 | Rap1 signaling pathway | 3.90E-02 |
| immune response | 1.50E-03 | Endocytosis | 4.10E-02 |
| cell-cell adhesion | 1.50E-03 | NF-kappa B signaling pathway | 4.20E-02 |
| positive regulation of peptidyl-tyrosine phosphorylation | 1.60E-03 | Melanoma | 4.30E-02 |
| cell surface receptor signaling pathway | 1.70E-03 | Alzheimer disease | 4.90E-02 |
| positive regulation of protein kinase B signaling | 2.00E-03 | - | - |
| translational elongation | 2.10E-03 | - | - |
| defense response to bacterium | 2.20E-03 | - | - |
| cell adhesion mediated by integrin | 2.30E-03 | - | - |
| skeletal system development | 2.50E-03 | - | - |
| viral entry into host cell | 2.80E-03 | - | - |
| animal organ morphogenesis | 3.00E-03 | - | - |
| endocytosis | 3.00E-03 | - | - |
| basement membrane organization | 3.20E-03 | - | - |
| synapse assembly | 3.30E-03 | - | - |
| positive regulation of cell migration | 3.40E-03 | - | - |
| regulation of peptidyl-tyrosine phosphorylation | 3.70E-03 | - | - |
| positive regulation of neutrophil extravasation | 3.80E-03 | - | - |
| negative regulation of monocyte chemotaxis | 3.80E-03 | - | - |
| collagen fibril organization | 3.90E-03 | - | - |
| response to food | 4.70E-03 | - | - |
| cellular response to reactive oxygen species | 4.80E-03 | - | - |
| zymogen activation | 5.20E-03 | - | - |
| complement activation | 5.20E-03 | - | - |
| signal transduction | 5.60E-03 | - | - |
| innate immune response | 6.00E-03 | - | - |
| renal absorption | 6.30E-03 | - | - |
| heterophilic cell-cell adhesion via plasma membrane cell adhesion molecules | 6.40E-03 | - | - |
| leukocyte migration | 6.50E-03 | - | - |
| protein autoprocessing | 6.50E-03 | - | - |
| positive regulation of phosphatidylinositol 3-kinase signaling | 6.60E-03 | - | - |
| lysosome organization | 6.80E-03 | - | - |
| negative regulation of endopeptidase activity | 7.20E-03 | - | - |
| cellular response to cAMP | 7.70E-03 | - | - |
| angiogenesis | 8.80E-03 | - | - |
| carbohydrate metabolic process | 1.10E-02 | - | - |
| regulation of blood coagulation | 1.10E-02 | - | - |
| cytolysis by host of symbiont cells | 1.10E-02 | - | - |
| tissue development | 1.20E-02 | - | - |
| neuromuscular process controlling posture | 1.30E-02 | - | - |
| positive regulation of integrin-mediated signaling pathway | 1.30E-02 | - | - |
| triglyceride metabolic process | 1.40E-02 | - | - |
| cell-cell adhesion via plasma-membrane adhesion molecules | 1.50E-02 | - | - |
| negative regulation of blood coagulation | 1.50E-02 | - | - |
| positive regulation of tumor necrosis factor production | 1.50E-02 | - | - |
| positive regulation of NIK/NF-kappaB signaling | 1.60E-02 | - | - |
| central nervous system development | 1.60E-02 | - | - |
| antimicrobial humoral immune response mediated by antimicrobial peptide | 1.70E-02 | - | - |
| negative regulation of intrinsic apoptotic signaling pathway in response to DNA damage | 1.70E-02 | - | - |
| peptide metabolic process | 1.70E-02 | - | - |
| positive regulation of cell proliferation | 1.80E-02 | - | - |
| integrin-mediated signaling pathway | 1.80E-02 | - | - |
| cellular oxidant detoxification | 1.90E-02 | - | - |
| high-density lipoprotein particle remodeling | 2.00E-02 | - | - |
| regulation of cell migration | 2.00E-02 | - | - |
| cellular response to mechanical stimulus | 2.00E-02 | - | - |
| protein catabolic process | 2.00E-02 | - | - |
| positive regulation of blood pressure | 2.20E-02 | - | - |
| receptor-mediated endocytosis | 2.20E-02 | - | - |
| kidney development | 2.30E-02 | - | - |
| cell-cell junction assembly | 2.50E-02 | - | - |
| regulation of embryonic development | 2.60E-02 | - | - |
| fibrinolysis | 2.70E-02 | - | - |
| negative regulation of anoikis | 2.70E-02 | - | - |
| triglyceride catabolic process | 3.00E-02 | - | - |
| response to peptide hormone | 3.10E-02 | - | - |
| myoblast differentiation | 3.30E-02 | - | - |
| lipoprotein metabolic process | 3.30E-02 | - | - |
| establishment of endothelial barrier | 3.30E-02 | - | - |
| positive regulation of ERK1 and ERK2 cascade | 3.60E-02 | - | - |
| positive regulation of vascular smooth muscle cell proliferation | 3.60E-02 | - | - |
| positive regulation of receptor-mediated endocytosis | 3.60E-02 | - | - |
| viral budding via host ESCRT complex | 3.60E-02 | - | - |
| lysosomal lumen acidification | 3.60E-02 | - | - |
| positive regulation of smooth muscle cell migration | 3.60E-02 | - | - |
| cell morphogenesis | 4.00E-02 | - | - |
| T cell extravasation | 4.10E-02 | - | - |
| susceptibility to natural killer cell mediated cytotoxicity | 4.10E-02 | - | - |
| smooth muscle cell proliferation | 4.10E-02 | - | - |
| adhesion of symbiont to host | 4.10E-02 | - | - |
| oligopeptide transport | 4.10E-02 | - | - |
| bone regeneration | 4.10E-02 | - | - |
| blood coagulation, common pathway | 4.10E-02 | - | - |
| positive regulation of vascular associated smooth muscle cell migration | 4.20E-02 | - | - |
| negative regulation of cell migration | 4.20E-02 | - | - |
| apoptotic process | 4.50E-02 | - | - |
| cellular response to platelet-derived growth factor stimulus | 4.60E-02 | - | - |
| immune system process | 4.70E-02 | - | - |
| positive regulation of cell adhesion | 4.70E-02 | - | - |
| response to lipopolysaccharide | 4.70E-02 | - | - |
| multicellular organism development | 4.80E-02 | - | - |

TableS5 Differential proteins in bipolar patients compared to depressed patients (FC≥1.5 or ≤0.67, P<0.05)

| **Accession** | **Protein names** | **Trend** | **FC** | **P-Value** |
| --- | --- | --- | --- | --- |
| A0A0U1RR32 | Histone H2A | ↓ | 0.38 | 3.03E-02 |
| Q53TN4 | Plasma membrane ascorbate-dependent reductase CYBRD1 | ↓ | 0.48 | 7.17E-03 |
| F8VNT9 | CD63 molecule | ↓ | 0.48 | 5.11E-03 |
| Q9H1C7 | Cysteine-rich and transmembrane domain-containing protein 1 | ↓ | 0.48 | 4.29E-03 |
| P20073 | Annexin A7 | ↓ | 0.54 | 1.13E-02 |
| O60293 | Zinc finger C3H1 domain-containing protein | ↓ | 0.56 | 1.40E-02 |
| Q9BUT1 | Dehydrogenase/reductase SDR family member 6 | ↓ | 0.57 | 4.12E-02 |
| P05154 | Plasma serine protease inhibitor | ↓ | 0.57 | 1.29E-02 |
| Q9BXP8 | Pappalysin-2 | ↓ | 0.57 | 3.30E-02 |
| P16278 | Beta-galactosidase | ↓ | 0.58 | 4.75E-02 |
| O43895 | Xaa-Pro aminopeptidase 2 | ↓ | 0.59 | 3.60E-02 |
| Q8WUM4 | Programmed cell death 6-interacting protein | ↓ | 0.59 | 2.27E-02 |
| P26038 | Moesin | ↓ | 0.60 | 6.23E-03 |
| Q9NZH0 | G-protein coupled receptor family C group 5 member B | ↓ | 0.60 | 2.27E-02 |
| Q8IWA5 | Choline transporter-like protein 2 | ↓ | 0.61 | 1.47E-02 |
| A0A0C4DFY5 | G protein-coupled receptor class C group 5 member C | ↓ | 0.61 | 1.08E-02 |
| P16870 | Carboxypeptidase E | ↓ | 0.63 | 4.24E-02 |
| O00115 | Deoxyribonuclease-2-alpha | ↓ | 0.64 | 4.42E-02 |
| O43633 | Charged multivesicular body protein 2a | ↓ | 0.65 | 8.83E-03 |
| Q7LBR1 | Charged multivesicular body protein 1b | ↓ | 0.65 | 3.37E-02 |
| P23284 | Peptidyl-prolyl cis-trans isomerase B | ↓ | 0.65 | 1.01E-03 |
| Q9H3G5 | Probable serine carboxypeptidase CPVL | ↓ | 0.65 | 4.57E-02 |
| A0A2R8Y5S7 | Radixin | ↓ | 0.66 | 1.26E-02 |
| P19440 | Glutathione hydrolase 1 proenzyme | ↓ | 0.66 | 1.14E-02 |
| Q08174 | Protocadherin-1 | ↑ | 1.50 | 3.61E-03 |
| O75594 | Peptidoglycan recognition protein 1 | ↑ | 1.50 | 2.77E-03 |
| A0M8Q6 | Immunoglobulin lambda constant 7 | ↑ | 1.50 | 4.62E-03 |
| Q24JP5 | Transmembrane protein 132A | ↑ | 1.51 | 3.29E-02 |
| Q06828 | Fibromodulin | ↑ | 1.51 | 2.54E-02 |
| P61981 | 14-3-3 protein gamma | ↑ | 1.51 | 2.62E-02 |
| A0A087WYL5 | Seizure related 6 homolog like 2 | ↑ | 1.51 | 2.22E-02 |
| P98172 | Ephrin-B1 | ↑ | 1.51 | 2.65E-02 |
| O43866 | CD5 antigen-like | ↑ | 1.52 | 3.30E-02 |
| G3V3X5 | Latent transforming growth factor beta binding protein 2 | ↑ | 1.52 | 2.02E-03 |
| Q9UM22 | Mammalian ependymin-related protein 1 | ↑ | 1.52 | 2.74E-02 |
| Q9BVM4 | Gamma-glutamylaminecyclotransferase | ↑ | 1.52 | 4.25E-02 |
| E5RHN3 | Hepatitis A virus cellular receptor 2 | ↑ | 1.52 | 1.55E-02 |
| P05362 | Intercellular adhesion molecule 1 | ↑ | 1.52 | 4.01E-02 |
| B1ALM4 | Transmembrane protein 9 | ↑ | 1.52 | 4.14E-02 |
| P98095 | Fibulin-2 | ↑ | 1.52 | 4.52E-02 |
| P15529 | Membrane cofactor protein | ↑ | 1.53 | 4.76E-02 |
| F8VUF6 | Decorin | ↑ | 1.53 | 3.22E-02 |
| P08138 | Tumor necrosis factor receptor superfamily member 16 | ↑ | 1.53 | 1.00E-02 |
| P18206 | Vinculin | ↑ | 1.54 | 9.94E-03 |
| Q14376 | UDP-glucose 4-epimerase | ↑ | 1.55 | 1.31E-02 |
| P01701 | Immunoglobulin lambda variable 1-51 | ↑ | 1.55 | 2.85E-02 |
| F6Q0M4 | TNF receptor superfamily member 14 | ↑ | 1.55 | 7.69E-03 |
| P09758 | Tumor-associated calcium signal transducer 2 | ↑ | 1.55 | 4.97E-02 |
| Q86V85 | Integral membrane protein GPR180 | ↑ | 1.55 | 1.47E-02 |
| Q02487 | Desmocollin-2 | ↑ | 1.55 | 2.37E-02 |
| P35754 | Glutaredoxin-1 | ↑ | 1.56 | 1.48E-02 |
| P36955 | Pigment epithelium-derived factor | ↑ | 1.56 | 4.85E-02 |
| Q9UIB8 | SLAM family member 5 | ↑ | 1.56 | 3.24E-03 |
| Q14508 | WAP four-disulfide core domain protein 2 | ↑ | 1.56 | 1.11E-02 |
| Q9NPY3 | Complement component C1q receptor | ↑ | 1.56 | 4.49E-02 |
| Q01459 | Di-N-acetylchitobiase | ↑ | 1.57 | 2.39E-02 |
| P10809 | 60 kDa heat shock protein, mitochondrial | ↑ | 1.57 | 1.73E-02 |
| O94910 | Adhesion G protein-coupled receptor L1 | ↑ | 1.57 | 6.58E-03 |
| H0Y3Z8 | IFNAR2-IL10RB readthrough | ↑ | 1.58 | 3.70E-02 |
| P13671 | Complement component C6 | ↑ | 1.58 | 2.35E-02 |
| A0A0B4J1X5 | Immunoglobulin heavy variable 3-74 | ↑ | 1.58 | 5.25E-03 |
| E9PMI0 | Layilin | ↑ | 1.59 | 1.16E-02 |
| B4DV12 | Ubiquitin B | ↑ | 1.59 | 4.90E-02 |
| B1AP13 | CD55 molecule | ↑ | 1.59 | 3.83E-02 |
| A0A0C4DGH0 | CD276 molecule | ↑ | 1.59 | 1.33E-02 |
| P78324 | Tyrosine-protein phosphatase non-receptor type substrate 1 | ↑ | 1.59 | 5.56E-03 |
| P16152 | Carbonyl reductase [NADPH] 1 | ↑ | 1.59 | 2.30E-02 |
| P13284 | Gamma-interferon-inducible lysosomal thiol reductase | ↑ | 1.60 | 1.31E-02 |
| Q92520 | Protein FAM3C | ↑ | 1.60 | 1.13E-03 |
| Q9Y646 | Carboxypeptidase Q | ↑ | 1.61 | 2.87E-03 |
| K7ELL7 | Glucosidase 2 subunit beta | ↑ | 1.62 | 6.07E-03 |
| P12814 | Alpha-actinin-1 | ↑ | 1.62 | 1.82E-03 |
| P20933 | N(4)-(beta-N-acetylglucosaminyl)-L-asparaginase | ↑ | 1.63 | 9.33E-03 |
| P15121 | Aldo-keto reductase family 1 member B1 | ↑ | 1.63 | 2.30E-02 |
| O96009 | Napsin-A | ↑ | 1.64 | 4.39E-02 |
| A0A096LP69 | CD99 molecule | ↑ | 1.64 | 4.84E-02 |
| P55290 | Cadherin-13 | ↑ | 1.64 | 3.38E-02 |
| P06681 | Complement C2 | ↑ | 1.64 | 1.31E-02 |
| A0A0U1RQC5 | Neurexin 3 | ↑ | 1.64 | 2.40E-02 |
| P80370 | Protein delta homolog 1 | ↑ | 1.65 | 1.07E-02 |
| O60939 | Sodium channel subunit beta-2 | ↑ | 1.66 | 3.81E-02 |
| Q12864 | Cadherin-17 | ↑ | 1.66 | 7.29E-03 |
| O75071 | EF-hand calcium-binding domain-containing protein 14 | ↑ | 1.66 | 1.86E-02 |
| P0DJD8 | Pepsin A-3 | ↑ | 1.66 | 4.87E-02 |
| A0A2R8Y430 | Glutathione synthetase | ↑ | 1.66 | 2.14E-02 |
| P0DOX8 | Immunoglobulin lambda-1 light chain | ↑ | 1.66 | 1.21E-02 |
| I3L192 | Basigin | ↑ | 1.66 | 1.68E-02 |
| A0A0A0MS43 | Solute carrier family 10 member 3 | ↑ | 1.66 | 2.54E-03 |
| P00747 | Plasminogen | ↑ | 1.67 | 1.47E-02 |
| J3KSN0 | Secreted and transmembrane 1 | ↑ | 1.68 | 1.21E-02 |
| Q07654 | Trefoil factor 3 | ↑ | 1.68 | 1.69E-02 |
| Q14126 | Desmoglein-2 | ↑ | 1.68 | 3.18E-02 |
| Q7Z4R8 | UPF0669 protein C6orf120 | ↑ | 1.68 | 9.72E-03 |
| Q96J84 | Kin of IRRE-like protein 1 | ↑ | 1.68 | 1.10E-03 |
| P0DOX5 | Immunoglobulin gamma-1 heavy chain | ↑ | 1.69 | 1.39E-03 |
| F8VVI9 | Cadherin 19 | ↑ | 1.69 | 1.18E-02 |
| Q6FHJ7 | Secreted frizzled-related protein 4 | ↑ | 1.70 | 1.63E-02 |
| P01717 | Immunoglobulin lambda variable 3-25 | ↑ | 1.70 | 3.16E-02 |
| P01178 | Oxytocin-neurophysin 1 | ↑ | 1.70 | 1.95E-02 |
| P02753 | Retinol-binding protein 4 | ↑ | 1.71 | 7.17E-03 |
| P10645 | Chromogranin-A | ↑ | 1.71 | 1.97E-02 |
| P19961 | Alpha-amylase 2B | ↑ | 1.72 | 4.17E-02 |
| E9PG71 | receptor protein-tyrosine kinase | ↑ | 1.72 | 3.00E-02 |
| P02768 | Albumin | ↑ | 1.72 | 1.18E-02 |
| P01834 | Immunoglobulin kappa constant | ↑ | 1.73 | 3.59E-02 |
| Q16581 | C3a anaphylatoxin chemotactic receptor | ↑ | 1.73 | 3.25E-02 |
| Q03403 | Trefoil factor 2 | ↑ | 1.73 | 3.11E-03 |
| P31997 | Carcinoembryonic antigen-related cell adhesion molecule 8 | ↑ | 1.73 | 1.83E-02 |
| Q02747 | Guanylin | ↑ | 1.74 | 4.13E-02 |
| P11362 | Fibroblast growth factor receptor 1 | ↑ | 1.74 | 1.00E-02 |
| Q16610 | Extracellular matrix protein 1 | ↑ | 1.75 | 4.59E-02 |
| Q04721 | Neurogenic locus notch homolog protein 2 | ↑ | 1.75 | 4.08E-03 |
| P54826 | Growth arrest-specific protein 1 | ↑ | 1.75 | 4.78E-02 |
| P27797 | Calreticulin | ↑ | 1.75 | 2.94E-03 |
| P26992 | Ciliary neurotrophic factor receptor subunit alpha | ↑ | 1.75 | 4.36E-03 |
| Q02413 | Desmoglein-1 | ↑ | 1.76 | 4.84E-03 |
| O76076 | CCN family member 5 | ↑ | 1.76 | 1.24E-02 |
| Q68CJ9 | Cyclic AMP-responsive element-binding protein 3-like protein 3 | ↑ | 1.77 | 1.02E-02 |
| Q9Y5E4 | Protocadherin beta-5 | ↑ | 1.77 | 2.81E-02 |
| J3QLM0 | CD7 molecule | ↑ | 1.77 | 1.26E-03 |
| P01210 | Proenkephalin-A | ↑ | 1.78 | 1.07E-02 |
| P41222 | Prostaglandin-H2 D-isomerase | ↑ | 1.79 | 4.14E-02 |
| Q99426 | Tubulin-folding cofactor B | ↑ | 1.80 | 1.92E-02 |
| Q68D85 | Natural cytotoxicity triggering receptor 3 ligand 1 | ↑ | 1.80 | 2.18E-03 |
| E7EV71 | Latent transforming growth factor beta binding protein 1 | ↑ | 1.80 | 3.11E-02 |
| P61769 | Beta-2-microglobulin | ↑ | 1.81 | 2.19E-02 |
| Q13308 | Inactive tyrosine-protein kinase 7 | ↑ | 1.81 | 4.85E-04 |
| Q16769 | Glutaminyl-peptide cyclotransferase | ↑ | 1.81 | 8.17E-03 |
| Q6ZVN8 | Hemojuvelin | ↑ | 1.83 | 4.96E-02 |
| Q7L266 | Isoaspartyl peptidase/L-asparaginase | ↑ | 1.83 | 2.78E-03 |
| P30085 | UMP-CMP kinase | ↑ | 1.83 | 3.72E-03 |
| A0A0A0MRJ7 | Coagulation factor V | ↑ | 1.83 | 3.51E-02 |
| P26842 | CD27 antigen | ↑ | 1.84 | 2.58E-03 |
| A0A0G2JSC0 | Immunoglobulin lambda variable 5-45 | ↑ | 1.84 | 4.05E-02 |
| Q92563 | Testican-2 | ↑ | 1.84 | 4.41E-02 |
| P22897 | Macrophage mannose receptor 1 | ↑ | 1.85 | 2.33E-02 |
| A0A075B6K5 | Immunoglobulin lambda variable 3-9 | ↑ | 1.85 | 3.20E-02 |
| B5A977 | Soluble TNFR1B variant 1 | ↑ | 1.85 | 9.52E-03 |
| A0A087X054 | Hypoxia up-regulated protein 1 | ↑ | 1.86 | 2.24E-02 |
| Q9H8L6 | Multimerin-2 | ↑ | 1.86 | 1.33E-03 |
| Q9BSG0 | Protease-associated domain-containing protein 1 | ↑ | 1.86 | 1.42E-02 |
| P98160 | Basement membrane-specific heparan sulfate proteoglycan core protein | ↑ | 1.86 | 9.08E-03 |
| A6NC48 | ADP-ribosyl cyclase/cyclic ADP-ribose hydrolase | ↑ | 1.86 | 1.82E-03 |
| A0A0J9YYC8 | Serine protease 2 | ↑ | 1.86 | 2.32E-02 |
| P31025 | Lipocalin-1 | ↑ | 1.86 | 1.99E-02 |
| P02749 | Beta-2-glycoprotein 1 | ↑ | 1.87 | 9.88E-04 |
| A0A0C4DH38 | Immunoglobulin heavy variable 5-51 | ↑ | 1.87 | 2.85E-03 |
| Q13291 | Signaling lymphocytic activation molecule | ↑ | 1.87 | 5.77E-03 |
| Q9BRA2 | Thioredoxin domain-containing protein 17 | ↑ | 1.87 | 9.54E-03 |
| O75084 | Frizzled-7 | ↑ | 1.88 | 2.69E-03 |
| A0A0J9YX35 | Immunoglobulin heavy variable 3-64D | ↑ | 1.88 | 1.48E-02 |
| Q9BQI0 | Allograft inflammatory factor 1-like | ↑ | 1.91 | 4.13E-02 |
| P50591 | Tumor necrosis factor ligand superfamily member 10 | ↑ | 1.91 | 2.38E-02 |
| Q9UGT4 | Sushi domain-containing protein 2 | ↑ | 1.91 | 3.31E-02 |
| Q8N307 | Mucin-20 | ↑ | 1.91 | 9.01E-03 |
| Q9Y279 | V-set and immunoglobulin domain-containing protein 4 | ↑ | 1.91 | 4.69E-02 |
| P12830 | Cadherin-1 | ↑ | 1.92 | 4.37E-02 |
| P15814 | Immunoglobulin lambda-like polypeptide 1 | ↑ | 1.92 | 8.12E-04 |
| P14174 | Macrophage migration inhibitory factor | ↑ | 1.92 | 1.26E-02 |
| B8ZZ73 | Interleukin 1 receptor type 1 | ↑ | 1.93 | 5.50E-04 |
| O94919 | Endonuclease domain-containing 1 protein | ↑ | 1.93 | 1.27E-02 |
| Q8TBP5 | Membrane protein FAM174A | ↑ | 1.93 | 4.99E-02 |
| Q9UHR4 | Brain-specific angiogenesis inhibitor 1-associated protein 2-like protein 1 | ↑ | 1.94 | 8.90E-03 |
| A0A0A0MS15 | Immunoglobulin heavy variable 3-49 | ↑ | 1.94 | 8.76E-04 |
| P31151 | Protein S100-A7 | ↑ | 1.95 | 1.90E-02 |
| O60279 | Sushi domain-containing protein 5 | ↑ | 1.96 | 1.40E-02 |
| A0A075B6R2 | Immunoglobulin heavy variable 4-4 | ↑ | 1.96 | 1.57E-03 |
| J3KPQ0 | Fibroblast growth factor receptor | ↑ | 1.97 | 8.14E-03 |
| P23083 | Immunoglobulin heavy variable 1-2 | ↑ | 1.98 | 6.43E-04 |
| J3QRS3 | Myosin light chain 12A | ↑ | 1.98 | 6.59E-04 |
| Q92626 | Peroxidasin homolog | ↑ | 1.98 | 4.88E-02 |
| Q92496 | Complement factor H-related protein 4 | ↑ | 1.99 | 6.92E-03 |
| P02760 | Protein AMBP | ↑ | 2.01 | 1.16E-02 |
| Q14315 | Filamin-C | ↑ | 2.01 | 6.51E-03 |
| Q14118 | Dystroglycan 1 | ↑ | 2.01 | 2.27E-02 |
| A0A0B4J2B5 | Immunoglobulin heavy variable 3/OR16-9 | ↑ | 2.01 | 9.88E-03 |
| O75368 | Adapter SH3BGRL | ↑ | 2.02 | 3.06E-03 |
| O00187 | Mannan-binding lectin serine protease 2 | ↑ | 2.03 | 2.99E-02 |
| A0A0J9YY99 | Ig-like domain-containing protein | ↑ | 2.03 | 6.26E-04 |
| A0A087X0H5 | Growth hormone receptor | ↑ | 2.03 | 4.41E-03 |
| P06396 | Gelsolin | ↑ | 2.04 | 3.25E-02 |
| A0A0A0MR25 | Fibroblast growth factor receptor | ↑ | 2.04 | 6.12E-03 |
| Q9UFM8 | Neuroplastin | ↑ | 2.04 | 2.87E-02 |
| Q9H461 | Frizzled-8 | ↑ | 2.05 | 1.02E-04 |
| P48745 | CCN family member 3 | ↑ | 2.07 | 3.41E-03 |
| P55000 | Secreted Ly-6/uPAR-related protein 1 | ↑ | 2.08 | 1.10E-04 |
| Q9H1E1 | Ribonuclease 7 | ↑ | 2.08 | 1.81E-02 |
| Q9GZM7 | Tubulointerstitial nephritis antigen-like | ↑ | 2.08 | 3.43E-04 |
| P02765 | Alpha-2-HS-glycoprotein | ↑ | 2.08 | 8.41E-03 |
| P32942 | Intercellular adhesion molecule 3 | ↑ | 2.09 | 9.22E-05 |
| E7EMR3 | Adhesion G protein-coupled receptor L3 | ↑ | 2.09 | 1.88E-02 |
| P00918 | Carbonic anhydrase 2 | ↑ | 2.11 | 5.39E-03 |
| P81605 | Dermcidin | ↑ | 2.12 | 5.81E-03 |
| P02452 | Collagen alpha-1 | ↑ | 2.13 | 2.48E-03 |
| H7C3P2 | Collagen type XXVIII alpha 1 chain | ↑ | 2.13 | 2.36E-03 |
| Q6ZNA5 | Ferric-chelate reductase 1 | ↑ | 2.14 | 3.44E-02 |
| I3L0L6 | E3 ubiquitin-protein ligase RNF167 | ↑ | 2.15 | 2.90E-02 |
| Q15375 | Ephrin type-A receptor 7 | ↑ | 2.16 | 5.28E-04 |
| Q13790 | Apolipoprotein F | ↑ | 2.19 | 1.34E-03 |
| K3W4U1 | Fc epsilon receptor II | ↑ | 2.20 | 1.11E-03 |
| P04196 | Histidine-rich glycoprotein | ↑ | 2.21 | 6.77E-03 |
| P07437 | Tubulin beta chain | ↑ | 2.21 | 8.04E-03 |
| O43570 | Carbonic anhydrase 12 | ↑ | 2.22 | 1.10E-05 |
| Q9ULV1 | Frizzled-4 | ↑ | 2.23 | 1.56E-02 |
| Q15828 | Cystatin-M | ↑ | 2.23 | 3.37E-02 |
| Q5T123 | SH3 domain-binding glutamic acid-rich-like protein 3 | ↑ | 2.25 | 1.46E-02 |
| O94760 | Dimethylarginine dimethylaminohydrolase 1 | ↑ | 2.25 | 2.80E-03 |
| O00401 | Actin nucleation-promoting factor WASL | ↑ | 2.27 | 3.31E-02 |
| Q96IU4 | Putative protein-lysine deacylase ABHD14B | ↑ | 2.27 | 1.07E-04 |
| P29692 | Elongation factor 1-delta | ↑ | 2.28 | 3.18E-03 |
| P83110 | Serine protease HTRA3 | ↑ | 2.30 | 7.00E-04 |
| A0A087WX80 | Laminin subunit alpha 2 | ↑ | 2.30 | 2.40E-03 |
| P01714 | Immunoglobulin lambda variable 3-19 | ↑ | 2.31 | 1.88E-02 |
| Q5KU26 | Collectin-12 | ↑ | 2.31 | 4.58E-02 |
| P0DOY3 | Immunoglobulin lambda constant 3 | ↑ | 2.31 | 1.52E-02 |
| F6SYF8 | Dickkopf WNT signaling pathway inhibitor 3 | ↑ | 2.32 | 1.57E-03 |
| P13647 | Keratin, type II cytoskeletal 5 | ↑ | 2.33 | 3.15E-02 |
| Q6PI73 | Leukocyte immunoglobulin-like receptor subfamily A member 6 | ↑ | 2.33 | 2.48E-02 |
| Q99983 | Osteomodulin | ↑ | 2.34 | 3.81E-02 |
| P22304 | Iduronate 2-sulfatase | ↑ | 2.35 | 1.45E-02 |
| P10109 | Adrenodoxin, mitochondrial | ↑ | 2.35 | 1.48E-02 |
| Q9Y5Z4 | Heme-binding protein 2 | ↑ | 2.37 | 3.18E-02 |
| O95388 | CCN family member 4 | ↑ | 2.38 | 1.59E-03 |
| Q96S96 | Phosphatidylethanolamine-binding protein 4 | ↑ | 2.40 | 1.35E-02 |
| O95971 | CD160 antigen | ↑ | 2.40 | 3.18E-03 |
| P04430 | Immunoglobulin kappa variable 1-16 | ↑ | 2.41 | 3.33E-02 |
| O95841 | Angiopoietin-related protein 1 | ↑ | 2.41 | 2.50E-03 |
| P29401 | Transketolase | ↑ | 2.44 | 2.98E-02 |
| P19823 | Inter-alpha-trypsin inhibitor heavy chain H2 | ↑ | 2.45 | 2.81E-02 |
| P55259 | Pancreatic secretory granule membrane major glycoprotein GP2 | ↑ | 2.46 | 1.18E-02 |
| A0A0C4DH43 | Immunoglobulin heavy variable 2-70D | ↑ | 2.51 | 1.65E-03 |
| Q8NFT8 | Delta and Notch-like epidermal growth factor-related receptor | ↑ | 2.52 | 8.38E-04 |
| Q9NY37 | Acid-sensing ion channel 5 | ↑ | 2.53 | 6.05E-03 |
| O15145 | Actin-related protein 2/3 complex subunit 3 | ↑ | 2.54 | 1.20E-02 |
| P16989 | Y-box-binding protein 3 | ↑ | 2.54 | 7.16E-03 |
| A0A3B3IRL2 | Cellular repressor of E1A stimulated genes 1 | ↑ | 2.55 | 1.81E-02 |
| A0A286YFJ8 | Immunoglobulin heavy constant gamma 4 | ↑ | 2.55 | 1.72E-02 |
| P04155 | Trefoil factor 1 | ↑ | 2.57 | 9.53E-03 |
| Q16348 | Solute carrier family 15 member 2 | ↑ | 2.58 | 4.87E-02 |
| F8W1A4 | Adenylate kinase 2, mitochondrial | ↑ | 2.59 | 2.60E-02 |
| Q92765 | Secreted frizzled-related protein 3 | ↑ | 2.59 | 5.62E-03 |
| Q9HB75 | p53-induced death domain-containing protein 1 | ↑ | 2.60 | 1.47E-03 |
| P00491 | Purine nucleoside phosphorylase | ↑ | 2.61 | 4.26E-02 |
| P32320 | Cytidine deaminase | ↑ | 2.62 | 2.22E-02 |
| Q9H741 | SREBP regulating gene protein | ↑ | 2.65 | 2.14E-02 |
| B1AH90 | Signal peptide, CUB domain and EGF like domain containing 1 | ↑ | 2.67 | 2.19E-02 |
| P11766 | Alcohol dehydrogenase class-3 | ↑ | 2.69 | 1.71E-04 |
| P32004 | Neural cell adhesion molecule L1 | ↑ | 2.70 | 8.43E-03 |
| P01031 | Complement C5 | ↑ | 2.71 | 4.92E-02 |
| O43852 | Calumenin | ↑ | 2.72 | 2.52E-03 |
| O94985 | Calsyntenin-1 | ↑ | 2.73 | 9.80E-03 |
| Q9UKY0 | Prion-like protein doppel | ↑ | 2.74 | 7.99E-03 |
| P30046 | D-dopachrome decarboxylase | ↑ | 2.75 | 1.76E-02 |
| Q96A22 | Uncharacterized protein C11orf52 | ↑ | 2.76 | 1.90E-02 |
| P36941 | Tumor necrosis factor receptor superfamily member 3 | ↑ | 2.77 | 4.47E-02 |
| P04179 | Superoxide dismutase [Mn], mitochondrial | ↑ | 2.79 | 4.54E-02 |
| O75487 | Glypican-4 | ↑ | 2.79 | 2.36E-02 |
| Q9Y6U3 | Scinderin | ↑ | 2.81 | 9.36E-03 |
| P13489 | Ribonuclease inhibitor | ↑ | 2.83 | 2.03E-02 |
| P30044 | Peroxiredoxin-5, mitochondrial | ↑ | 2.86 | 5.29E-03 |
| Q16661 | Guanylate cyclase activator 2B | ↑ | 2.89 | 9.09E-03 |
| Q96I82 | Kazal-type serine protease inhibitor domain-containing protein 1 | ↑ | 2.90 | 4.99E-03 |
| P13861 | cAMP-dependent protein kinase type II-alpha regulatory subunit | ↑ | 2.91 | 3.77E-02 |
| F5H1S8 | Malectin | ↑ | 2.91 | 6.11E-03 |
| Q15833 | Syntaxin-binding protein 2 | ↑ | 2.94 | 1.32E-02 |
| P30475 | HLA class I histocompatibility antigen, B alpha chain | ↑ | 2.95 | 2.11E-03 |
| E9PL83 | Pro-adrenomedullin | ↑ | 2.99 | 1.84E-02 |
| O95633 | Follistatin-related protein 3 | ↑ | 3.01 | 1.81E-02 |
| A0A0C4DH72 | Immunoglobulin kappa variable 1-6 | ↑ | 3.03 | 2.39E-03 |
| P25391 | Laminin subunit alpha-1 | ↑ | 3.04 | 5.06E-03 |
| P18669 | Phosphoglycerate mutase 1 | ↑ | 3.11 | 3.15E-02 |
| Q9Y281 | Cofilin-2 | ↑ | 3.11 | 2.29E-02 |
| P28066 | Proteasome subunit alpha type-5 | ↑ | 3.11 | 3.40E-02 |
| Q9UJ72 | Annexin A10 | ↑ | 3.15 | 7.77E-03 |
| P05556 | Integrin beta-1 | ↑ | 3.17 | 2.14E-04 |
| C9JM33 | Interferon alpha and beta receptor subunit 2 | ↑ | 3.21 | 1.55E-02 |
| K7ELM9 | Apolipoprotein C1 | ↑ | 3.22 | 1.24E-03 |
| A0A1B0GTG2 | Aldehyde dehydrogenase 7 family member A1 | ↑ | 3.27 | 1.09E-02 |
| P27348 | 14-3-3 protein theta | ↑ | 3.28 | 5.50E-03 |
| P17655 | Calpain-2 catalytic subunit | ↑ | 3.30 | 2.52E-04 |
| Q5VY43 | Platelet endothelial aggregation receptor 1 | ↑ | 3.32 | 4.49E-02 |
| Q9H0E2 | Toll-interacting protein | ↑ | 3.37 | 3.77E-02 |
| P05186 | Alkaline phosphatase, tissue-nonspecific isozyme | ↑ | 3.40 | 2.10E-02 |
| Q9BY89 | Uncharacterized protein KIAA1671 | ↑ | 3.41 | 1.80E-02 |
| O00144 | Frizzled-9 | ↑ | 3.43 | 1.07E-02 |
| O95980 | Reversion-inducing cysteine-rich protein with Kazal motifs | ↑ | 3.44 | 6.50E-03 |
| A0A0C4DGB5 | Calpastatin | ↑ | 3.48 | 2.72E-02 |
| P00738 | Haptoglobin | ↑ | 3.57 | 4.67E-02 |
| P80748 | Immunoglobulin lambda variable 3-21 | ↑ | 3.59 | 3.92E-02 |
| G5E9G7 | Neurexin 2 | ↑ | 3.61 | 7.17E-05 |
| P00748 | Coagulation factor XII | ↑ | 3.63 | 2.10E-02 |
| P12931 | Proto-oncogene tyrosine-protein kinase Src | ↑ | 3.66 | 6.34E-05 |
| P22692 | Insulin-like growth factor-binding protein 4 | ↑ | 3.68 | 4.02E-02 |
| Q14142 | Tripartite motif-containing protein 14 | ↑ | 3.68 | 3.87E-02 |
| P28072 | Proteasome subunit beta type-6 | ↑ | 3.70 | 2.00E-02 |
| G3V4P8 | Glia maturation factor beta | ↑ | 3.71 | 4.55E-02 |
| E7EX17 | Eukaryotic translation initiation factor 4B | ↑ | 3.73 | 4.97E-02 |
| Q9UJ99 | Cadherin-22 | ↑ | 3.73 | 1.18E-02 |
| O75891 | Cytosolic 10-formyltetrahydrofolate dehydrogenase | ↑ | 3.74 | 2.44E-02 |
| Q5T2W1 | Na(+)/H(+) exchange regulatory cofactor NHE-RF3 | ↑ | 3.78 | 1.38E-02 |
| D6RHW5 | Endomucin | ↑ | 3.79 | 4.55E-02 |
| Q3LXA3 | Triokinase/FMN cyclase | ↑ | 3.82 | 3.39E-02 |
| P20774 | Mimecan | ↑ | 3.83 | 3.33E-02 |
| Q92484 | Acid sphingomyelinase-like phosphodiesterase 3a | ↑ | 3.85 | 3.90E-03 |
| A0A0A0MSA0 | Laminin subunit alpha-3 | ↑ | 3.90 | 4.37E-02 |
| O95817 | BAG family molecular chaperone regulator 3 | ↑ | 3.91 | 3.39E-02 |
| Q7Z794 | Keratin, type II cytoskeletal 1b | ↑ | 3.93 | 2.02E-03 |
| A0A0A0MSS8 | Aldo-keto reductase family 1 member C3 | ↑ | 3.95 | 2.30E-02 |
| Q8IW52 | SLIT and NTRK-like protein 4 | ↑ | 3.95 | 4.75E-02 |
| O95171 | Sciellin | ↑ | 3.98 | 3.82E-02 |
| Q8WWA0 | Intelectin-1 | ↑ | 4.21 | 1.30E-02 |
| A0A0C4DGE4 | Syntaxin 3 | ↑ | 4.25 | 2.37E-02 |
| Q8N474 | Secreted frizzled-related protein 1 | ↑ | 4.25 | 3.63E-02 |
| P02042 | Hemoglobin subunit delta | ↑ | 4.27 | 6.34E-03 |
| Q8WWY7 | WAP four-disulfide core domain protein 12 | ↑ | 4.32 | 2.58E-02 |
| P09486 | Secreted protein acidic and rich in cysteine | ↑ | 4.32 | 3.80E-02 |
| Q14050 | Collagen alpha-3 | ↑ | 4.35 | 2.01E-02 |
| P25788 | Proteasome subunit alpha type-3 | ↑ | 4.39 | 4.12E-02 |
| P35579 | Myosin-9 | ↑ | 4.48 | 2.73E-02 |
| P29992 | Guanine nucleotide-binding protein subunit alpha-11 | ↑ | 4.53 | 2.02E-02 |
| P05026 | Sodium/potassium-transporting ATPase subunit beta-1 | ↑ | 4.54 | 4.45E-03 |
| P00740 | Coagulation factor IX | ↑ | 4.55 | 2.15E-02 |
| A8MVW5 | HEPACAM family member 2 | ↑ | 4.57 | 4.69E-02 |
| A8K878 | cDNA FLJ77177, highly similar to Homo sapiens arginine-rich, mutated in early stage tumors | ↑ | 4.59 | 6.41E-03 |
| H3BTI0 | Cysteine rich secretory protein LCCL domain containing 2 | ↑ | 4.60 | 2.81E-02 |
| A0A1W2PNV4 | Actin-related protein 2/3 complex subunit 1A | ↑ | 4.61 | 3.35E-02 |
| Q13103 | Secreted phosphoprotein 24 | ↑ | 4.63 | 2.33E-02 |
| P63096 | Guanine nucleotide-binding protein G | ↑ | 4.68 | 3.84E-02 |
| P00390 | Glutathione reductase, mitochondrial | ↑ | 4.68 | 3.76E-02 |
| Q8WWZ8 | Oncoprotein-induced transcript 3 protein | ↑ | 4.68 | 4.86E-02 |
| Q9BY43 | Charged multivesicular body protein 4a | ↑ | 4.68 | 1.94E-02 |
| P52961 | GPI-linked NAD | ↑ | 4.71 | 3.65E-03 |
| P05023 | Sodium/potassium-transporting ATPase subunit alpha-1 | ↑ | 4.73 | 4.20E-02 |
| A0A0C4DH07 | Latent transforming growth factor beta binding protein 4 | ↑ | 4.76 | 1.18E-03 |
| Q14697 | Neutral alpha-glucosidase AB | ↑ | 4.93 | 4.60E-02 |
| Q8WWQ8 | Stabilin-2 | ↑ | 4.95 | 1.99E-02 |
| A0A0D9SG04 | Cordon-bleu WH2 repeat protein like 1 | ↑ | 5.01 | 9.28E-03 |
| P36543 | V-type proton ATPase subunit E 1 | ↑ | 5.05 | 4.18E-02 |
| P26641 | Elongation factor 1-gamma | ↑ | 5.05 | 1.46E-02 |
| P37837 | Transaldolase | ↑ | 5.13 | 3.57E-02 |
| P21399 | Cytoplasmic aconitate hydratase | ↑ | 5.15 | 4.07E-02 |
| A0A075B6K4 | Immunoglobulin lambda variable 3-10 | ↑ | 5.20 | 1.70E-02 |
| Q9BTY2 | Plasma alpha-L-fucosidase | ↑ | 5.20 | 4.69E-02 |
| Q99584 | Protein S100-A13 | ↑ | 5.23 | 3.80E-02 |
| Q9UHI8 | A disintegrin and metalloproteinase with thrombospondin motifs 1 | ↑ | 5.30 | 3.33E-03 |
| Q86UN2 | Reticulon-4 receptor-like 1 | ↑ | 5.40 | 1.69E-02 |
| P06858 | Lipoprotein lipase | ↑ | 5.40 | 1.06E-03 |
| Q9NZ08 | Endoplasmic reticulum aminopeptidase 1 | ↑ | 5.42 | 9.08E-03 |
| P50502 | Hsc70-interacting protein | ↑ | 5.43 | 2.60E-02 |
| Q9P1F3 | Costars family protein ABRACL | ↑ | 5.47 | 4.36E-02 |
| Q10471 | Polypeptide N-acetylgalactosaminyltransferase 2 | ↑ | 5.48 | 4.89E-02 |
| P07357 | Complement component C8 alpha chain | ↑ | 5.48 | 9.60E-03 |
| Q9UGB7 | Inositol oxygenase | ↑ | 5.52 | 4.03E-02 |
| P31327 | Carbamoyl-phosphate synthase [ammonia], mitochondrial | ↑ | 5.52 | 4.10E-02 |
| B1AK87 | F-actin-capping protein subunit beta | ↑ | 5.56 | 3.08E-02 |
| A6NKB8 | Arginyl aminopeptidase | ↑ | 5.57 | 1.34E-02 |
| O60449 | Lymphocyte antigen 75 | ↑ | 5.57 | 6.72E-03 |
| P46940 | Ras GTPase-activating-like protein IQGAP1 | ↑ | 5.58 | 4.44E-02 |
| P06748 | Nucleophosmin | ↑ | 5.62 | 3.73E-02 |
| P48740 | Mannan-binding lectin serine protease 1 | ↑ | 5.73 | 4.87E-02 |
| P68036 | Ubiquitin-conjugating enzyme E2 L3 | ↑ | 5.73 | 4.69E-02 |
| Q9BQR3 | Serine protease 27 | ↑ | 5.73 | 3.55E-02 |
| Q01518 | Adenylyl cyclase-associated protein 1 | ↑ | 5.74 | 4.99E-02 |
| A0A0J9YWL0 | Crystallin beta-gamma domain containing 1 | ↑ | 5.75 | 1.75E-02 |
| P24752 | Acetyl-CoA acetyltransferase, mitochondrial | ↑ | 5.78 | 2.50E-02 |
| Q15848 | Adiponectin | ↑ | 5.80 | 9.38E-03 |
| Q86XT2 | Vacuolar protein sorting-associated protein 37D | ↑ | 5.87 | 5.40E-03 |
| P98161 | Polycystin-1 | ↑ | 5.96 | 4.42E-02 |
| P50395 | Rab GDP dissociation inhibitor beta | ↑ | 5.96 | 2.79E-03 |
| P31371 | Fibroblast growth factor 9 | ↑ | 5.96 | 2.89E-02 |
| A0A024R571 | EH domain containing 1 | ↑ | 6.06 | 6.49E-03 |
| Q14240 | Eukaryotic initiation factor 4A-II | ↑ | 6.07 | 1.24E-02 |
| O00764 | Pyridoxal kinase | ↑ | 6.09 | 4.44E-02 |
| P24666 | Low molecular weight phosphotyrosine protein phosphatase | ↑ | 6.14 | 4.84E-02 |
| Q9H9H4 | Vacuolar protein sorting-associated protein 37B | ↑ | 6.16 | 4.15E-02 |
| P61978 | Heterogeneous nuclear ribonucleoprotein K | ↑ | 6.16 | 2.48E-02 |
| A0A0G2JLS4 | Leukocyte immunoglobulin-like receptor subfamily B member 1 | ↑ | 6.21 | 9.97E-03 |
| P55072 | Transitional endoplasmic reticulum ATPase | ↑ | 6.21 | 2.01E-02 |
| Q93099 | Homogentisate 1,2-dioxygenase | ↑ | 6.24 | 4.04E-02 |
| A0A0C4DFN5 | Chromosome 10 open reading frame 61 | ↑ | 6.26 | 4.54E-02 |
| P22894 | Neutrophil collagenase | ↑ | 6.29 | 1.53E-02 |
| Q9H223 | EH domain-containing protein 4 | ↑ | 6.37 | 3.13E-02 |
| Q9Y5H8 | Protocadherin alpha-3 | ↑ | 6.47 | 4.48E-02 |
| P01817 | Immunoglobulin heavy variable 2-5 | ↑ | 6.49 | 4.51E-02 |
| G3V180 | Dipeptidyl peptidase 3 | ↑ | 6.53 | 2.02E-02 |
| Q86VZ4 | Low-density lipoprotein receptor-related protein 11 | ↑ | 6.58 | 2.09E-02 |
| P22314 | Ubiquitin-like modifier-activating enzyme 1 | ↑ | 6.59 | 3.05E-02 |
| Q9NRA1 | Platelet-derived growth factor C | ↑ | 6.59 | 4.42E-02 |
| Q6UXN8 | C-type lectin domain family 9 member A | ↑ | 6.72 | 2.36E-02 |
| A0A3B3IQ51 | Complement factor H related 2 | ↑ | 6.76 | 1.96E-02 |
| Q9NP85 | Podocin | ↑ | 6.78 | 2.12E-02 |
| B5MBX2 | Transcobalamin-2 | ↑ | 6.81 | 4.76E-02 |
| P62070 | Ras-related protein R-Ras2 | ↑ | 6.88 | 3.30E-02 |
| O75354 | Ectonucleoside triphosphate diphosphohydrolase 6 | ↑ | 6.94 | 3.80E-02 |
| H3BMA1 | Mesothelin | ↑ | 6.97 | 2.77E-03 |
| A0A087WSY5 | Carboxypeptidase B2 | ↑ | 7.12 | 2.93E-02 |
| Q9UHY7 | Enolase-phosphatase E1 | ↑ | 7.13 | 3.08E-02 |
| Q15181 | Inorganic pyrophosphatase | ↑ | 7.17 | 8.95E-03 |
| Q9Y5K6 | CD2-associated protein | ↑ | 7.18 | 2.91E-02 |
| O00461 | Golgi integral membrane protein 4 | ↑ | 7.21 | 1.74E-02 |
| P61020 | Ras-related protein Rab-5B | ↑ | 7.26 | 1.33E-02 |
| Q92890 | Ubiquitin recognition factor in ER-associated degradation protein 1 | ↑ | 7.28 | 4.85E-02 |
| P31150 | Rab GDP dissociation inhibitor alpha | ↑ | 7.31 | 2.70E-02 |
| Q13867 | Bleomycin hydrolase | ↑ | 7.39 | 1.67E-02 |
| Q12765 | Secernin-1 | ↑ | 7.41 | 1.46E-02 |
| P40197 | Platelet glycoprotein V | ↑ | 7.45 | 7.87E-03 |
| Q9UFP1 | Golgi-associated kinase 1A | ↑ | 7.46 | 2.78E-02 |
| M0QYN0 | Myeloid derived growth factor | ↑ | 7.49 | 2.18E-03 |
| P28332 | Alcohol dehydrogenase 6 | ↑ | 7.51 | 3.16E-02 |
| P21964 | Catechol O-methyltransferase | ↑ | 7.53 | 1.92E-02 |
| P21583 | Kit ligand | ↑ | 7.54 | 2.35E-02 |
| E9PGC5 | protein-tyrosine-phosphatase | ↑ | 7.56 | 2.58E-03 |
| Q6ZMJ2 | Scavenger receptor class A member 5 | ↑ | 7.59 | 1.55E-02 |
| Q86TD4 | Sarcalumenin | ↑ | 7.65 | 4.96E-02 |
| Q96KN2 | Beta-Ala-His dipeptidase | ↑ | 7.71 | 1.85E-02 |
| P00742 | Coagulation factor X | ↑ | 7.82 | 5.44E-03 |
| A0A075B788 | Protein tyrosine phosphatase receptor type C | ↑ | 7.89 | 1.08E-02 |
| A0A0C4DGN2 | Sex hormone-binding globulin | ↑ | 7.89 | 2.79E-02 |
| F5H4M7 | Transmembrane p24 trafficking protein 3 | ↑ | 7.95 | 3.08E-02 |
| B5MCA4 | Epithelial cell adhesion molecule | ↑ | 8.03 | 3.75E-02 |
| C9J0J7 | Profilin | ↑ | 8.07 | 2.19E-02 |
| Q99795 | Cell surface A33 antigen | ↑ | 8.14 | 5.63E-03 |
| B7ZM79 | PCDH9 protein | ↑ | 8.22 | 2.08E-02 |
| P00533 | Epidermal growth factor receptor | ↑ | 8.22 | 3.40E-02 |
| Q14353 | Guanidinoacetate N-methyltransferase | ↑ | 8.23 | 5.25E-03 |
| Q5VT99 | Leucine-rich repeat-containing protein 38 | ↑ | 8.27 | 2.86E-02 |
| P00492 | Hypoxanthine-guanine phosphoribosyltransferase | ↑ | 8.28 | 4.73E-03 |
| P07384 | Calpain-1 catalytic subunit | ↑ | 8.44 | 2.15E-02 |
| O43396 | Thioredoxin-like protein 1 | ↑ | 8.50 | 2.21E-02 |
| P09525 | Annexin A4 | ↑ | 8.55 | 4.63E-03 |
| P40227 | T-complex protein 1 subunit zeta | ↑ | 8.58 | 1.79E-02 |
| B0YIW2 | Apolipoprotein C-III | ↑ | 8.60 | 9.29E-03 |
| Q15366 | Poly(rC)-binding protein 2 | ↑ | 8.73 | 1.24E-02 |
| Q6H9L7 | Isthmin-2 | ↑ | 8.78 | 2.75E-02 |
| P06576 | ATP synthase subunit beta, mitochondrial | ↑ | 8.84 | 1.67E-02 |
| Q6P4A8 | Phospholipase B-like 1 | ↑ | 8.89 | 7.76E-03 |
| B8ZZ19 | Parvalbumin | ↑ | 9.06 | 4.36E-02 |
| A0A0U1RQV3 | EGF containing fibulin extracellular matrix protein 1 | ↑ | 9.07 | 2.09E-02 |
| D6RAF8 | Heterogeneous nuclear ribonucleoprotein D | ↑ | 9.12 | 4.32E-02 |
| P31948 | Stress-induced-phosphoprotein 1 | ↑ | 9.16 | 1.17E-02 |
| Q9BR76 | Coronin-1B | ↑ | 9.18 | 3.36E-02 |
| Q9UJJ9 | N-acetylglucosamine-1-phosphotransferase subunit gamma | ↑ | 9.68 | 2.00E-02 |
| Q9HCH3 | Copine-5 | ↑ | 9.70 | 2.87E-02 |
| Q9UN75 | Protocadherin alpha-12 | ↑ | 9.73 | 3.30E-02 |
| P32189 | Glycerol kinase | ↑ | 10.02 | 1.33E-02 |
| P29373 | Cellular retinoic acid-binding protein 2 | ↑ | 10.22 | 4.12E-03 |
| Q96BW5 | Phosphotriesterase-related protein | ↑ | 10.85 | 2.76E-03 |
| F8WCF6 | Actin-related protein 2/3 complex subunit 4 | ↑ | 10.87 | 2.82E-02 |
| P37235 | Hippocalcin-like protein 1 | ↑ | 10.94 | 2.58E-02 |
| G3XAF7 | Brain enriched myelin associated protein 1 | ↑ | 10.96 | 3.37E-02 |
| A0A096LPE2 | SAA2-SAA4 readthrough | ↑ | 10.97 | 3.60E-03 |
| Q15149 | Plectin | ↑ | 11.07 | 2.35E-03 |
| C9JBI3 | Phosphoserine phosphatase | ↑ | 11.20 | 1.57E-02 |
| Q9NYQ8 | Protocadherin Fat 2 | ↑ | 11.27 | 3.17E-02 |
| O60784 | Target of Myb1 membrane trafficking protein | ↑ | 11.30 | 1.39E-02 |
| E9PIT4 | Transmembrane protein 25 | ↑ | 11.41 | 3.05E-02 |
| P24298 | Alanine aminotransferase 1 | ↑ | 11.45 | 3.25E-02 |
| Q9UJU6 | Drebrin-like protein | ↑ | 11.66 | 3.69E-02 |
| Q6P1N0 | Coiled-coil and C2 domain-containing protein 1A | ↑ | 11.67 | 4.97E-02 |
| P54652 | Heat shock-related 70 kDa protein 2 | ↑ | 11.70 | 1.12E-02 |
| A6NFX8 | Nudix hydrolase 5 | ↑ | 11.92 | 2.30E-02 |
| P18428 | Lipopolysaccharide-binding protein | ↑ | 12.15 | 4.31E-02 |
| Q9BW04 | Specifically androgen-regulated gene protein | ↑ | 12.15 | 4.45E-02 |
| O95154 | Aflatoxin B1 aldehyde reductase member 3 | ↑ | 12.23 | 1.07E-02 |
| P51149 | Ras-related protein Rab-7a | ↑ | 12.62 | 5.25E-03 |
| P17405 | Sphingomyelin phosphodiesterase | ↑ | 12.96 | 2.45E-02 |
| Q8N8Z6 | Discoidin, CUB and LCCL domain-containing protein 1 | ↑ | 13.47 | 2.51E-02 |
| Q9BXJ7 | Protein amnionless | ↑ | 13.84 | 1.06E-03 |
| O14732 | Inositol monophosphatase 2 | ↑ | 14.05 | 3.00E-02 |
| Q09328 | Alpha-1,6-mannosylglycoprotein 6-beta-N-acetylglucosaminyltransferase A | ↑ | 14.27 | 7.63E-03 |
| C9JL73 | Vacuolar proton pump subunit B | ↑ | 14.30 | 4.80E-02 |
| Q04917 | 14-3-3 protein eta | ↑ | 14.34 | 1.07E-02 |
| B4DUR8 | T-complex protein 1 subunit gamma | ↑ | 14.64 | 3.46E-02 |
| Q9Y490 | Talin-1 | ↑ | 14.73 | 8.81E-03 |
| Q96HC4 | PDZ and LIM domain protein 5 | ↑ | 14.78 | 1.41E-02 |
| P16562 | Cysteine-rich secretory protein 2 | ↑ | 15.01 | 6.75E-03 |
| A0A0B4J2C3 | Translationally-controlled tumor protein | ↑ | 15.06 | 1.39E-02 |
| O75264 | Small integral membrane protein 24 | ↑ | 15.84 | 1.16E-02 |
| H0Y4H3 | CD99 antigen-like protein 2 | ↑ | 16.01 | 9.45E-03 |
| A5D8V6 | Vacuolar protein sorting-associated protein 37C | ↑ | 16.07 | 6.63E-03 |
| Q9Y6X5 | Bis(5'-adenosyl)-triphosphatase ENPP4 | ↑ | 16.22 | 8.25E-03 |
| P28838 | Cytosol aminopeptidase | ↑ | 16.30 | 2.70E-02 |
| B7ZAR1 | T-complex protein 1 subunit epsilon | ↑ | 16.74 | 2.19E-02 |
| Q9HBJ8 | Collectrin | ↑ | 17.23 | 1.28E-02 |
| O14737 | Programmed cell death protein 5 | ↑ | 17.53 | 2.22E-02 |
| Q9BUP0 | EF-hand domain-containing protein D1 | ↑ | 17.57 | 1.59E-02 |
| P09382 | Galectin-1 | ↑ | 17.68 | 2.60E-02 |
| O00253 | Agouti-related protein | ↑ | 17.72 | 1.93E-02 |
| P04424 | Argininosuccinate lyase | ↑ | 17.99 | 1.66E-02 |
| E7END6 | Vitamin K-dependent protein C | ↑ | 18.30 | 1.25E-02 |
| O15144 | Actin-related protein 2/3 complex subunit 2 | ↑ | 18.60 | 2.32E-02 |
| P28908 | Tumor necrosis factor receptor superfamily member 8 | ↑ | 18.86 | 3.78E-02 |
| P49641 | Alpha-mannosidase 2x | ↑ | 19.83 | 3.50E-02 |
| E7END7 | Ras-related protein Rab-1A | ↑ | 20.71 | 2.39E-03 |
| Q92597 | Protein NDRG1 | ↑ | 21.13 | 1.40E-02 |
| P00568 | Adenylate kinase isoenzyme 1 | ↑ | 21.51 | 1.04E-02 |
| Q9H190 | Syntenin-2 | ↑ | 21.59 | 9.22E-03 |
| H7BY57 | Neurofascin | ↑ | 22.82 | 4.66E-05 |
| K7EIK7 | EMAP like 2 | ↑ | 24.28 | 2.87E-03 |
| P45974 | Ubiquitin carboxyl-terminal hydrolase 5 | ↑ | 27.83 | 1.17E-02 |

TableS6 Differential protein in depressed patients compared to bipolar patients (FC≥2 or ≤0.5, P<0.01)

| **Accession** | **Protein names** | **Trend** | **FC** | **P-Value** |
| --- | --- | --- | --- | --- |
| Q53TN4 | Plasma membrane ascorbate-dependent reductase CYBRD1 | ↓ | 0.48 | 7.17E-03 |
| F8VNT9 | CD63 molecule | ↓ | 0.48 | 5.11E-03 |
| Q9H1C7 | Cysteine-rich and transmembrane domain-containing protein 1 | ↓ | 0.48 | 4.29E-03 |
| Q14315 | Filamin-C | ↑ | 2.01 | 6.51E-03 |
| A0A0B4J2B5 | Immunoglobulin heavy variable 3/OR16-9 | ↑ | 2.01 | 9.88E-03 |
| O75368 | Adapter SH3BGRL | ↑ | 2.02 | 3.06E-03 |
| A0A0J9YY99 | Ig-like domain-containing protein | ↑ | 2.03 | 6.26E-04 |
| A0A087X0H5 | Growth hormone receptor | ↑ | 2.03 | 4.41E-03 |
| A0A0A0MR25 | Fibroblast growth factor receptor | ↑ | 2.04 | 6.12E-03 |
| Q9H461 | Frizzled-8 | ↑ | 2.05 | 1.02E-04 |
| P48745 | CCN family member 3 | ↑ | 2.07 | 3.41E-03 |
| P55000 | Secreted Ly-6/uPAR-related protein 1 | ↑ | 2.08 | 1.10E-04 |
| Q9GZM7 | Tubulointerstitial nephritis antigen-like | ↑ | 2.08 | 3.43E-04 |
| P02765 | Alpha-2-HS-glycoprotein | ↑ | 2.08 | 8.41E-03 |
| P32942 | Intercellular adhesion molecule 3 | ↑ | 2.09 | 9.22E-05 |
| P00918 | Carbonic anhydrase 2 | ↑ | 2.11 | 5.39E-03 |
| P81605 | Dermcidin | ↑ | 2.12 | 5.81E-03 |
| P02452 | Collagen alpha-1 | ↑ | 2.13 | 2.48E-03 |
| H7C3P2 | Collagen type XXVIII alpha 1 chain | ↑ | 2.13 | 2.36E-03 |
| Q15375 | Ephrin type-A receptor 7 | ↑ | 2.16 | 5.28E-04 |
| Q13790 | Apolipoprotein F | ↑ | 2.19 | 1.34E-03 |
| K3W4U1 | Fc epsilon receptor II | ↑ | 2.20 | 1.11E-03 |
| P04196 | Histidine-rich glycoprotein | ↑ | 2.21 | 6.77E-03 |
| P07437 | Tubulin beta chain | ↑ | 2.21 | 8.04E-03 |
| O43570 | Carbonic anhydrase 12 | ↑ | 2.22 | 1.10E-05 |
| O94760 | N(G)-dimethylarginine dimethylaminohydrolase 1 | ↑ | 2.25 | 2.80E-03 |
| Q96IU4 | Putative protein-lysine deacylase ABHD14B | ↑ | 2.27 | 1.07E-04 |
| P29692 | Elongation factor 1-delta | ↑ | 2.28 | 3.18E-03 |
| P83110 | Serine protease HTRA3 | ↑ | 2.30 | 7.00E-04 |
| A0A087WX80 | Laminin subunit alpha 2 | ↑ | 2.30 | 2.40E-03 |
| F6SYF8 | Dickkopf WNT signaling pathway inhibitor 3 | ↑ | 2.32 | 1.57E-03 |
| O95388 | CCN family member 4 | ↑ | 2.38 | 1.59E-03 |
| O95971 | CD160 antigen | ↑ | 2.40 | 3.18E-03 |
| O95841 | Angiopoietin-related protein 1 | ↑ | 2.41 | 2.50E-03 |
| A0A0C4DH43 | Immunoglobulin heavy variable 2-70D | ↑ | 2.51 | 1.65E-03 |
| Q8NFT8 | Delta and Notch-like epidermal growth factor-related receptor | ↑ | 2.52 | 8.38E-04 |
| Q9NY37 | Acid-sensing ion channel 5 | ↑ | 2.53 | 6.05E-03 |
| P16989 | Y-box-binding protein 3 | ↑ | 2.54 | 7.16E-03 |
| P04155 | Trefoil factor 1 | ↑ | 2.57 | 9.53E-03 |
| Q92765 | Secreted frizzled-related protein 3 | ↑ | 2.59 | 5.62E-03 |
| Q9HB75 | p53-induced death domain-containing protein 1 | ↑ | 2.60 | 1.47E-03 |
| P11766 | Alcohol dehydrogenase class-3 | ↑ | 2.69 | 1.71E-04 |
| P32004 | Neural cell adhesion molecule L1 | ↑ | 2.70 | 8.43E-03 |
| O43852 | Calumenin | ↑ | 2.72 | 2.52E-03 |
| O94985 | Calsyntenin-1 | ↑ | 2.73 | 9.80E-03 |
| Q9UKY0 | Prion-like protein doppel | ↑ | 2.74 | 7.99E-03 |
| Q9Y6U3 | Scinderin | ↑ | 2.81 | 9.36E-03 |
| P30044 | Peroxiredoxin-5, mitochondrial | ↑ | 2.86 | 5.29E-03 |
| Q16661 | Guanylate cyclase activator 2B [Cleaved into: Guanylate cyclase C-activating peptide 2 | ↑ | 2.89 | 9.09E-03 |
| Q96I82 | Kazal-type serine protease inhibitor domain-containing protein 1 | ↑ | 2.90 | 4.99E-03 |
| F5H1S8 | Malectin | ↑ | 2.91 | 6.11E-03 |
| P30475 | HLA class I histocompatibility antigen, B alpha chain | ↑ | 2.95 | 2.11E-03 |
| A0A0C4DH72 | Immunoglobulin kappa variable 1-6 | ↑ | 3.03 | 2.39E-03 |
| P25391 | Laminin subunit alpha-1 | ↑ | 3.04 | 5.06E-03 |
| Q9UJ72 | Annexin A10 | ↑ | 3.15 | 7.77E-03 |
| P05556 | Integrin beta-1 | ↑ | 3.17 | 2.14E-04 |
| K7ELM9 | Apolipoprotein C1 | ↑ | 3.22 | 1.24E-03 |
| P27348 | 14-3-3 protein theta | ↑ | 3.28 | 5.50E-03 |
| P17655 | Calpain-2 catalytic subunit | ↑ | 3.30 | 2.52E-04 |
| O95980 | Reversion-inducing cysteine-rich protein with Kazal motifs | ↑ | 3.44 | 6.50E-03 |
| G5E9G7 | Neurexin 2 | ↑ | 3.61 | 7.17E-05 |
| P12931 | Proto-oncogene tyrosine-protein kinase Src | ↑ | 3.66 | 6.34E-05 |
| Q92484 | Acid sphingomyelinase-like phosphodiesterase 3a | ↑ | 3.85 | 3.90E-03 |
| Q7Z794 | Keratin, type II cytoskeletal 1b | ↑ | 3.93 | 2.02E-03 |
| P02042 | Hemoglobin subunit delta | ↑ | 4.27 | 6.34E-03 |
| P05026 | Sodium/potassium-transporting ATPase subunit beta-1 | ↑ | 4.54 | 4.45E-03 |
| A8K878 | cDNA FLJ77177, highly similar to Homo sapiens arginine-rich, mutated in early stage tumors | ↑ | 4.59 | 6.41E-03 |
| P52961 | GPI-linked NAD | ↑ | 4.71 | 3.65E-03 |
| A0A0C4DH07 | Latent transforming growth factor beta binding protein 4 | ↑ | 4.76 | 1.18E-03 |
| A0A0D9SG04 | Cordon-bleu WH2 repeat protein like 1 | ↑ | 5.01 | 9.28E-03 |
| Q9UHI8 | A disintegrin and metalloproteinase with thrombospondin motifs 1 | ↑ | 5.30 | 3.33E-03 |
| P06858 | Lipoprotein lipase | ↑ | 5.40 | 1.06E-03 |
| Q9NZ08 | Endoplasmic reticulum aminopeptidase 1 | ↑ | 5.42 | 9.08E-03 |
| P07357 | Complement component C8 alpha chain | ↑ | 5.48 | 9.60E-03 |
| O60449 | Lymphocyte antigen 75 | ↑ | 5.57 | 6.72E-03 |
| Q15848 | Adiponectin | ↑ | 5.80 | 9.38E-03 |
| Q86XT2 | Vacuolar protein sorting-associated protein 37D | ↑ | 5.87 | 5.40E-03 |
| P50395 | Rab GDP dissociation inhibitor beta | ↑ | 5.96 | 2.79E-03 |
| A0A024R571 | EH domain containing 1 | ↑ | 6.06 | 6.49E-03 |
| A0A0G2JLS4 | Leukocyte immunoglobulin-like receptor subfamily B member 1 | ↑ | 6.21 | 9.97E-03 |
| H3BMA1 | Mesothelin | ↑ | 6.97 | 2.77E-03 |
| Q15181 | Inorganic pyrophosphatase | ↑ | 7.17 | 8.95E-03 |
| P40197 | Platelet glycoprotein V | ↑ | 7.45 | 7.87E-03 |
| M0QYN0 | Myeloid derived growth factor | ↑ | 7.49 | 2.18E-03 |
| E9PGC5 | protein-tyrosine-phosphatase | ↑ | 7.56 | 2.58E-03 |
| P00742 | Coagulation factor X | ↑ | 7.82 | 5.44E-03 |
| Q99795 | Cell surface A33 antigen | ↑ | 8.14 | 5.63E-03 |
| Q14353 | Guanidinoacetate N-methyltransferase | ↑ | 8.23 | 5.25E-03 |
| P00492 | Hypoxanthine-guanine phosphoribosyltransferase | ↑ | 8.28 | 4.73E-03 |
| P09525 | Annexin A4 | ↑ | 8.55 | 4.63E-03 |
| B0YIW2 | Apolipoprotein C-III | ↑ | 8.60 | 9.29E-03 |
| Q6P4A8 | Phospholipase B-like 1 | ↑ | 8.89 | 7.76E-03 |
| P29373 | Cellular retinoic acid-binding protein 2 | ↑ | 10.22 | 4.12E-03 |
| Q96BW5 | Phosphotriesterase-related protein | ↑ | 10.85 | 2.76E-03 |
| A0A096LPE2 | SAA2-SAA4 readthrough | ↑ | 10.97 | 3.60E-03 |
| Q15149 | Plectin | ↑ | 11.07 | 2.35E-03 |
| P51149 | Ras-related protein Rab-7a | ↑ | 12.62 | 5.25E-03 |
| Q9BXJ7 | Protein amnionless [Cleaved into: Soluble protein amnionless] | ↑ | 13.84 | 1.06E-03 |
| Q09328 | Alpha-1,6-mannosylglycoprotein 6-beta-N-acetylglucosaminyltransferase A | ↑ | 14.27 | 7.63E-03 |
| Q9Y490 | Talin-1 | ↑ | 14.73 | 8.81E-03 |
| P16562 | Cysteine-rich secretory protein 2 | ↑ | 15.01 | 6.75E-03 |
| H0Y4H3 | CD99 antigen-like protein 2 | ↑ | 16.01 | 9.45E-03 |
| A5D8V6 | Vacuolar protein sorting-associated protein 37C | ↑ | 16.07 | 6.63E-03 |
| Q9Y6X5 | Bis(5'-adenosyl)-triphosphatase ENPP4 | ↑ | 16.22 | 8.25E-03 |
| E7END7 | Ras-related protein Rab-1A | ↑ | 20.71 | 2.39E-03 |
| Q9H190 | Syntenin-2 | ↑ | 21.59 | 9.22E-03 |
| H7BY57 | Neurofascin | ↑ | 22.82 | 4.66E-05 |
| K7EIK7 | EMAP like 2 | ↑ | 24.28 | 2.87E-03 |

TableS7 Biological Processes and Signaling Pathways Enriched for Differential Proteins Produced in the Biphasic Group Compared to the Depressed Group Under Relaxed Conditions(P<0.05)

| **BP** | **P-Value** | **pathway** | **P-Value** |
| --- | --- | --- | --- |
| cell adhesion | 4.70E-13 | Complement and coagulation cascades | 2.70E-11 |
| complement activation, classical pathway | 1.20E-12 | Regulation of actin cytoskeleton | 1.40E-09 |
| homophilic cell adhesion via plasma membrane adhesion molecules | 9.30E-10 | Necroptosis | 1.00E-06 |
| proteolysis | 5.80E-09 | Systemic lupus erythematosus | 9.80E-06 |
| innate immune response | 7.40E-09 | Proteoglycans in cancer | 4.20E-05 |
| cell-cell adhesion | 4.00E-08 | Shigellosis | 1.70E-04 |
| phagocytosis, recognition | 4.50E-08 | Endocytosis | 2.20E-04 |
| phagocytosis, engulfment | 5.40E-08 | Adherens junction | 9.00E-04 |
| positive regulation of B cell activation | 2.70E-07 | Amoebiasis | 1.80E-03 |
| positive regulation of canonical Wnt signaling pathway | 3.70E-07 | ECM-receptor interaction | 2.50E-03 |
| defense response to bacterium | 5.50E-07 | Focal adhesion | 2.70E-03 |
| adaptive immune response | 1.20E-06 | PI3K-Akt signaling pathway | 3.00E-03 |
| heterochromatin assembly | 1.80E-06 | Bacterial invasion of epithelial cells | 3.60E-03 |
| immunoglobulin mediated immune response | 2.20E-06 | Neutrophil extracellular trap formation | 4.10E-03 |
| B cell receptor signaling pathway | 2.60E-06 | Leukocyte transendothelial migration | 4.20E-03 |
| zymogen activation | 3.30E-06 | Cell adhesion molecules | 5.50E-03 |
| viral budding via host ESCRT complex | 1.60E-05 | Alcoholism | 8.90E-03 |
| immune response | 1.80E-05 | Rap1 signaling pathway | 9.20E-03 |
| non-canonical Wnt signaling pathway | 2.10E-05 | Tight junction | 9.70E-03 |
| adherens junction organization | 2.20E-05 | Biosynthesis of amino acids | 1.10E-02 |
| axon guidance | 3.80E-05 | Thiamine metabolism | 1.20E-02 |
| peptidyl-tyrosine phosphorylation | 7.90E-05 | Carbon metabolism | 1.30E-02 |
| blood coagulation | 1.20E-04 | Protein digestion and absorption | 2.00E-02 |
| multivesicular body assembly | 1.30E-04 | Pathogenic Escherichia coli infection | 2.90E-02 |
| blood coagulation, intrinsic pathway | 1.30E-04 | Tyrosine metabolism | 2.90E-02 |
| angiogenesis | 1.30E-04 | Mitophagy - animal | 3.00E-02 |
| signal transduction | 3.20E-04 | Staphylococcus aureus infection | 3.80E-02 |
| viral entry into host cell | 3.20E-04 | - | - |
| regulation of actin filament polymerization | 4.70E-04 | - | - |
| complement activation | 4.80E-04 | - | - |
| endocytosis | 5.40E-04 | - | - |
| synapse assembly | 6.00E-04 | - | - |
| cellular oxidant detoxification | 6.50E-04 | - | - |
| cell morphogenesis | 7.50E-04 | - | - |
| nucleobase-containing small molecule interconversion | 8.80E-04 | - | - |
| modification-dependent protein catabolic process | 8.80E-04 | - | - |
| ubiquitin-dependent protein catabolic process via the multivesicular body sorting pathway | 1.10E-03 | - | - |
| cell-cell junction assembly | 1.20E-03 | - | - |
| fibrinolysis | 1.30E-03 | - | - |
| calcium-dependent cell-cell adhesion via plasma membrane cell adhesion molecules | 1.40E-03 | - | - |
| ATP metabolic process | 1.50E-03 | - | - |
| antibacterial humoral response | 1.60E-03 | - | - |
| renal protein absorption | 2.00E-03 | - | - |
| immunoglobulin production | 2.20E-03 | - | - |
| negative regulation of cell death | 2.20E-03 | - | - |
| positive regulation of ERK1 and ERK2 cascade | 2.30E-03 | - | - |
| positive regulation of apoptotic process | 2.40E-03 | - | - |
| positive regulation of peptidyl-tyrosine phosphorylation | 2.50E-03 | - | - |
| regulation of peptidyl-tyrosine phosphorylation | 2.70E-03 | - | - |
| cell-cell adhesion via plasma-membrane adhesion molecules | 2.90E-03 | - | - |
| defense response to Gram-negative bacterium | 3.10E-03 | - | - |
| response to mechanical stimulus | 3.50E-03 | - | - |
| membrane fission | 3.60E-03 | - | - |
| Arp2/3 complex-mediated actin nucleation | 3.70E-03 | - | - |
| positive regulation of MAP kinase activity | 3.80E-03 | - | - |
| negative regulation of fibrinolysis | 4.10E-03 | - | - |
| receptor-mediated endocytosis | 4.10E-03 | - | - |
| response to lipopolysaccharide | 4.20E-03 | - | - |
| multicellular organism development | 4.40E-03 | - | - |
| plasma membrane repair | 4.90E-03 | - | - |
| response to cold | 4.90E-03 | - | - |
| canonical Wnt signaling pathway | 5.10E-03 | - | - |
| ossification | 5.40E-03 | - | - |
| cell-cell adhesion mediated by cadherin | 5.60E-03 | - | - |
| protein autoprocessing | 5.60E-03 | - | - |
| cell migration | 5.70E-03 | - | - |
| membrane to membrane docking | 6.40E-03 | - | - |
| regulation of organelle assembly | 6.40E-03 | - | - |
| positive regulation of tumor necrosis factor production | 6.40E-03 | - | - |
| defense response to Gram-positive bacterium | 6.90E-03 | - | - |
| protein homotrimerization | 7.60E-03 | - | - |
| maintenance of gastrointestinal epithelium | 7.60E-03 | - | - |
| positive regulation of T cell proliferation | 7.80E-03 | - | - |
| excretion | 8.10E-03 | - | - |
| platelet aggregation | 8.60E-03 | - | - |
| transmembrane receptor protein tyrosine kinase signaling pathway | 8.70E-03 | - | - |
| macroautophagy | 9.00E-03 | - | - |
| leukocyte cell-cell adhesion | 9.10E-03 | - | - |
| negative regulation of cell migration | 9.10E-03 | - | - |
| response to peptide hormone | 9.40E-03 | - | - |
| extracellular matrix organization | 9.50E-03 | - | - |
| animal organ morphogenesis | 1.00E-02 | - | - |
| positive regulation of kinase activity | 1.00E-02 | - | - |
| negative regulation of cell-substrate adhesion | 1.10E-02 | - | - |
| cell surface receptor signaling pathway | 1.10E-02 | - | - |
| positive regulation of neutrophil extravasation | 1.30E-02 | - | - |
| negative regulation of macrophage chemotaxis | 1.30E-02 | - | - |
| response to bacterium | 1.30E-02 | - | - |
| response to starvation | 1.50E-02 | - | - |
| actin filament bundle assembly | 1.50E-02 | - | - |
| triglyceride metabolic process | 1.50E-02 | - | - |
| positive regulation of I-kappaB kinase/NF-kappaB signaling | 1.50E-02 | - | - |
| negative regulation of peptidase activity | 1.60E-02 | - | - |
| response to glucocorticoid | 1.60E-02 | - | - |
| carbohydrate metabolic process | 1.60E-02 | - | - |
| actin polymerization or depolymerization | 1.60E-02 | - | - |
| establishment of endothelial barrier | 1.60E-02 | - | - |
| myoblast differentiation | 1.60E-02 | - | - |
| regulation of centrosome duplication | 1.60E-02 | - | - |
| glycosaminoglycan catabolic process | 1.60E-02 | - | - |
| positive regulation of interferon-gamma production | 1.70E-02 | - | - |
| lung-associated mesenchyme development | 1.70E-02 | - | - |
| fibroblast growth factor receptor signaling pathway | 1.70E-02 | - | - |
| acute-phase response | 1.80E-02 | - | - |
| protein transport | 1.80E-02 | - | - |
| T cell costimulation | 2.00E-02 | - | - |
| actin filament polymerization | 2.00E-02 | - | - |
| cellular iron ion homeostasis | 2.10E-02 | - | - |
| regulation of synapse organization | 2.10E-02 | - | - |
| barbed-end actin filament capping | 2.10E-02 | - | - |
| positive regulation of Wnt signaling pathway | 2.10E-02 | - | - |
| positive regulation of early endosome to late endosome transport | 2.10E-02 | - | - |
| cyclooxygenase pathway | 2.10E-02 | - | - |
| negative regulation of complement activation, classical pathway | 2.10E-02 | - | - |
| protein folding | 2.20E-02 | - | - |
| epidermis development | 2.30E-02 | - | - |
| positive regulation of cell proliferation | 2.60E-02 | - | - |
| protein localization to plasma membrane | 2.60E-02 | - | - |
| multicellular organismal iron ion homeostasis | 2.60E-02 | - | - |
| cellular response to platelet-derived growth factor stimulus | 2.60E-02 | - | - |
| response to food | 2.60E-02 | - | - |
| positive regulation of establishment of protein localization to telomere | 2.60E-02 | - | - |
| branching involved in salivary gland morphogenesis | 2.60E-02 | - | - |
| morphogenesis of an epithelial sheet | 2.60E-02 | - | - |
| nucleoside monophosphate phosphorylation | 2.60E-02 | - | - |
| positive regulation of protein localization to early endosome | 2.60E-02 | - | - |
| interleukin-1-mediated signaling pathway | 2.90E-02 | - | - |
| negative regulation of cell proliferation | 3.00E-02 | - | - |
| multivesicular body-lysosome fusion | 3.20E-02 | - | - |
| vesicle fusion with vacuole | 3.20E-02 | - | - |
| positive regulation of protein localization to Cajal body | 3.20E-02 | - | - |
| hydrogen peroxide catabolic process | 3.20E-02 | - | - |
| response to xenobiotic stimulus | 3.30E-02 | - | - |
| negative regulation of interleukin-2 production | 3.50E-02 | - | - |
| leukocyte migration | 3.50E-02 | - | - |
| cellular response to retinoic acid | 3.50E-02 | - | - |
| protein stabilization | 3.50E-02 | - | - |
| intracellular transport | 3.60E-02 | - | - |
| wound healing | 3.60E-02 | - | - |
| regulation of blood coagulation | 3.70E-02 | - | - |
| regulation of complement activation | 3.70E-02 | - | - |
| regulation of cell-cell adhesion | 3.70E-02 | - | - |
| cell surface pattern recognition receptor signaling pathway | 3.70E-02 | - | - |
| negative regulation of cytokine-mediated signaling pathway | 3.70E-02 | - | - |
| response to electrical stimulus | 3.90E-02 | - | - |
| nucleus organization | 3.90E-02 | - | - |
| cellular response to reactive oxygen species | 4.00E-02 | - | - |
| positive regulation of cell migration | 4.10E-02 | - | - |
| receptor internalization | 4.30E-02 | - | - |
| epidermal growth factor receptor signaling pathway | 4.30E-02 | - | - |
| ephrin receptor signaling pathway | 4.30E-02 | - | - |
| cellular response to interleukin-7 | 4.30E-02 | - | - |
| actin filament severing | 4.30E-02 | - | - |
| positive regulation of integrin-mediated signaling pathway | 4.30E-02 | - | - |
| actin nucleation | 4.30E-02 | - | - |
| regulation of gene expression | 4.50E-02 | - | - |
| cellular response to mechanical stimulus | 4.80E-02 | - | - |
| negative regulation of angiogenesis | 4.80E-02 | - | - |
| osteoblast differentiation | 4.80E-02 | - | - |
| epithelial cell differentiation | 4.80E-02 | - | - |
| negative regulation of Wnt signaling pathway | 4.80E-02 | - | - |
| autophagy | 4.90E-02 | - | - |

表S8 抑郁组单例样本与健康组相比12例及以上样本共有的差异蛋白

| **Accession** | **Protein names** | **Trend** |
| --- | --- | --- |
| Q96IQ7 | V-set and immunoglobulin domain-containing protein 2 | 13↑ |
| P0DOX5 | Immunoglobulin gamma-1 heavy chain | 13↓ |
| A0A075B6S2 | Immunoglobulin kappa variable 2D-29 | 13↓ |
| P01833 | Polymeric immunoglobulin receptor | 13↓ |
| O00187 | Mannan-binding lectin serine protease 2 | 13↓ |
| P50591 | Tumor necrosis factor ligand superfamily member 10 | 13↓ |
| P01859 | Immunoglobulin heavy constant gamma 2 | 13↓ |
| P01834 | Immunoglobulin kappa constant | 13↓ |
| P10153 | Non-secretory ribonuclease | 13↓ |
| P10451 | Osteopontin | 13↓ |
| C9IZ46 | Protein shisa-5 | 13↓ |
| O75339 | Cartilage intermediate layer protein 1 | 13↓ |
| Q96FE7 | Phosphoinositide-3-kinase-interacting protein 1 | 13↓ |
| P07998 | Ribonuclease pancreatic | 13↓ |
| P01009 | Alpha-1-antitrypsin | 13↓ |
| P01042 | Kininogen-1 | 13↓ |
| P10253 | Lysosomal alpha-glucosidase | 2↑11↓ |
| P15328 | Folate receptor alpha | 2↑11↓ |
| P00734 | Prothrombin | 2↑11↓ |
| Q8WZ75 | Roundabout homolog 4 | 2↑11↓ |
| B8ZZQ6 | Prothymosin alpha | 2↑11↓ |
| A0A087X0K0 | Collagen type XV alpha 1 chain | 2↑11↓ |
| P02790 | Hemopexin | 2↑11↓ |
| P02790 | Hemopexin | 2↑11↓ |
| A0A2R8Y478 | CD9 molecule | 7↑6↓ |
| Q13740 | CD166 antigen | 12↑ |
| Q86SF2 | N-acetylgalactosaminyltransferase 7 | 12↑ |
| Q9Y5F6 | Protocadherin gamma-C5 | 12↑ |
| I3L4I4 | Target of myb1 like 1 membrane trafficking protein | 12↑ |
| Q8IWU5 | Extracellular sulfatase Sulf-2 | 12↓ |
| P0DP57 | Secreted Ly-6/uPAR domain-containing protein 2 | 12↓ |
| Q6EMK4 | Vasorin | 12↓ |
| P07602 | Prosaposin | 12↓ |
| A0A0G2JLV7 | Leukocyte-associated immunoglobulin-like receptor 1 | 12↓ |
| Q16651 | Prostasin | 12↓ |
| P02760 | Protein AMBP | 12↓ |
| P12109 | Collagen alpha-1(VI) chain | 12↓ |
| O75594 | Peptidoglycan recognition protein 1 | 12↓ |
| P05090 | Apolipoprotein D | 12↓ |
| G3V4U0 | Fibulin 5 | 12↓ |
| P01876 | Immunoglobulin heavy constant alpha 1 | 12↓ |
| J3QQX6 | Intercellular adhesion molecule 2 | 12↓ |
| D6RBV2 | Lectin, mannose binding 2 | 12↓ |
| P02768 | Albumin | 12↓ |
| O60494 | Cubilin | 1↑11↓ |
| Q6UXB8 | Peptidase inhibitor 16 | 1↑11↓ |
| P01034 | Cystatin-C | 1↑11↓ |
| P60022 | Beta-defensin 1 | 1↑11↓ |
| Q9NZP8 | Complement C1r subcomponent-like protein | 1↑11↓ |
| P61970 | Nuclear transport factor 2 | 1↑11↓ |
| P0DOX8 | Immunoglobulin lambda-1 light chain | 1↑11↓ |
| P61769 | Beta-2-microglobulin | 1↑11↓ |
| P05060 | Secretogranin-1 | 1↑11↓ |
| P04745 | Alpha-amylase 1A | 1↑11↓ |
| A0M8Q6 | Immunoglobulin lambda constant 7 | 1↑11↓ |
| Q9HCU0 | Endosialin | 2↑10↓ |
| P01624 | Immunoglobulin kappa variable 3-15 | 2↑10↓ |
| P08294 | Extracellular superoxide dismutase [Cu-Zn] | 3↑9↓ |
| Q7LBR1 | Charged multivesicular body protein 1b | 3↑9↓ |
| Q9NPF0 | CD320 antigen | 3↑9↓ |
| Q13201 | Multimerin-1 | 4↑8↓ |
| O00560 | Syntenin-1 | 4↑8↓ |
| Q99784 | Neuronal olfactomedin-related ER localized protein | 8↑4↓ |
| P11717 | Cation-independent mannose-6-phosphate receptor | 5↑7↓ |
| A0A0C4DH35 | Probable non-functional immunoglobulin heavy variable 3-35 | 7↑5↓ |
| Q7Z7M0 | Multiple epidermal growth factor-like domains protein 8 | 6↑6↓ |
